# CHO/LY-B cell growth under limiting sphingolipid supply: correlation between lipid composition and biophysical properties of sphingolipid-restricted cell membranes^+^

**DOI:** 10.1101/2020.07.05.188474

**Authors:** Bingen G. Monasterio, Noemi Jiménez-Rojo, Aritz B. García-Arribas, Howard Riezman, Félix M. Goñi, Alicia Alonso

**Affiliations:** Instituto Biofisika (CSIC, UPV/EHU), Universidad del País Vasco, 48940 Leioa, Spain; Departamento de Bioquímica, Universidad del País Vasco, 48940 Leioa, Spain; NCCR Chemical Biology, Department of Biochemistry, University of Geneva, 1211 Geneva, Switzerland

**Keywords:** CHO, LY-B, laurdan, membrane fluidity, AFM, plasma membrane, lipidomics, mass-spectroscopy, sphingolipids, sphingomyelin

## Abstract

Sphingolipids (SL) are ubiquitous in mammalian cell membranes, yet there is little data on the behavior of cells under SL-restriction conditions. LY-B cells derive from a CHO line in which serine palmitoyl transferase (SPT), thus *de novo* SL synthesis, is suppressed, while maintaining the capacity of taking up and metabolizing exogenous sphingoid bases from the culture medium. In the present study LY-B cells were adapted to grow in a fetal bovine serum (FBS)-deficient medium to avoid external uptake of lipids. The lowest FBS concentration that allowed LY-B cell growth, though at a slow rate, under our conditions was 0.04%, i.e. 250-fold less than the standard (10%) concentration. Cells grown under limiting SL concentrations remained viable for at least 72 h. Enriching with sphingomyelin the SL-deficient medium allowed the recovery of control LY-B cell growth rates. Studies including whole cells, plasma membrane preparations, and derived lipid vesicles were carried out. Laurdan fluorescence was recorded to measure membrane molecular order, showing a significant decrease in the rigidity of LY-B cells, not only in plasma membrane but also in whole cell lipid extract, as a result of SL limitation in the growth medium. Plasma membrane preparations and whole cell lipid extracts were also studied using atomic force microscopy in the force spectroscopy mode. Force measurements demonstrated that lower breakthrough forces were required to penetrate samples obtained from SL-poor LY-B cells than those obtained from control cells. Mass-spectroscopic analysis was also a helpful tool to understand the rearrangement undergone by the LY-B cell lipid metabolism. The most abundant SL in LY-B cells, sphingomyelin, decreased by about 85% as a result of SL limitation in the medium, the bioactive lipid ceramide and the ganglioside precursor hexosylceramide decreased similarly, together with cholesterol. Quantitative SL analysis showed that a 250-fold reduction in sphingolipid supply to LY-B cells led to a 6-fold decrease in membrane sphingolipids, underlining the resistance to changes in composition of these cells. Plasma membrane compositions exhibited similar changes, at least qualitatively, as the whole cells with SL restriction. A linear correlation was observed between the sphingomyelin concentration in the membranes, the degree of lipid order as measured by laurdan fluorescence, and membrane breakthrough forces assessed by atomic force microscopy. Concomitant changes were detected in glycerophospholipids under SL-restriction conditions.

## Introduction

Sphingolipids (SL) are characterized by a sphingoid structural backbone, sphingosine being the most abundant base in mammals^1^. *In vivo* studies of the SL roles are hampered, among other reasons, by the fact that they can be either synthesized de *novo* or taken up from the diet. A plausible approach would be to investigate mutant cells containing the smallest possible amounts of SL, or even none at all, but this is not a straightforward procedure because SL appear to be essential for cell growth and survival^2–4^. In particular, SL are considered as instrumental in the architecture of eukaryotic cell membranes. Apart from stabilizing the lamellar structure and helping to maintain its asymmetry, the tendency of certain SL to undergo lateral phase separation to form micro or nanodomains has been characterized^5–8^. Lipids and proteins co-localize with these domains, in which sphingomyelin (SM) is the most abundant SL^9^, and cholesterol (Chol) is often present^7,8^. The sphingoid base provides SL with hydrogen-bonding acceptors and donors (amide and free hydroxyl groups, respectively) that are rare in glycerophospholipids, leading to a dense intermolecular hydrogen-bonding network^10,11^. Hydrogen bonding allows SM to interact preferentially with Chol^12,13^ and with ceramide (Cer)^6^, and recent studies have found that sphingomyelin (SM), Cer and Chol are able to coexist in a single ternary gel phase (at a 54:23:23 mol ratio) with intermediate properties between SM-Cer enriched gel domains and Chol-driven liquid-ordered phases^14^. The above data have been obtained mostly from model membrane studies. Investigations at the cellular level have allowed to assign a wide variety of functions to SL, including apoptosis, cell growth, cell membrane function, tumor formation, drug resistance, degranulation, and phagocytosis, among others^15–18^. Alterations in the normal activities of SM-cycle enzymes have been linked to many central nervous system-related pathologies such as Alzheimer’s, Parkinson’s, ischemia/hypoxia, depression, schizophrenia or Niemann-Pick diseases^19^.Specific SL functions have often been characterized in cells with decreased amounts of SL, using either SL-degrading enzymes (e.g. sphingomyelinases or ceramidases)^20^, or specific enzyme inhibitors^15,21–23^, of which the SPT inhibitor myriocin is a good example^24–27^.

Procedures to decrease the SL contents of cells constitute good tools to determine their potential functions *in vivo*. *De novo* SL biosynthesis is initiated by the condensation of L-serine with palmitoyl CoA. This reaction is catalyzed by SPT to generate 3-ketodihydrosphingosine. 3-Ketodihydrosphingosine is then converted to dihydrosphingosine, which is N-acylated and (most of it) dehydrogenated at the endoplasmic reticulum to form Cer. After moving to the Golgi apparatus, Cer is converted to SM or glycosphingolipids^6,28^. Finally, these complex SL are translocated to the PM^29,30^. Thus, SPT is a key enzyme for the regulation of cellular SL content^9,29^. Using a genetic selection method in CHO cells^31^, the Hanada lab isolated the defective LY-B cell line, which had a loss of function of serine palmitoyltransferase (SPT) enzyme activity through a defective SPTLC1 subunit. The mutant cells maintained the ability to take up and metabolize exogenous sphingoid bases from the culture medium^31^. Mutant LY-B and wild type CHO cells could be comparatively studied to determine the effect of SL depletion on the biophysical properties of cell membranes. LY-B cells have been used in multiple studies exploring SL effects, and their interaction with glycerophospholipid metabolism^32–35^. SM synthases, which use Cer and phosphatidylcholine as substrates to produce SM and diacylglyceride, are used in the *de novo* synthesis pathway, sometimes also involved in reutilization of ceramide^30,36^ from exogenous or endogenous sources. In a previous work^37^ plasma membrane (PM) preparations from CHO cells and model membranes prepared from their lipid extracts had been characterized. In the present study we have applied those methods to the study of SL-synthesis deficient LY-B cells to determine the role of SL on cell growth, membrane physical properties and composition.

## Materials and methods

### Cell growth

Wild type CHO (ATCC, Manassas, Virginia, U.S.) and a serine-SPT deficient CHO cell line, known as LY-B^31^ (RIKEN BioResource Research Center, Koyadai, Japan), were used in this study. Unless otherwise mentioned, cells were grown on DMEM:F12 (Dulbecco’s Modified Eagle Medium: Nutrient Mixture F-12) medium containing 10% FBS (Fetal Bovine Serum), 100 U/ml penicillin, 100 U/ml streptomycin, and 6 mM glutamine (GlutaMax supplemented) at 37°C and 5% CO_2_ humidified atmosphere. All cell culture products were purchased from Thermofisher (Waltham, MA).

#### Cell adaptation: Standard vs. deficient (low-FBS) medium

CHO and LY-B cells were adapted to growth in deficient (low-FBS) medium. For this purpose, cells were first seeded in DMEM:F12 medium containing 10% FBS, 100 U/ml penicillin and 100 U/ml streptomycin, and 6 mM glutamine (this medium will be referred to as ‘standard medium’). After 24-h cell growth, when a 15-25% confluence was reached, the standard medium was discarded, cells were washed with PBS buffer (137 mM NaCl, 3 mM KCl, 80 mM Na_2_HPO_4,_ 7 mM KH_2_PO_4_), and DMEM:F12 medium containing 0.04% FBS, 100 U/ml penicillin and 100 U/ml streptomycin and 6 mM Glutamine was added (this medium will be named ‘FBS-deficient’ or ‘SL-deficient medium’). Cells were grown in the appropriate medium for 24, 48, 72 or 96 h. Other low-FBS media (5%, 2.5% or 1.25%) were also used in some specific cases.

### Growth rate and viability tests

#### Cell growth

2.65*10^5^ cells were seeded in 25 cm^2^ flasks in standard medium and grown for 24 h until 15 – 25% confluence. Then, the standard medium was discarded, cells were washed twice with PBS, and the appropriate medium (standard or deficient) was added. Cells were grown for 24, 48, 72 or 96 h. Quantification was performed by cell counting with a hemocytometer (BioRad TC20 Automated Cell Counter, Hercules, CA). Protein was assayed with the colorimetric Pierce BCA Protein Assay Kit (Thermofisher, Waltham, MA).

#### Viability test

Fluorescence-activated cell sorting (FACS) was performed to evaluate how the decreased FBS concentration in the medium affected cell viability^38^. Cells were stained with Annexin-V-FITC and propidium iodide as indicated in the manual of the annexin V-FITC detection kit (CalbioChem, Darmstadt, Germany) and fluorescence was measured using a FACS Calibur flow cytometer (Becton-Dickinson, Franklin Lakes, NJ) as in Ahyayauch *et al.*^39^. Annexin V-FITC fluorescence intensity was measured in fluorescence channel FL-1 with λ_ex_ = 488 nm and λ_em_ = 530 nm, while FL-3 was used for propidium iodide detection, with λ_ex_ = 532 nm and λ_em_ = 561 nm. All measurements were performed in triplicate. Data analysis was performed using Flowing Software 2.

### Sample preparation

Intact cells (whole cells), two different PM preparations (giant plasma membrane vesicles, known as GPMV or blebs, and PM patches) and SUV or GUV formed with whole-cell or PM lipid extracts were used.

#### PM preparations

PM preparations were obtained as described in Monasterio *et al.*^37^. Briefly, GPMV formation was induced adding the GPMV formation reagent [freshly prepared 2 mM dithiothreitol, 25 mM paraformaldehyde in GPMV buffer (2 mM CaCl_2_, 10 mM HEPES, 150 mM NaCl, pH 7.4)] to T25 flasks with cells at confluence. Cells were incubated for 1 h at 37 °C. After incubation, the GPMV-containing GPMV reagent was collected from the flasks and centrifuged at 14,000g for 20 min. Supernatant was discarded and several washing and centrifugation steps were conducted to remove traces of dithiothreitol and paraformaldehyde. Finally, the GPMV were resuspended in 500 μl GMPV buffer^40^.

PM patches were isolated by a modification^37^ of the protocol described by Bezrukov *et al.*^42^. In summary, cells were seeded at approximately 50% confluence and incubated for 2 h so that they adhered to the support. After incubation, 2 washing steps were performed using cold TBS (Tris Buffer Saline: 150 mM NaCl, 25 mM Tris-HCl, 2 mM KCl) to discard non-attached cells. Then, cold distilled water was added for 2 min to induce cell swelling. Mechanical cell disruption was achieved using a pressure stream from a 20-ml syringe coupled to a 19X1-1/2(TW)A needle. In the process, intracellular contents were released, while PM stayed attached to the support. Several washing steps were performed to discard the released intracellular contents. Purification quality was checked using Di-4-ANEPPDHQ (λ_ex_ = 465 nm, λ_em_ = 635 nm) as a general fluorescent staining, together with organelle-specific fluorophores as described in Monasterio *et al.*^37^. Images were taken in a Leica TCS SP5 II microscope (Leica Microsystems GmbH, Wetzlar, Germany) at room temperature with ImageJ software. The fluorescence intensities of the various markers were comparatively measured in PM patches and intact cells, so that specific organelle contamination could be estimated.

#### Whole cell lipid extract

Lipid extraction was performed following the method used in Ahyayauch *et al.*^39^. Briefly, cell pellets were first dispersed in aqueous perchloric acid (60% v/v), then centrifuged at 14,000g for 15 min, and the supernatant was discarded. Pellets were resuspended in 2.5 ml chloroform:methanol (2:1, v/v) and samples were mixed for 15 min. Then, 5 ml cold 0.1 mM HCl was added to the mixture. After homogenizing, samples were centrifuged at 1,700g for 20 min. Supernatants were discarded while the lipid-containing organic phase remained in the bottom layer. Phospholipid concentration was assayed as inorganic phosphorus after acid digestion.

#### PM patch lipid extraction

PM patches were formed following the above mentioned protocol^37,42^. Chloroform:methanol (2:1) organic solvent was used to recover the lipid fraction of the attached PM patches.

#### GUV formation

GUV were formed in a PRETGUV 4 chamber supplied by Industrias Técnicas ITC (Bilbao, Spain) using the modified electroformation method^43^ first developed by Angelova and Dimitrov^44^.

#### SUV formation

The sample was kept under vacuum for 2 h to remove solvent traces and the lipids were swollen in PBS buffer. SUV were obtained by sonication of the swollen lipid suspensions with a probe-type Soniprep 150 sonicator (MSK, London, U.K.) for 10 min, in 10-s on, 10-s off intervals.

### SM quantification with lysenin

#### Lysenin-mCherry expression and purification

The non-toxic monomeric C-terminal domain of the SM-specific toxin, NT-lysenin, was expressed and purified as described by Carquin *et al.*^45^. Briefly, the expression plasmid pET28/lysenin encoded NT-lysenin as a fusion protein with an N-terminal 6xHis-tag followed by the monomeric red fluorescent protein mCherry. The plasmid was expanded in *Escherichia coli* BL21 (DE3) and the recombinant protein was expressed in lysogeny broth (LB) medium at 16°C for 72 h in the presence of 0.4 mM isopropyl β-D-thiogalactoside. Bacterial extracts were prepared as described^46^ and the recombinant protein was purified using an Ni-NTA Superflow cartridge (Qiagen, Hilden, Germany) and eluted with imidazol^47^. Fraction analysis by SDS-PAGE revealed recombinant NT-lysenin with the expected size (45 kDa). The most enriched fractions were pooled, concentrated, and desalted. The aliquots were stored in 20 mM NaCl and 25 mM Hepes pH 7.2 and 5% glycerol at −80°C. Protein concentration was calculated by measuring absorbance at 280 nm.

#### SM staining and quantification with lysenin-mCherry

Whole cells and PM patches containing SM were stained with lysenin-mCherry and samples were then visualized using a confocal microscopy Nikon D-ECLIPSE C1 (Nikon, Melville, NY). In whole cells the mCherry signal was also quantified using a FL-3 FACS Calibur flow cytometer (Becton-Dickinson, Franklin Lakes, NJ) with λ_ex_ = 532 nm and λ_em_ = 561 nm.

For sample visualization, cells were seeded in glass-bottom dishes and grown as above. Cells were first stained with 100 μM NBD-PE as a control for all membrane staining. A washing step was performed with PBS, and lysenin-mCherry was added at 100 μM. For FACS analysis, cells were stained in suspension at a final concentration of 100 μM lysenin-mCherry.

### Laurdan General Polarization

Laurdan is a fluorescence polarity probe whose emission undergoes a spectral shift due to the reorientation of water molecules in the glycerol backbone region of the membrane, and this shift can be correlated to the lipid phase^48^. In the gel phase, when little water is present, laurdan maximum emission is around 440 nm, whereas in the liquid crystalline phase the spectrum is red shifted to around 490 nm. Intact cells, PM preparations and model membranes formed with lipid extracts have been used to compare the laurdan fluorescence of CHO and LY-B cells grown in standard and deficient media. Fluorescence microscopy imaging and spectrofluorometric analysis have been performed for laurdan fluorescence characterization.

#### Confocal microscopy

For intact cells, cells grown in glass-bottom dishes in FBS-containing medium were stained with 5 μM laurdan (Molecular Probes, Eugene, OR) for 5 min. Several washes were performed with PBS before cells were visualized. GPMV were stained with 5 μM laurdan and transferred to polylysine-coated glass-bottom dishes (MatTek, Ashland, OR). Vesicles were left to sediment for 3 h before visualization^40^. For GUV, 0.2 mM lipid extracts in chloroform:methanol (2:1, v/v) were mixed with 0.01 mM laurdan. 3 μl of the lipid stocks were added onto the surface of Pt electrodes and solvent traces were removed under high vacuum for at least 2 h. The Pt electrodes were then covered with 400 μl of 300 mM sucrose buffer and the Pt wires were connected to an electric wave generator (TG330 function generator, Thurlby Thandar Instruments, Huntington, United Kingdom) under alternating current field conditions (10 Hz, 2.5 VRMS for 2 h) at 37°C. After GUV formation, the chamber was placed on an inverted confocal fluorescence microscope for GUV visualization.

#### Image acquisition and analysis

A Leica TCS SP5 II microscope (Leica Microsystems GmbH, Wetzlar, Germany) was used for image acquisition. A 63x water-immersion objective (numerical aperture NA = 1.2) was used and samples were imaged at 512 x 512 pixel and 400 Hz per scanning line. Equatorial planes were imaged to avoid photoselection effects. A pulsed titanium-sapphire (Mai-Tai Deepsee, Spectra-Physics) laser tuned at 780 nm was used for two-photon imaging of laurdan-labeled samples. Fluorescence emission was collected by non-descanned (NDD) hybrid detectors, as they offer higher sensitivity compared to descanned photomultipliers. The blue edge of the emission spectrum was collected by NDD 1 at 435 ± 20 nm and the red edge by NDD 2 at 500 ± 10 nm. Irradiance at the sample plane was ≈500 GW·cm^−2^ for two-photon excitation^49^.

Generalized polarization (GP) value of samples was calculated using a MATLAB (MathWorks, Natick, MA) based software. Images were smooth in each channel with 2 pixel averaging, and the GP value was calculated using the following equation^50^:

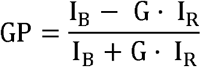

where I_B_ is the intensity collected by NDD 1, I_R_ is the intensity collected by NDD 2, and G is the correction factor. The G factor is calculated measuring the GP value of the same fluorophore concentration used in sample staining, dissolved in this case in pure DMSO^51^. In whole cell images, the region of interest, i.e. the PM, was selected when required.

#### Fluorescence spectroscopic analysis

PM preparations, and SUV formed with whole cell and PM lipid extracts were measured in a spectrofluorometer. Samples (82.5 μM lipid concentration) were labeled with 0.75 μM laurdan. For this purpose lipid extracts in chloroform:methanol (2:1) were mixed with laurdan and the solvent was evaporated to dryness under a stream of N_2_. Then, the sample was kept under vacuum for 2 h to remove solvent traces and the lipids were swollen in buffer (NaCl 150 mM, Hepes 25 mM, pH 7.4). Sonicated SUV were obtained as described above and fluorescence measurements were performed using a QuantaMaster 40 spectrofluorometer (Photon Technology International, Lawrenceville, NJ)^52^.

### AFM

Contact mode AFM imaging has been used to study bilayer topography, looking at possible lateral segregation effects through bilayer thickness analysis. A NanoWizard II AFM (JPK Instruments, Berlin, Germany) was used to perform topographic measurements under contact mode scanning (constant vertical deflection). For measurements, the AFM was coupled to a Leica microscope and mounted onto a Halcyonics Micro 40 antivibration table (Halcyonics, Inc., Menlo Park, CA) and inside an acoustic enclosure (JPK Instruments, Berlin, Germany)^53^. V-shaped MLCT Si_3_N_4_ cantilevers (Bruker, Billerica, MA) with nominal spring constants of 0.1 or 0.5 N/m. The sample thickness was estimated by cross-section height analysis^54^.

#### Force Spectroscopy

V-shaped MLCT Si_3_N_4_ cantilevers (Bruker, Billerica, MA) with nominal spring constants of 0.1 or 0.5 N/m were individually calibrated in a lipid-free mica substrate in assay buffer using the thermal noise method. After proper bilayer area localization by means of AFM topography and direct epifluorescence microscopy, force spectroscopy was performed at a speed of 1 μm/s. Force steps were determined for each of the indentation curves as reproducible jumps within the extended traces. At least three independent sample preparations were scanned for each case and 50-100 curves were measured in each sample.

Topographic images and force spectroscopy analysis of PM patches and supported planar bilayers (SPB) formed from lipid extracts, and force spectroscopy analysis of GPMV were performed. GPMV topographic observations could not be performed for experimental reasons, these structures would not flatten on the mica for AFM examination.

SPB were prepared on high V-2 quality scratch-free mica substrates (Asheville-Schoonmaker Mica Co., Newport News, VA). 180 μL assay buffer containing 3 mM CaCl_2_ was added onto a 1.2 cm^2^ freshly cleaved mica substrate mounted onto a BioCell (JPK Instruments, Berlin, Germany). Then, 80 μL sonicated 0.4 mM SUV formed with CHO or LY-B lipid extract was added on top of the mica. BioCell temperature was gradually increased (5°C every 5 min) up to 80°C. Vesicles were let to adsorb and extend for 30 min keeping the sample temperature at 80°C. Samples were left to equilibrate for 30 min at room temperature before performing five washing steps with non-CaCl_2_ buffer in order to discard non-adsorbed vesicles and remove the remaining Ca^2+^ cations^53^. Isolated PM patches for force spectroscopy were prepared as previously described^37,42^, this time using polylysine-coated mica slips instead of glass-bottom dishes. GPMV were first stained using Di-4-ANEPPQHD to allow detection on the mica slip. Then, samples were left for 3 h to sediment over the polylysine-coated mica slip before measurements were performed.

### Mass spectroscopic analysis

Mass spectroscopic analysis was performed essentially as described in Monasterio *et al*.^37^. A methodological summary follows.

#### Sample treatment

Lipid extraction was performed using a modified methyl *tert*-butyl ether (MTBE) protocol^55^. Briefly, cells were washed with cold PBS and scraped off in 500 μl cold PBS on ice. The suspension was transferred to a 2 ml tube in which it was spun down at 3200 rpm for 5 min at 4 °C. After removing the PBS, samples were stored at −20°C or directly used for further extraction. GPMV and PM patch samples were prepared as previously mentioned. Then, 360 μl methanol was added and vortexed. A mixture of lipid standards (see Table 1) was added and samples were vortexed for 10 min at 4 °C using a Cell Disruptor Genie (Scientific Industries, Inc., Bohemia, NY). MTBE (1.2 mL) was then added and the samples were incubated for 1 h at room temperature with shaking (750 rpm). Phase separation was induced by adding 200 μl H_2_O. After 10 min incubation at room temperature, the sample was centrifuged at 1,000 x g for 10 min. The upper (organic) phase was transferred to a 13-mm screw-cap glass tube and the lower phase was extracted with 400 μl artificial upper phase (MTBE/methanol/water (10:3:1.5, v/v/v)). The two upper phases were combined and the total lipid extract was divided in 3 equal aliquots (one for phospholipids (TL), one for sterols (S) in 2-mL amber vials, and one for sphingolipid (SL) detection in a 13-mm glass tube) and dried in a Centrivap at 50°C or under a nitrogen flow. The SL aliquot was deacylated by methylamine treatment (Clarke method) to remove glycerophospholipids. 0.5 mL monomethylamine reagent [MeOH/H_2_O/n-butanol/methylamine solution (4:3:1:5 v/v)] was added to the dried lipid, followed by sonication (5 min). Samples were then mixed and incubated for 1 h at 53°C and dried (as above). The monomethylamine-treated lipids were desalted by n-butanol extraction. 300 μl H_2_O-saturated n-butanol was added to the dried lipids. The sample was vortexed, sonicated for 5 min and 150 μl MS-grade water was added. The mixture was vortexed thoroughly and centrifuged at 3200 x g for 10 min. The upper phase was transferred to a 2-mL amber vial. The lower phase was extracted twice more with 300 μl H_2_O-saturated n-butanol and the upper phases were combined and dried (as above).

**Table 1.** MS Detection Conditions for the Different Lipid Classes.

| **Lipid Class** | **Standard** | **Polarity** | **Mode** | **m/z ion** | **Collision Energy** |
| --- | --- | --- | --- | --- | --- |
| Phosphatidylcholine [M+H]<sup>+</sup> | DLPC | + | Product ion | 184.07 | 30 |
| Phosphatidylethanolamine [M+H]<sup>+</sup> | PE31:1 | + | Neutral ion loss | 141.02 | 20 |
| Phosphatidylinositol [M-H]<sup>-</sup> | PI31:1 | - | Product ion | 241.01 | 44 |
| Phosphatidylserine [M-H]<sup>-</sup> | PS31:1 | - | Neutral ion loss | 87.03 | 23 |
| Cardiolipin [M-2H]<sup>2-</sup> | CL56:0 | - | Product ion | acyl chain | 32 |
| Ceramide [M+H]<sup>+</sup> | C17Cer | + | Product ion | 264.34 | 25 |
| Dihydroceramide [M+H]<sup>+</sup> | C17Cer | + | Product ion | 266.40 | 25 |
| Hexosylceramide [M+H]<sup>+</sup> | C8GC | + | Product ion | 264.34 | 30 |
| Hexosyldihydroceramide [M+H]<sup>+</sup> | C8GC | + | Product ion | 266.40 | 30 |
| Sphingomyelin [M+H]<sup>+</sup> | C12SM | + | Product ion | 184.07 | 26 |

#### Glycerophospholipid and sphingolipid detection on a Triple Quadrupole Mass Spectrometer

TL and SL aliquots were resuspended in 250 μl chloroform/methanol (1:1 v/v) (LC-MS/HPLC grade) and sonicated for 5 min. The samples were pipetted in a 96-well plate (final volume = 100 μl). The TL were diluted 1:4 in negative-mode solvent (chloroform/methanol (1:2) + 5 mM ammonium acetate) and 1:10 in positive-mode solvent (chloroform/methanol/water (2:7:1 v/v) + 5 mM ammonium acetate). The SL were diluted 1:10 in positive-mode solvent and infused onto the mass spectrometer. Tandem mass spectrometry for the identification and quantification of sphingolipid molecular species was performed using Multiple Reaction Monitoring (MRM) with a TSQ Vantage Triple Stage Quadrupole Mass Spectrometer (Thermofisher Scientific, Waltham, MA) equipped with a robotic nanoflow ion source, Nanomate HD (Advion Biosciences, Ithaca, NY). The collision energy was optimized for each lipid class. The detection conditions for each lipid class are listed below (Table 1). Cer species were also quantified with a loss of water in the first quadrupole. Each biological replica was read in 2 technical replicas (TR). Each TR comprised 3 measurements for each transition. Lipid concentrations were calculated relative to the relevant internal standards and then normalized to the total lipid content of each lipid extract (mol %).

### Gas chromatography–mass spectrometry for cholesterol assay

Lipid extracts were analyzed by GC-MS as described previously^56^. Briefly, samples were injected into a VARIAN CP-3800 gas chromatograph equipped with a FactorFour Capillary Column VF-5ms 15 m × 0.32 mm i.d. DF = 0.10, and analyzed in a Varian 320 MS triple quadrupole with electron energy set to –70 eV at 250°C. Samples were applied to the column oven at 45°C, held for 4 min, then raised to 195°C (20°C/min). Sterols were eluted with a linear gradient from 195 to 230°C (4°C/min), followed by rising to 320°C (10°C/min). Cholesterol was identified by its retention time (compared with an ergosterol standard) and fragmentation patterns, which were compared with the NIST library.

### Quantitation of lipids per cell

An estimate of the amounts of lipids per cell, or per weight protein, was obtained as follows. Kurano *et al*.^57^ measured the average dry weight of CHO cells as 37 ng/cell. Alberts *et al.*^58^ indicated that the mammalian cell contained 10 dry wt% phospholipids and 7 dry wt% other lipids. Moreover, the average amount of protein per cell was measured experimentally with the BCA protein assay. Cell numbers were counted with a hemocytometer. From the above data, and knowing from MS analysis the nanomolar concentration of a given lipid in a sample, the amount of such lipid per cell and per wt protein could be estimated.

## Results

### CHO-derived mutant cells can grow and survive with extremely low sphingolipid concentrations in the culture medium

#### Growth and viability

The extent to which CHO (wild type) and LY-B (SPT-defective) cell adaptation to a SL-deficient medium affected cell division ratio and integrity was assessed. FBS was the only external source of SL for cell growth, thus FBS in the growth medium was the only SL source for LY-B cells. SL-deficient growth media were prepared containing 0.04% FBS, i.e. a 250-fold decrease with respect to the standard conditions (10% FBS). Cell count measurements were performed using a BioRad TC20 hemocytometer. Figure 1A shows a comparison between cell growth in standard (containing 10% FBS) or SL-deficient (containing 0.04% FBS) medium. Both cell lines (CHO and LY-B) grew steadily for at least 96 h in full medium (10% FBS), and both divided, even if slowly, for the first 72 h in a low-SL medium. After 72 h in the SL-limited medium cell quantity decreased. The difference between CHO and LY-B cell-growth ratios was not statistically significant when they were grown in standard medium. Nevertheless, in SL-deficient medium CHO and LY-B cells behaved differently. After 72 h, CHO cell number was 47% of control (complete medium), while LY-B growth was only 18%.

**Figure 1.**
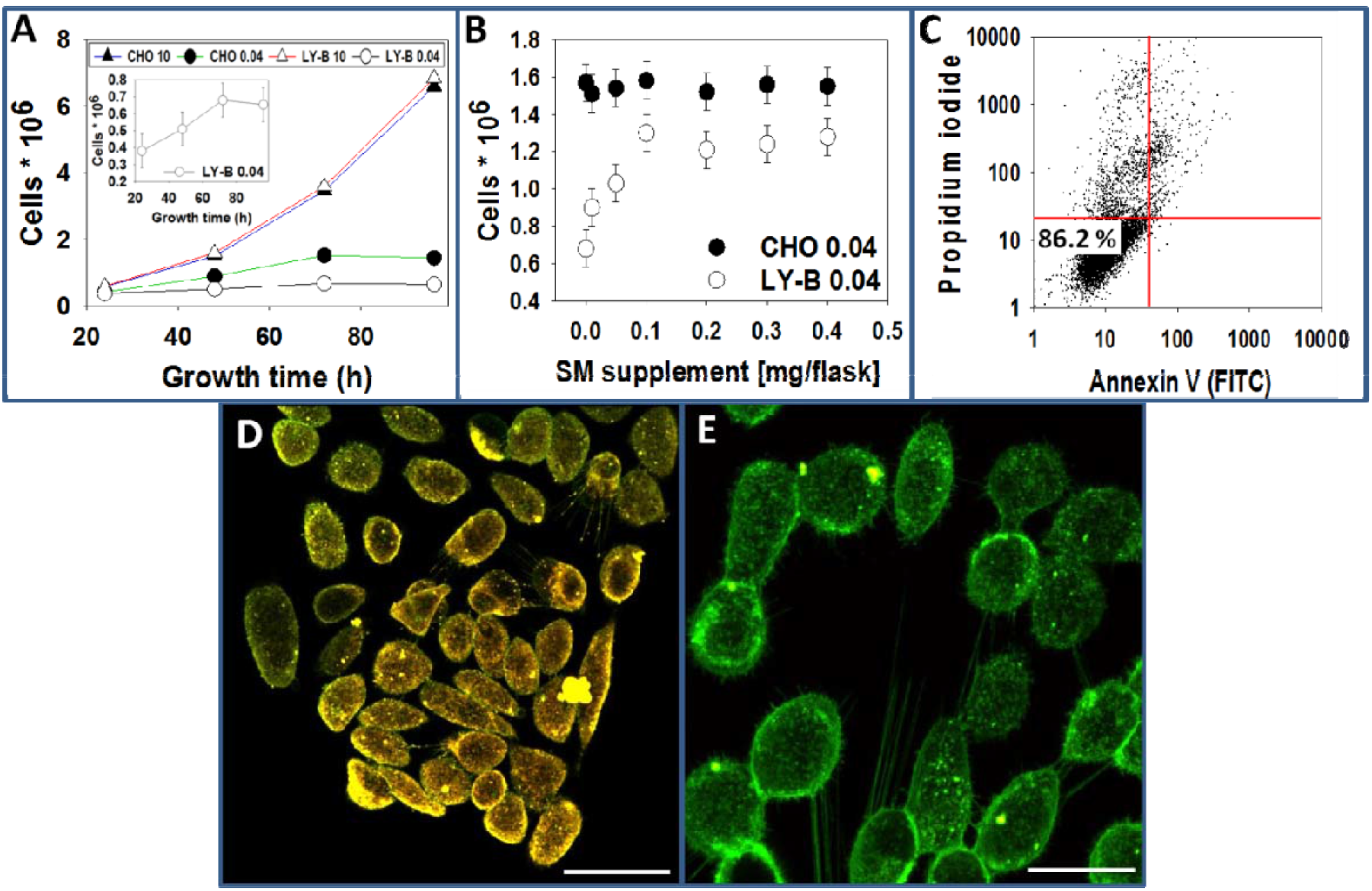
LY-B cells can grow with very small amounts of sphingolipids. LY-B and CHO cell growth as a function of time in standard (10% FBS) and sphingolipid-deficient (0.04% FBS) medium (**A**). LY-B and CHO cell growth after 72 h in sphingolipid-deficient medium supplemented with SM (seeded cells: 0.25 * 10^6^) (**B**). FACS assessment of LY-B cell viability after 72 h in sphingolipid-deficient medium (**C**). Fluorescence images of LY-B cells after 72 h growth in standard (D) and SL-deficient (E) medium; stain: mCherry-lysenin (SM-specific) + NBD-PE (general membrane stain), bar = 10 μm. No lysenin stain is observed in E. For A, B, statistical significance was calculated with ANOVA and Student’s t-test, with similar results: (*) p<0.05; (**) p<0.01; (***) p<0.001 (n = 3).

Ancillary experiments were performed in which cell growth after 72 h in media with different degrees of SL-limitation was measured (Fig. S1). Cell counts were performed as in Fig. 1 (Fig. S1A) and total cell protein was quantified with a BCA protein assay (Fig. S1B). After 72 h, differences in growth were already significant between CHO and LY-B cells when medium contained 5% FBS. No LY-B cell growth could be reliably measured with FBS concentrations below 0.04%.

To ascertain that the poor cell growth at low FBS concentrations was indeed due to a lack of SL, we tested whether LY-B cells were able to reach the full growth rates when the SL-deficient medium was supplemented with SL. For this purpose, equimolar mixtures of egg PC and the sphingolipid under study were sonicated in buffer and added to the culture flasks in various amounts. The best recovery was achieved with sphingomyelin (SM) or sphinganine. Figure 1B shows that LY-B cells grown in SL-deficient medium for 72 h reached ≈80% of the control growth when 0.2 mg SM was added per T25 culture flask (5 ml). With 0.0125 mg sphinganine/flask, cells grown in SL-deficient medium under the same conditions reached 81% of the control (not shown in the figure). It was concluded that it is the lack of SPT activity that makes the main difference between CHO and LY-B cell division ratios in SL-deficient medium.

The time-course of cell growth in SL-deficient medium supplemented with SM can be seen in Figure S1D. LY-B cell growth in deficient medium supplemented with 0.2 mg SM/T25 flask was significantly higher than when no SM was added, while there was no difference in the case of CHO cells (Fig. S1E). All further experiments were performed on cells grown for 72 h.

The possible effect of low cell growth/ SL-deficient medium on cell viability was then tested using fluorescence-activated cell sorting (FACS) analysis with Annexin-V-FITC and propidium iodide. FACS analyses demonstrated that, despite the low growth rate, 86% of LY-B cells grown in SL-deficient medium for 72 h remained viable (Fig. 1C). Additional FACS results are shown in Fig. S2. Ethanol-treated CHO cells (Fig. S2A) were used as a positive control for non-viable cells. 95% CHO cells (Fig. S2B) and 95% LY-B cells (Fig. S2D) grown in standard medium were viable. With respect to cells grown in SL-deficient medium, 92% CHO (Fig. S2C) and 86% LY-B Fig. S2E), as well as 89% LY-B grown in SM-supplemented medium (Fig. S2F) were viable. Considering that even LY-B cells grown in 0.04 FBS retained a fair viability, these cells were considered as a good tool to obtain reliable information on the putative effects of a defective SPT activity on their biophysical properties.

#### Lysenin-staining

Cells were stained with SM-specific NT-lysenin-mCherry and visualized with confocal microscopy. LY-B cells grown in standard medium were stained not only with the general membrane stain NBD-PE (green) and also with lysenin fused to mCherry(red) (Fig. 1D). LY-B cells grown in SL-deficient medium appeared thoroughly stained with NBD-PE, but only little dots of mCherry were seen (Fig. 1E), indicating a remarkable decrease of SM in this sample.

In Fig. S3 mCherry-lysenin staining and FACS quantification is shown in detail. CHO (Fig. S3A) and LY-B (Fig S3B) grown in standard medium were both stained with lysenin and NBD-PE, but FACS analysis revealed that mCherry intensity was lower in LY-B cells (Fig. S3B) than in CHO ones (Fig. S3A). Moreover, when CHO cells were grown in SL-deficient medium, staining remained almost unchanged during the experiment (Fig. S3C). However, with LY-B cells grown in deficient medium (Figs. S3D and S4), lysenin fluorescence was largely decreased in the first 24 h of growth, and 48 h were enough to achieve a full lysenin-mCherry signal depletion (Fig. S4). In parallel with these observations (Fig. S5), the same behavior was detected in PM patches, prepared as described previously^37,42^ (Fig. S5). PM patches from LY-B cells grown in deficient medium for 72 h (Fig. S5D) exhibited very little lysenin-mCherry signal as compared with other PM patches (Fig. S5A, B, C).

In summary, both CHO and LY-B cell lines grew to a similar extent in standard medium. In SL-deficient-medium LY-B cells growth was lower than that of CHO cells, but they were still viable. Normal growth was recovered when the SL-deficient medium was supplemented with SM. Lysenin fluorescence was largely decreased in LY-B in the first 24 h of growth in deficient-medium, and 48 h were enough to achieve a full lysenin-mCherry signal depletion, while, under the same conditions, CHO cells showed only a slight decrease in lysenin-reacting SM.

### Membrane lipid order decreases with sphingolipid restriction

Laurdan generalized polarization (GP) experiments were performed to evaluate how the SPT activity suppression affected the rigidity/fluidity of cell membranes. GP provides an estimation of membrane lipid molecular order. GP values of whole cell lipid extracts, PM patches, GPMV and PM lipid extracts were measured using both two-photon microscopy and spectrofluorometry. In our previous work we performed laurdan GP experiments of CHO cell derived samples^37^. Now we have also measured GP values of SPT-suppressed LY-B cells. Figure 2A depicts a CHO cell grown in standard medium and stained with laurdan. Figure 2E illustrates the GP distribution of figure 2A, in a scale from −1 to 1, in which red color and a value close to 1 indicated a more rigid/ordered region, while blue color and a value close to −1 indicated a more fluid region. In figure 2A two different regions could be distinguished, the more ordered PM and the more disordered/fluid intracellular membranes. These two populations were clearly distinguished in the pixel intensity distribution graph (Fig. 2E), where two maxima were seen (blue line), respectively around 0.2 and around 0.5 as in Monasterio *et al.*^37^. The PM was less fluid than the intracellular membranes, perhaps due to its barrier role^51^. The red line in Fig. 2E indicates an average of the two peak intensities.

**Figure 2.**
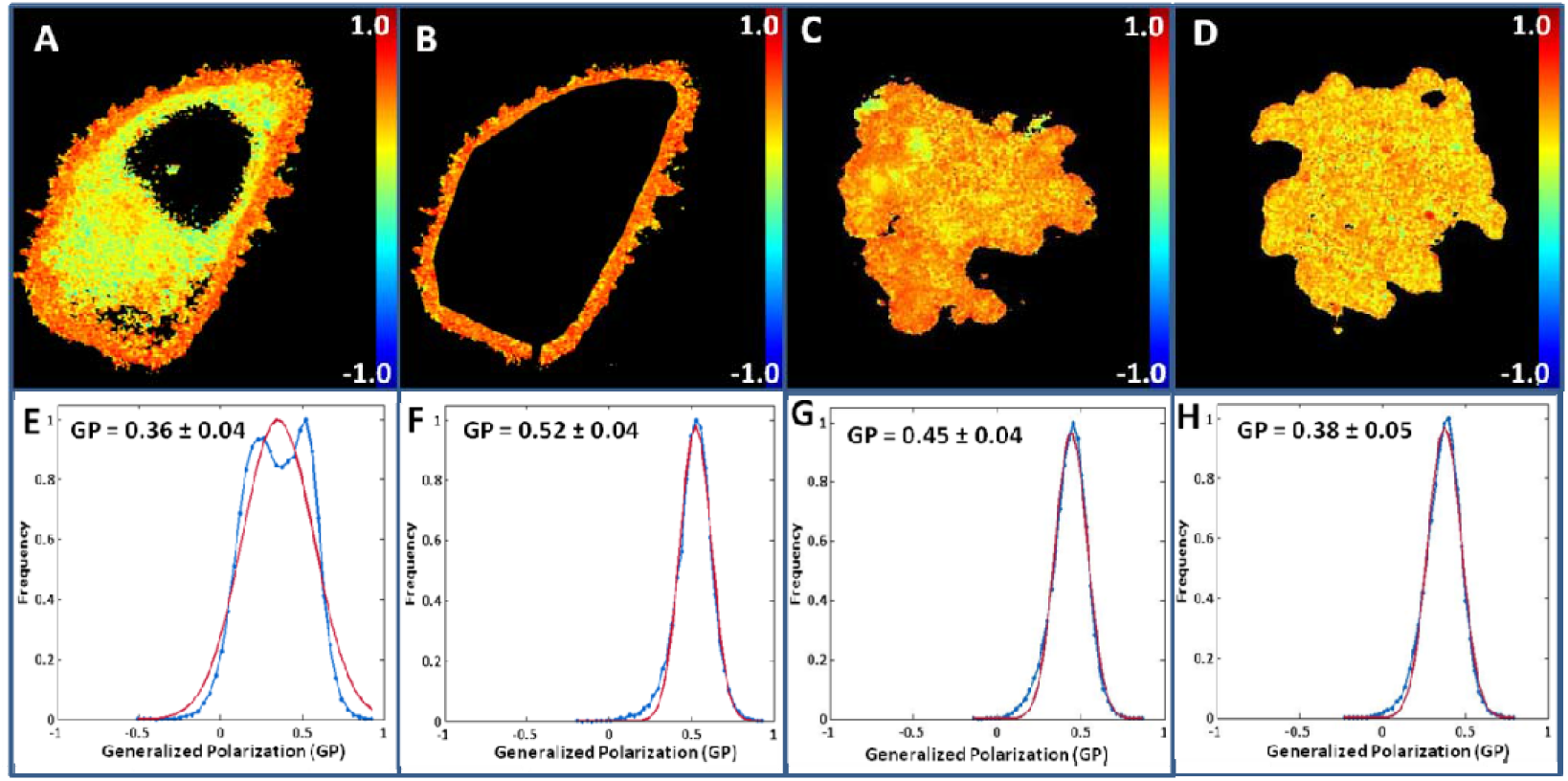
Laurdan staining and GP processing of cells and patches. A representative CHO cell grown in standard medium (**A**), and the PM selection of the cell in panel A (**B**). Representative PM patches of: a LY-B cell grown in standard medium (**C**), a LY-B cell grown in SL-deficient (0.04% FBS) medium (**D**). The scales at the right hand of the images are pseudo-color for GP values. Laurdan generalized polarization plots of the structures in panels A (**E**), B (**F**), C (**G**), and D (**H**) respectively. Blue lines indicate pixel intensity distribution; red lines indicate average of the blue line. Note in E that we can distinguish two different regions (blue line): the more rigid PM and the more fluid intracellular membranes.

The present study is focused on the PM. In figure 2B, a selection of PM pixels was performed in order to discard intracellular GP signal. Figure 5F shows the pixel intensity distribution of figure 2B: PM had a homogenous GP value around 0.5 as in Monasterio *et al.*^37^. A similar selection was performed with CHO and LY-B cells grown in standard and SL-deficient medium for 72 h. As seen in table 1 the GP values of the selected PM region decreased when FBS in the medium was lowered to 0.04%; LY-B cell PM GP went from 0.52 down to 0.43, while CHO cell GP value decreased more moderately, from 0.52 to 0.47. Student’s t test revealed that the difference between cells grown in standard and SL-deficient medium was statistically significant, both for CHO and LY-B cells. GP images of PM appeared to be homogenous, but the presence of nanodomains could not be ruled out because of the spatial and temporal resolution limit of conventional two photon microscopy^59–61^. GP values of isolated PM preparations were also measured. GP of PM patches derived from LY-B cells decreased on average from 0.45, for cells grown in standard medium (Fig. 2C, G), to 0.38, for those grown in SL-deficient medium (Fig. 2D, H).

Figure S6 shows examples of all CHO and LY-B cell-derived samples that were imaged with laurdan. The corresponding GP values and the statistically significant differences according to Student’s t test are summarized in Table 2. First, two different PM preparations (PM patches and GPMV) were measured. With GPMV (Fig. S6A and S6B) and PM patches (Fig. S6E and S6F) no significant differences between CHO and LY-B cells grown in standard medium were found. However, when cells were grown in SL-deficient medium, the GP value decreased in both cell lines. This was observed for all samples, but was particularly noticeable for LY-B cells and their derived PM preparations. This is an indication that LY-B cells grown in SL-deficient medium possess more fluid/less ordered membranes than when grown in standard medium.

**Table 2.** Laurdan GP values obtained from two-photon microscopy images. Statistically significant differences were calculated with ANOVA and Student’s t-test, with similar results. n=150. Significance of differences between CHO 10 and CHO 0.04: (#) p<0.05; (##) p<0.01. Differences between LY-B 10 and LY-B 0.04: (***) p<0.001.

|  | **CHO 10** | **CHO 0.04** |  | **LYB 10** | **LYB 0.04** |  |
| --- | --- | --- | --- | --- | --- | --- |
| **PM selection of 2-photon microscopy images** | 0.52 ± 0.04 | 0.47 ± 0.05 | ## | 0.52 ± 0.06 | 0.44 ± 0.04 | *** |
| **GPMV** | 0.44 ± 0.04 | 0.40 ± 0.03 | ## | 0.45 ± 0.04 | 0.36 ± 0.04 | *** |
| **PM patches** | 0.45 ± 0.03 | 0.41 ± 0.04 | # | 0.45 ± 0.04 | 0.38 ± 0.05 | *** |
| **GUVs of PM patches lipid extract** | 0.40 ± 0.05 | 0.36 ± 0.05 | # | 0.38 ± 0.03 | 0.31 ± 0.04 | *** |
| **GUVs of whole cell lipid extract** | 0.22 ± 0.03 | 0.13 ± 0.04 | ## | 0.18 ± 0.04 | 0.06 ± 0.03 | *** |

Mechanically, proteins are usually regarded as rigid inclusions in the lipid bilayer. Many theoretical studies have proposed that a membrane will become stiffer due to the presence of these rigid bodies^62^. Taking into account that the membrane bilayer is crowded with proteins, which may affect the rigidity/fluidity index, GUVs were formed with whole cell and PM lipid extracts as in Monasterio *et al.*^37^ and their GP values were also measured (Fig S6 and Table 1). The data in Figure S6 show that, according to GP values, the bilayers formed with the PM lipid extract of LY-B cells grown in SL-deficient medium (Fig. S6L) were significantly more fluid than those formed with the corresponding CHO PM lipids (Fig. S6K) (Table 1). GUVs formed with whole cell lipid extract were also studied. Again, statistically significant differences existed (Table 2) between LY-B grown in deficient medium and those grown in standard medium (Fig. S6N and S6P).

GP values in whole cell lipid extracts (cells grown in standard medium) were lower than the corresponding values in whole membranes, probably because of the rigidifying effect of proteins^62–65^, and a statistically significant difference existed between GP values of CHO and LY-B GP lipid extracts. To confirm this point, SUV formed from whole cell lipid extracts were used to measure the emission spectrum of laurdan at different temperatures as in Monasterio *et al.*^37^. The spectrum at 20°C showed only slight differences between the 4 samples (Fig. S7A). All samples had their emission maxima around 440 nm, indicating the rigid behavior of the membrane at this temperature. Still, the GP values derived from these emission spectra showed statistically significant differences between CHO and LY-B cells grown in standard medium, GP = 0.20 and 0.16 respectively (Fig. S7D). This difference increased when cells were grown in SL-deficient medium, with GP = 0.14 and 0.04 respectively (Fig. S7D).

At 30°C (Fig. S7B), the emission maxima of samples grown in deficient medium exhibited a spectral component centered at ≈490 nm. This was particularly clear in the spectrum of SUV formed from whole cell lipid extract of LY-B grown in SL-deficient medium, indicating a more fluid behavior. These effects were even more marked when the temperature was increased to 40°C. GP values derived from these emission spectra are summarized in figure S7D. Laurdan emission spectra of PM preparations at 40°C were also recorded (Fig. S8). In most cases the spectra were red shifted (towards higher disorder) when the SL source was made scarce, however GPMV samples appeared to keep their degree of high order irrespective of the SL source.

Thus, when CHO and LY-B cells were grown in standard medium, PM preparations of both cell lines exhibited similar GP values (around 0.45). This parameter suffered a larger decrease in LY-B (to around 0.37) than in CHO (to around 0.40) when cells were grown in SL-deficient medium. However, LY-B lipid extract GP values were lower than the corresponding CHO samples when cells were grown in either SL-deficient medium or in standard medium.

### Breakthrough forces of plasma membranes decrease with sphingolipid restriction

Whole cell lipid extracts, PM patches and GPMV were studied with an atomic force microscope (AFM), mainly in the force spectroscopy mode, to detect putative differences in breakthrough forces between the various samples. In our previous work we performed comparative AFM experiments of exclusively CHO-cell derived samples^37^. In this work we extend those findings, measuring also SL-depleted cells. Figure 3 shows the topology of PM patches from LY-B cells grown in standard (Fig. 3A) or SL-deficient (Fig. 3B) medium. The corresponding images of PM patches from CHO and LY-B cells are shown in parallel in Fig. S9. According to topographic images, a minimum thickness of 5 nm was regularly observed in all samples, corresponding to the thickness of a lipid bilayer^66^. Other membrane components, perhaps (glyco)proteins or protein aggregates, gave rise to the higher elements in the AFM pictures. No significant differences were seen between PM patch thicknesses of the various samples. The breakthrough force distributions showed no significant differences between the PM patches from CHO (4.52 nN) and LY-B (4.64 nN) cells grown in 10% FBS standard medium (Fig. 3C). Nevertheless, when the FBS contents in the medium was 5% or lower, the breakthrough forces decreased gradually for both samples, but particularly in PM patches from LY-B cells (Fig. 3C). Significant differences existed between PM patches of both cell lines when FBS in the medium was 5% or lower (Fig. 3C). With the minimum FBS concentration in the growth medium (0.04%), the breakthrough forces decreased from 4.52 (standard medium) to 3.76 nN (SL-deficient medium) for CHO cells, while LY-B cell PM patches underwent a larger decrease from 4.64 nN (standard medium) to 2.98 nN (deficient medium).

**Figure 3.**
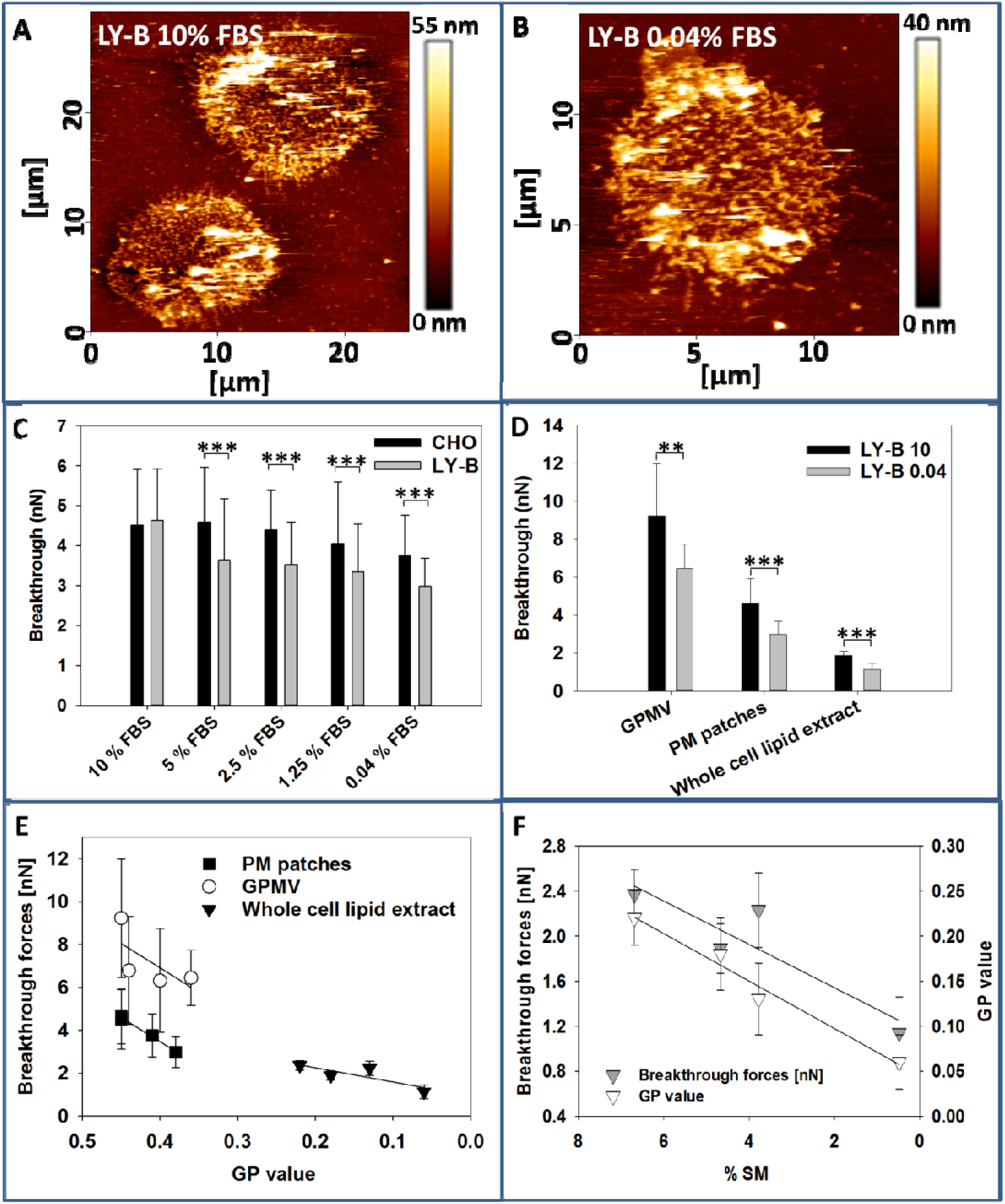
AFM measurements and correlation with laurdan GP and composition data. Topographic image of PM patches from LY-B cells grown in **(A)** standard or **(B)** sphingolipid-deficient (0.04% FBS) medium. **Force spectroscopy: (C)** Breakthrough forces for PM patches from CHO (black bars) and LY-B (gray bars) cells grown in media containing different FBS concentrations. **(D)** Breakthrough forces for GPMV, PM patches and whole cell lipid extracts from LY-B cells grown in standard (black bars) or sphingolipid-deficient (0.04% FBS) (gray bars) medium. Average values ± S.D. (n= 150-170). Statistical significance was calculated with ANOVA or Student’s t-test, with similar results: (*) p<0.05; (**) p<0.01; (***) p<0.001. **(E) AFM – laurdan fluorescence correlation**. The correlation between Laurdan GP and bilayer breakthrough forces are shown as regression lines. The experimental points correspond to PM patches (full squares, y_0_ = −5.54 ± 0.59; a = 22.51 ± 1.39; r = 0.992), GPMV (empty circles, y_0_ = −2.17 ± 7.13; a = 22.70 ± 17.22; r^2^ = 0.465) and whole cells (full triangles, y_0_ = 0.93 ± 0.50; a = 6.62 ± 3.14; r^2^ = 0.689). **(F) Correlation between [SM] and physical properties**. The correlations between SM concentration in bilayers (whole cell lipid extracts), and Laurdan GP and bilayer breakthrough forces, are shown as regression lines. Breakthrough forces: (grey triangles, y_0_: 1.16 ± 0.29; a: 0.19 ± 0.06; r:0.82) or GP values: (empty triangles, y_0_: 0.045 ± 0.013; a: 0.026 ± 0.000; r^2^:0.97). Average values ± S.D. (n = 3-5).

GPMV breakthrough forces were also measured. In GPMV derived from LY-B cells they decreased significantly when cells were grown in SL-deficient medium (9.23 in standard vs. 6.45 in deficient medium). Breakthrough forces of GPMV derived from CHO cells grown in standard and deficient media were also assessed (Fig S11), but no significant differences were found in this case. For lipid-supported planar bilayers (SPB) Figure 3D shows that the breakthrough force required for piercing the SPB from LY-B lipid extract decreased significantly when cells were grown in deficient medium (1.89 nN in standard medium vs. 1.14 nN in SL-deficient medium).

Topographic images of SPB from whole CHO and LY-B cell lipid extracts grown in standard medium were also scanned (Fig. S10). The images did not exhibit statistically significant differences; thickness was around 5 nm in all cases and lipid domains could not be distinguished. Breakthrough forces of SPB formed with the CHO cell lipid extract were also measured (Fig S11). In agreement with the results obtained from lipid extract GP measurements (Table 2), the breakthrough forces were significantly different between CHO (Fig. S10A) and LY-B (Fig. S10B) cells grown in standard medium (Fig. S11). This difference became larger when CHO (Fig. S10C) and LY-B (Fig. S10D) cells were grown in deficient medium (Fig S11).

Breakthrough force measurements were in good agreement with laurdan GP values. Whole cell lipid extracts and PM patches exhibited the expected behavior, the LY-B cells grown in deficient medium showing the lowest values in both techniques (Fig. 3E,F). A linear correlation was observed between changes in SM levels and changes in GP and breakthrough forces (Fig. 3E,F).

To summarize, CHO and LY-B PM patch breakthrough forces had similar values (around 4.5 nN) when the cells were grown in standard medium. In deficient medium this value decreased more markedly for LY-B (2.98 nN) than for CHO (3.76 nN). As to breakthrough forces in GPMV, those in vesicles derived from LY-B were also decreased (from 9.23 nN to 6.45 nN), but not those obtained from CHO. Finally, lipid extract GP value differences were also appreciable when cells were grown in standard medium (1.89 nN for LY-B and 2.37 nN for CHO). The latter values decrease in LY-B (to 1.14 nN), but not in CHO (2.23 nN), when growing in SL-deficient medium.

### Sphingolipid composition changes after a 250-fold reduction in sphingolipid supply, with little change in glycerophospholipids

Analyzing the lipid redistribution undergone by cells with suppressed SPT activity is crucial for a proper understanding of the existing membrane order/disorder differences between CHO and LY-B cells in high- and low-SL media, and their capacity of homeostatic regulation. In order to clarify the changes in the cell lipid pathways, a lipidomic study of whole cell and PM preparations of CHO and LY-B cells grown in standard and deficient media was performed. For GPMV formation, but not for PM patch preparations, cells had been in contact with dithiothreitol and paraformaldehyde, thus different controls were used for each case as in Monasterio *et al.*^37^.

Figure 4A-C shows the amounts of different SL (SM, Cer and HexCer) when CHO and LY-B cells were grown in medium containing different FBS concentrations. These three specific SL were selected among the lipidomic data respectively because SM is the most abundant SL, Cer is particularly important in cell signaling, and HexCer is at the origin of the biosynthetic pathway leading to the complex glycosphingolipids. Significant differences were seen between SM amounts in CHO and LY-B cells (Fig. 4A). When cells were grown in standard medium, the SPT-deficient LY-B cells contained 43% less SM than CHO cells. This suggests that over one half of the total SM is synthesized by the *de novo* pathway.

**Figure 4.**
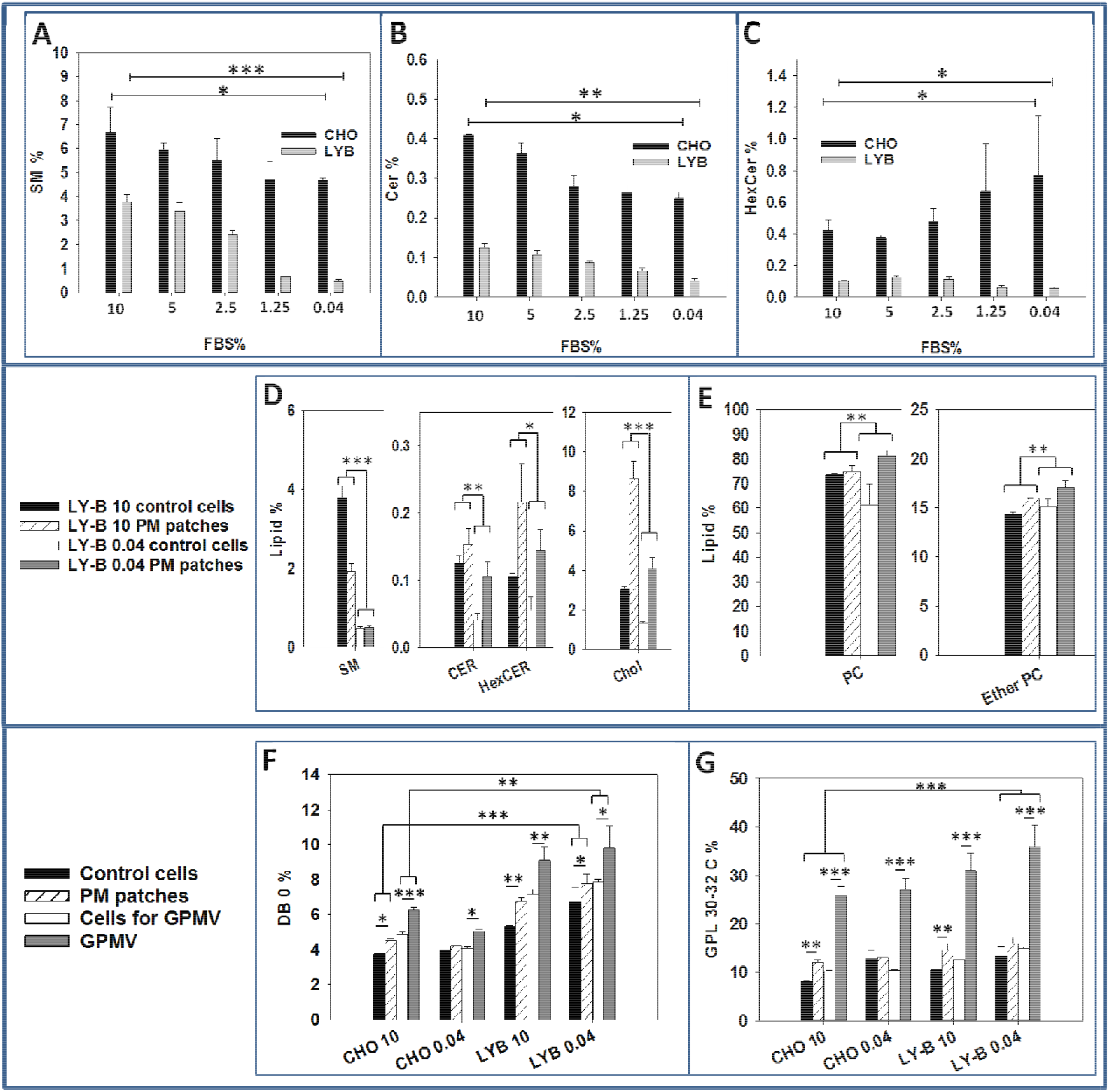
Lipidomic analysis of whole cells and plasma membrane preparations. Total SM (**A**), Cer (**B**) and HexCer (**C**) from CHO (black bars) or LY-B (gray bars) whole cells grown in media containing various concentrations of FBS. A comparison of lipid compositions of whole cell and PM patches of LY-B cells grown in standard or sphingolipid-deficient (0.04% FBS) media (**D, E**). Only selected lipids are included in the figure. A comprehensive description of the various lipid compositions can be seen in the Supplementary Material Table S1 and Figure S12. Fully saturated (DB = double bond) (**F**) and short-chain (30-32C) GPL (**G**) of whole cells treated for GPMV preparation, GPMV, cells treated for PM patch preparation, and PM patches. n=3. Statistical significance was calculated with ANOVA or Student’s t-test, with similar results. Significance: (*) p<0.05; (**) p<0.01; (***) p<0.001.

When the amount of FBS in the medium was decreased, total SM was also lower in both wild type and mutant cells. Nevertheless, the decrease was greater in LY-B than in CHO cells, 90% vs 25% with the lowest FBS concentration (0.04%) (Fig. 4A). In cells grown in standard medium, LY-B contained 70% less Cer than CHO cells. When FBS concentration was decreased, the total Cer amount was also diminished in both cell lines. As with SM, the reduction was larger in LY-B than in CHO cells, 66% vs. 38% (in 0.04% FBS-containing medium). Figure 4C shows the corresponding values for HexCer. LY-B grown in 10%-FBS medium contained 70% less HexCer than CHO, in agreement with the Cer data (Fig 4B). When FBS in the medium was decreased, HexCer also decreased by 50% in LY-B cells grown in 0.04% FBS-containing medium, but it increased by 85% in CHO (Fig. 4C). Considering the PM patches, SM, Cer and HexCer were all lower in LY-B 0.04 PM patches than in LY-B 10 ones. In summary, in LY-B cells the three SL under study exhibited a similar decrease (55-70%) with the reduction of FBS, at variance with CHO cells, indicating that, in the latter, SL are largely synthesized by the *de novo* pathway. The situation with GPMV was, however, different, mainly in that Cer and HexCer increased when the cells (either CHO or LY-B) were grown in 0.04 FBS medium (Fig. S12B, C).

With low-FBS medium Chol levels were decreased in both cell lines (Fig. S12D). Since FBS is a major source of lipids and proteins for cell growth under our conditions, the drastic reduction from 10% to 0.04% in the SL-deficient medium induces partial cell starvation. Chol synthesis is known to decrease in fasting conditions^67–69^. Moreover, LY-B PM preparations had larger amounts of Chol than the whole cell average, for cells grown in high- and low-FBS (Fig.4D). This had been observed by Monasterio *et al.*^37^ for the case of CHO cells.

Glycerophospholipid (GPL) acyl chain also changed along with SL deprivation. In figure 4E we can see that ether PC was increased in LY-B 0.04 cells. GPL acyl chain saturation and length also play an important role in the physical properties of the membrane bilayer, specifically on its disorder/fluidity. Specifically, unsaturated and shorter acyl-chain containing GPL increase membrane fluidity^70^. In figure 4F the distribution of fully saturated GPL of control and two PM preparations (patches and GPMV) is shown. PM preparations had more fully saturated and less polyunsaturated (2-6 double bounds) GPL (Fig. S12J-L) than their respective controls in all measured samples^37^. The differences were larger in the case of GPMV. Comparing CHO and LY-B grown in standard medium, LY-B had more fully saturated GPL in all cases. In addition, when FBS in the medium was decreased, the saturated GPL increased in LY-B, while in CHO cells they remained almost constant (Fig. 4F).

As to the GPL chain length distribution, PM preparations were richer in 30-32 C chains than whole cells, the difference being larger for the GPMV (Fig. 4G)^37^. GPL of LY-B grown in standard medium contained more 30-32 C acyl chains than CHO 10 in all measured samples. This difference was increased when LY-B were grown in 0.04% FBS medium. Conversely, CHO 0.04 chain-length values remained constant (Fig. 4G). In summary LY-B cells synthesized shorter and more saturated GPL in their homeostatic response to SM depletion. A comprehensive description of the various lipid compositions can be seen in the Supplementary Material Table S1 and Figure S13.

A quantitative estimate of the amount of SM and Chol, two representative lipids in this context, was carried out as described under Methods. The results can be seen in Table S2. With respect to Chol (Table S2), the concentration in CHO cells grown in SL-medium was 33% of those grown in standard medium (data in ng Chol/cell), and the corresponding figure for LY-B was 45%. As for SM data, CHO cells grown on medium with 0.04% FBS contained 68% of the SM found in cells grown on 10% FBS (data in ng SM/cell), or 53% (in fg SM/μg protein). For the SPT-defective LY-B cells, the corresponding figures are 15% and 16%. Thus, a 250-fold reduction in sphingolipid supply to LY-B cells leads to a 6-fold decrease in membrane sphingolipids.

As a summary of the lipidomic results, SM, Cer and HexCer concentrations were lower in LY-B PM patches than in CHO ones when grown in standard medium. All three SL were similarly decreased (55-70%) with the reduction in FBS. LY-B cells contained larger amounts of Chol than CHO ones in both standard and SL-deficient media. With respect to the GPL fatty acyl distribution, LY-B had more saturated and shorter GPL fatty acids than CHO cells. These groups of fatty acids, together with PC ethers, were increased in LY-B and maintained in CHO when FBS concentration was decreased. For LY-B in SL-deficient medium SM decreased both in GPMV and in PM patches but Cer and HexCer were increased with lower FBS concentrations (Fig. S12A-C).

## Discussion

LY-B cells grown in SL-deficient medium were used to understand the effects that a defective SPT activity might have on the biophysical properties of the cell. SPT-defective cells grown with very low SL concentrations were viable (Fig. 1E) and they were able to recover the control growth rates when the SL-deficient medium was supplemented with SM (Fig. 1B).

### CHO and LY-B cells grown in standard medium

#### Whole cells

Comparing SL levels in CHO and LY-B cells grown in standard medium, they happened to be markedly lower in the mutant cells. Considering the three most abundant SL, SM was 43% lower in LY-B cells (Fig. 4A), Cer was 66% lower (Fig. 4B), and HexCer was 70% lower (Fig 4C). This indicated that the *de novo* pathway was a major SL synthesis source. This result is in agreement with the one published by Ziulkoski *et al.*^30^ where they used fumonisin B1 and β-chloroalanine to determine the contribution of the different pathways to the synthesis of SM in Sertoli cells. They found that 40% of 16:0 and 61% of 18:0, 18:1 and 18:3 SM was synthesized by the *de novo* pathway. They also observed that these values could be increased when the requirement for cell membranes was greater, as in rapidly dividing cells^30^. In contrast with the SL results, Chol concentrations were similar in CHO and LY-B cells grown in standard medium (Fig. S12D). In addition, comparison of both kinds of cells in standard medium showed small changes in GPL, LY-B contained more fully saturated, and less monounsaturated GPL than CHO, chain length distributions in GPL being virtually the same (Fig. 4F,G).

#### PM preparations

Important differences were found between the whole cell and PM lipid compositions, as anticipated from the studies in CHO by Monasterio et al.^37^ Both for CHO and LY-B preparations, PM patches contained less SM (about one half), more HexCer and more Chol than the whole cells. GPMV had also less SM but contained higher amounts of HexCer, and particularly of Cer and Chol. Changes in GPL were moderate or low, except for cardiolipin, that was almost absent in the PM preparations (Fig. S12A-D, I). The higher amounts of Chol in PM patches (Fig. S12D) could be a major factor responsible for compensating PM molecular order even with lower SM levels. Neither laurdan GP of GPMV nor laurdan GP or breakthrough forces of PM patches showed any statistically significant difference between CHO and LY-B grown in 10% FBS (Table 2, Fig. 3C).

GPMV constitute a frequently used PM preparation ^37,40,41,71^. However, the lipidomic data show that their lipid composition departs from those of the whole cells and from other PM preparations (patches). In particular, GPMV were enriched in Cer (Fig. S12B) and HexCer (Fig. S12C), and they also exhibited an unusual enrichment in PI (Fig. S12G). Furthermore, their GPL are enriched in saturated fatty acids (Fig. S12J), and contain correspondingly less unsaturated chains, specifically with 2-6 double bonds per GPL molecule (Fig. S12L). Also, the proportion of medium-length fatty acids (C30-32 per GPL molecule) increases at the expense of the longer ones (C34-40) (Fig. S12 M,N). All these are peculiarities of GPMV, in which they differ from all other cell and membrane preparations, with either 10% or 0.04% FBS. The fact that these changes are not modified by SL depletion, and that some of them affect mainly GPL, makes GPMV a less useful membrane preparation in the context of our study. GPMV penetration required consistently higher breakthrough forces than PM patches (Fig. S11). This could be related to the enrichment in Cer and HexCer found in GPMV with respect to whole cells (Fig. S12B, C). Both Cer and HexCer are known to increase membrane lipid order^72–74^ GPMV have been shown to be permeable to hydrophilic macromolecules^71^, and this could again be related to the increased Cer, and partly HexCer, since these SL happen to increase membrane permeability^72,73^. The observed increases in GP and breakthrough forces could be secondary to the use of dithiothreitol in GPMV formation. As seen in Epstein *et al*^75^. dithiothreitol can be responsible for increasing Cer concentrations even in SPT-suppressed cell lines, without altering SM values. Those authors concluded that dithiothreitol could induce the ‘unfolded protein response’ and this would lead to an over-expression of the SPT LCB1 subunit mRNA, partially recovering its activity^75^. Dithiothreitol was also shown to affect lipid–lipid and lipid– protein interactions and to integrate directly into lipid membranes^76^.

### CHO and LY-B cells grown in SL-deficient medium

When LY-B cells were grown in SL-deficient medium, SM and Chol percent levels were markedly decreased, respectively by about 5-fold and 2-fold, with no comparable changes in Cer or HexCer, and the derived PM patches followed parallel trends (Fig. 4A,D, Fig. S12A,D). In CHO cells the decrease in SM concentration was less clear, and HexCer levels actually increased somewhat, other SL varying as in LY-B (Fig. 4A,C), with the corresponding PM patches showing similar trends (Fig. S12A-D). Growth in SL-deficient medium did not cause any remarkable changes in GPL, nor in their associated fatty acids (Fig. S12E-N), with the exception that the very long fatty acids (C42-44 per GPL molecule) whose concentration was in any case very low, were further decreased with the low FBS medium. Note that the largest decrease in SM, the most abundant SL, occurs in LY-B cells deprived of SL in the growth medium, thus the two factors are required, lack of SL in the nutrients and lack of capacity to synthesize the sphingosine precursor, to obtain low-SL cells.

With respect to the PM preparation, laurdan GP indicated a decreased lipid order (increased bilayer fluidity) in all samples under study (Table 2) and breakthrough forces decreased accordingly, more in LY-B than in CHO cells and membranes (Fig. 3B,C). As a result, PM patches from LY-B cells were less ordered and more easily penetrable (Table 2, Fig. 3C). The close correlation between decrease in SM concentration in cell membranes, as a result of SL deprivation in the nutrients, decrease in GP values and decrease in breakthrough forces can be seen in Fig. 3C,D. (Only data from whole cell lipid extracts are included in Fig. 3D, for simplicity).

### Homeostatic adaptations

At least some of the observed changes in membrane lipid composition as a result of gene suppression or of changes in nutrient media could be explained in terms of homeostatic responses to the novel situations. Perhaps the main observation in terms of adaptation is the remarkable resilience of LY-B cells that, when grown under extremely low SL concentrations (250-fold below standard conditions), are still able to divide while keeping SL concentrations 6-fold lower than standard and membrane physical properties not far away from the wild-type cells. Examining the data in more detail, and specifically comparing CHO and LY-B cells grown in 10% FBS medium, hints on adaptation to lack of *de novo* SL synthesis could be retrieved. In particular, as described above, the only notable change between the lipidomes of those two cell lines, grown under standard conditions, is the clear decrease in SL as percent total lipids (one-half on average) in the LY-B cells (Fig. S12), while Chol levels did not vary. The percent concentration of SM, the most abundant sphingolipid, went from 6.7% to 3.8% (Fig. S12A). Parallel changes were recorded in PM patches derived from those cells. This was not accompanied by any changes in the measured physical properties of the membranes, laurdan GP (Table 2) or AFM breakthrough forces (Fig. 3C). Perhaps the observed variation in SL concentration was not enough to cause any observable physical changes, and a very minor, or no adaptation was required.

The situation was different when cells were grown in SL-deficient medium. In LY-B cells SM concentration dropped by one order of magnitude when cells were grown in 0.04% instead of 10% FBS. Other SL, as well as Chol, were decreased in parallel. PM patches underwent similar changes as the whole cell lipids. Perhaps as a consequence of these changes, the membranes became more easily penetrable, and lipids became less ordered (Table 2 and Fig. 3) when FBS concentration was lowered. Simultaneously, fatty acyl unsaturation was decreased (Fig. S12J-L), a phenomenon that could have the effect of increasing lipid order, thus tending to compensate the decrease in bilayer-ordering SM.

When CHO cells were grown in SL-deficient medium the proportion of HexCer was considerably increased, by about 2-fold. This increased HexCer synthesis (that could not occur in LY-B cells because of their low sphingosine availability, due to the lack of SPT activity), may be one of the homeostatic responses that wild-type cells carry out under starvation. HexCer is at the origin of the complex glycosphingolipid biosynthetic pathway^1^. Glycosphingolipids are required for cellular differentiation and there are human diseases resulting from defects in their synthesis^77^. This may be one of the reasons for the different dividing ratios of CHO and LY-B in SL-deficient medium (Fig. 1A). CHO, but not LY-B cells, may over-express the HexCer synthesis to continue cell division in order to buffer the nutrient depletion condition. As discussed above, Chol levels in CHO and LY-B grown in standard medium remained invariant, and this could help in maintaining membrane rigidity under conditions of low SM (Fig. S12D,J). Nevertheless, as Chol synthesis decreases under cell starvation conditions, rigidity cannot be maintained in this manner under SL-deficient conditions. When FBS in the medium was decreased, saturated GPL were increased in LY-B (Fig. 4F), while in CHO cells they remained almost constant.

### Concluding remarks

The present study has demonstrated that in cells lacking the SPT activity, SM, Cer and HexCer are markedly decreased in all measured samples (controls and PM preparations).

Fully saturated GPL are increased and polyunsaturated ones are decreased. Synthesizing more saturated GPL can be the way that LY-B cells have to compensate the low SM. Cholesterol may also have some influence in that response but its effect is minimized when its levels are decreased because of starvation. The SM-depleted cells try to maintain membrane order undergoing a homeostatic response, although they achieve it only partially as their PM are more fragile when grown in SL-deficient medium.

These changes in lipid order and membrane rigidity caused by low SL could be linked to a variety of phenomena in cell physiology and pathology. Alterations in the normal activities of the SM-cycle enzymes have been associated to many central nervous system and neurodegenerative diseases^19^. Specific SM species have been found to bind membrane proteins thereby modifying their functions^78^. The capacity shown by certain cells in this paper to grow under extremely demanding low concentrations of SL opens the way to a variety of functional studies on the role of SL in membranes.

## Acknowledgements

We thank Dr. Alfred Merrill (Georgia Tech) for suggesting that we examine the properties of LYB cells in sphingolipid-depleted medium. The authors are also grateful to Dr. D. Tyteca for the kind gift of plasmid pET28/Dronpa-NT-lysenin. This work was supported in part by grants from the Spanish Ministry of Economy (grant FEDER MINECO PGC2018-099857-B-I00) and the Basque Government (grants No. IT1264-19 and IT1270-19), as well as by Fundación Biofísica Bizkaia and the Basque Excellence Research Centre (BERC) program of the Basque Government, and by the Swiss National Science Foundation (310030-184949).

