## Supplementary material for "CHO/LY-B cell growth under limiting sphingolipid supply: correlation between lipid composition and biophysical properties of sphingolipid-restricted cell membranes^+^": Supp Mat.pdf

**Supplementary Figures S1 – S12, Supplementary Table 2**

### CHO/LY-B cell growth under limiting sphingolipid supply: correlation between lipid composition and biophysical properties of sphingolipid-restricted cell membranes[+].

Bingen G. Monasterio[1,2], Noemi Jiménez-Rojo[3], Aritz B. García-Arribas[1,2], Howard Riezman [3], Félix M. Goñi[1,2], Alicia Alonso*[1,2]

[1]Instituto Biofisika (CSIC, UPV/EHU) and [2]Departamento de Bioquímica, Universidad del País Vasco, 48940 Leioa, Spain; [3]NCCR Chemical Biology, Department of Biochemistry, University of Geneva, 1211 Geneva, Switzerland.

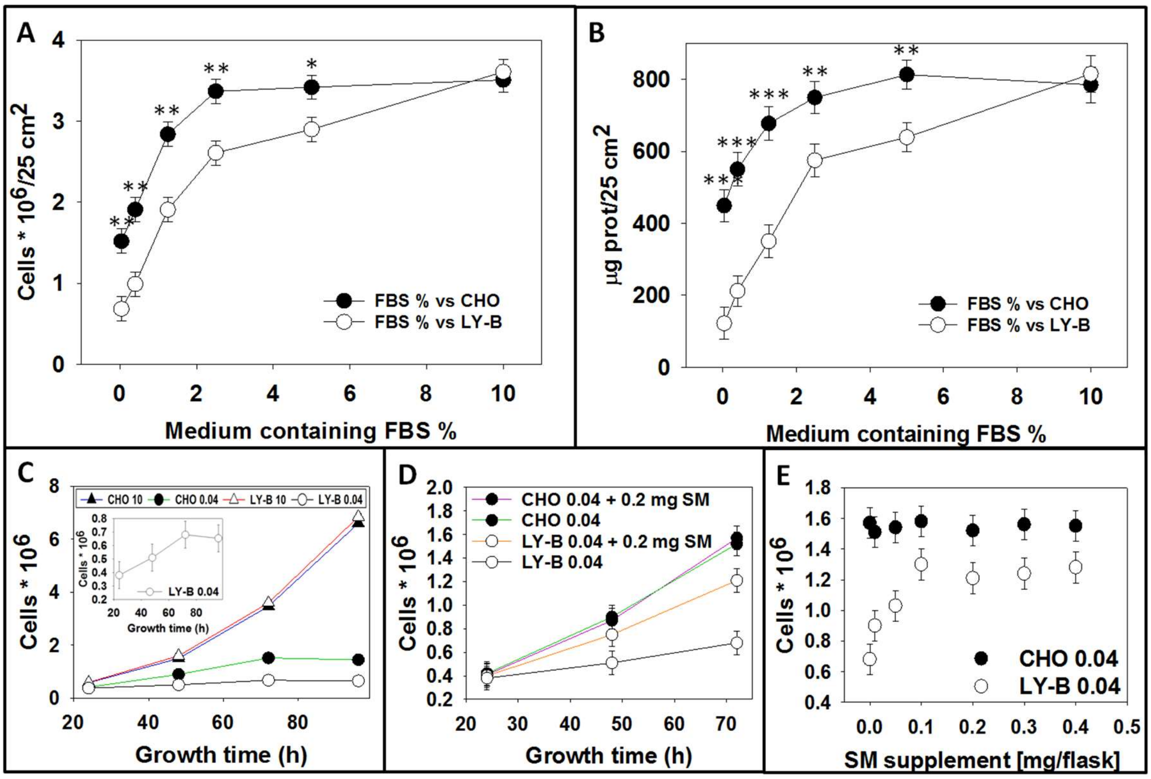


**Figure S1. CHO and LY-B cell growth measurements.** Cell counts (**A**), and protein contents (**B**), as a function of %FBS in the medium (growth time: 72 h). Cell counts of LY-B and CHO as a function of time in standard (10% FBS) and sphingolipid-deficient (0.04% FBS) medium (**C**), sphingolipid-deficient medium and sphingolipid-deficient medium supplemented with SM (**D**) and LY-B and CHO cell growth after 72 h in sphingolipid-deficient medium supplemented with SM (seeded cells: $0.25 * 10^6$) (**E**). Statistical significance was calculated with ANOVA and Student´s t-test. Significance: (*) $p<0.05$; (**) $p<0.01$; (***) $p<0.001$.

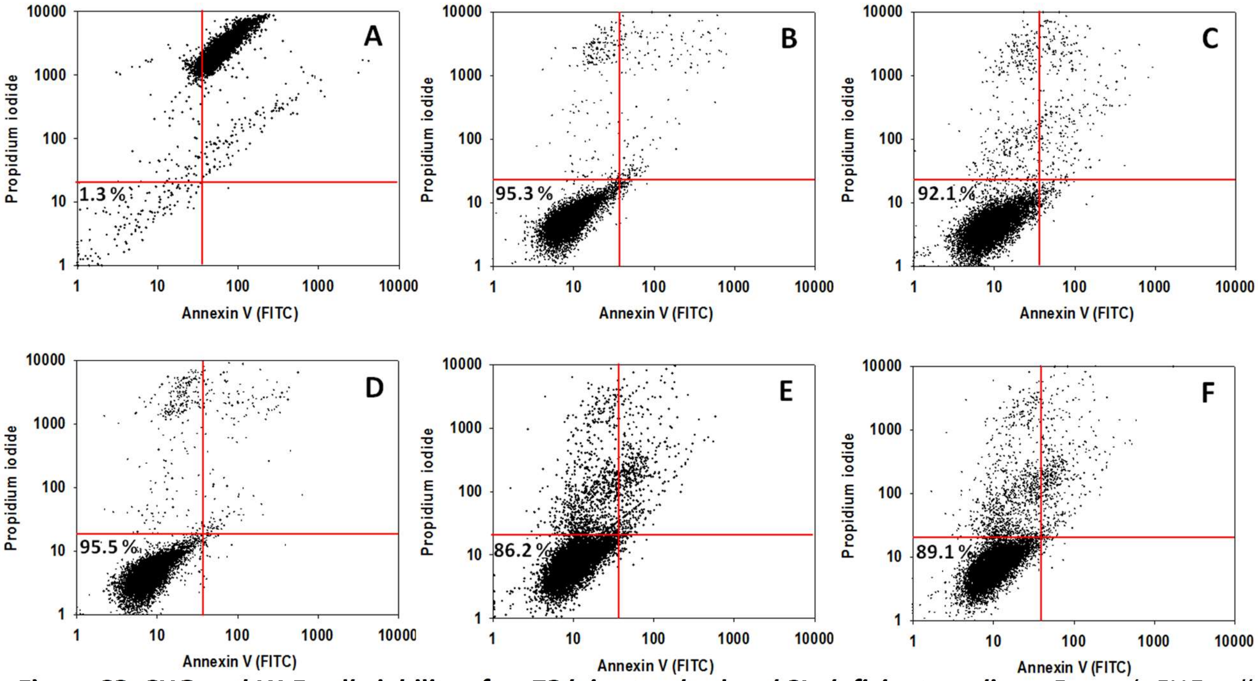


**Figure S2. CHO and LY-B cell viability after 72 h in standard and SL-deficient medium**. Control, CHO cells treated with EtOH (**A**). CHO 10 (**B**). CHO 0.04 (**C**). LYB 10 (**D**). LYB 0.04 (**E**). LYB 0.04 + 0.2mg SM (**F**).

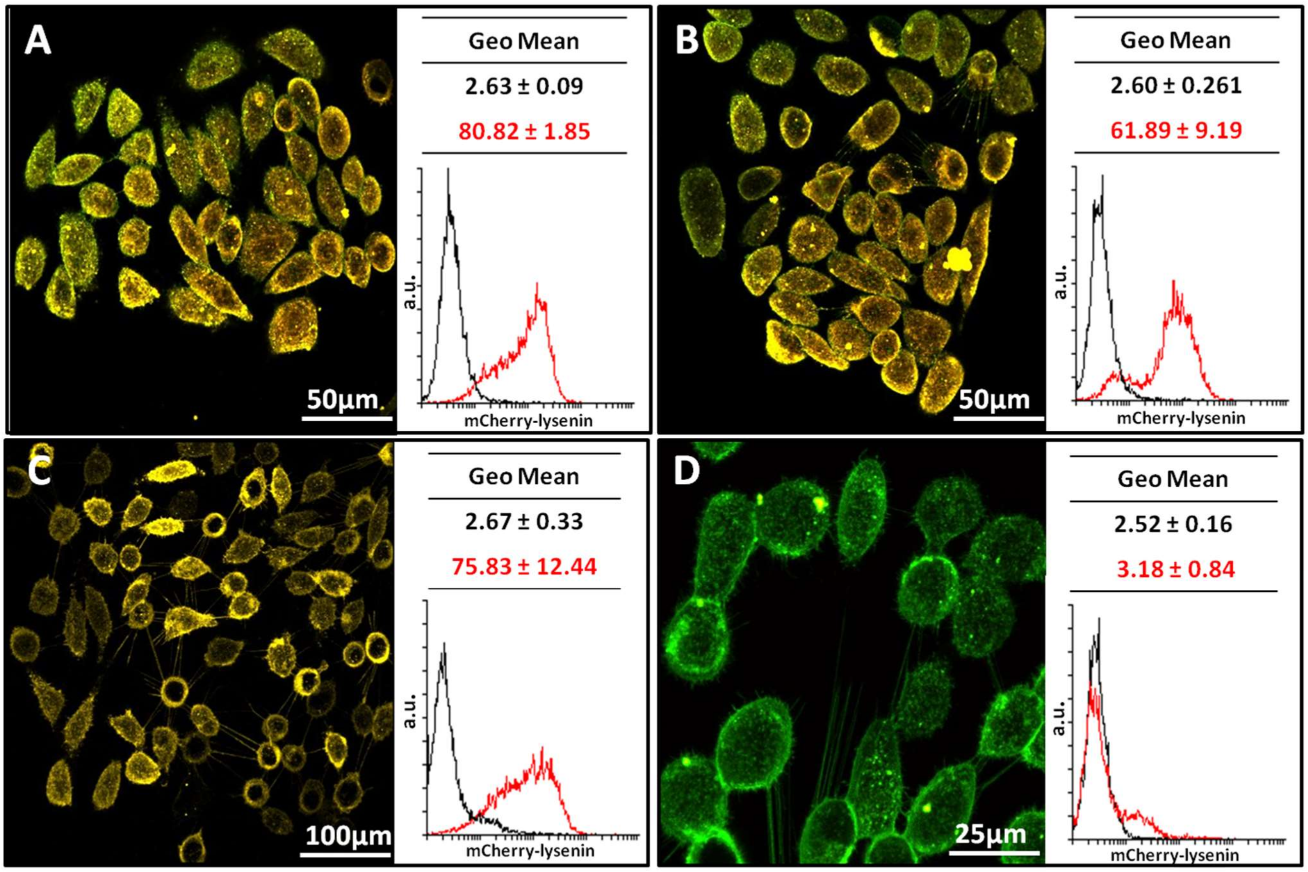


**Figure S3. Fluorescence images and FACS-mediated mCherry-lysenin (red) quantification.** Measurements are shown after 72 h growth. CHO (**A**) and LY-B (**B**) cells grown in standard medium**.** CHO (**C**) and LY-B (**D**) cells grown in SL-deficient medium. Black histograms correspond to control cells (without mCherry-lysenin staining) and red ones to the sample of interest (mCherry-lysenin signal). NBD-PE (green) was used in fluorescence images for general membrane staining. Geometric mean ± S.D. (n = 3).

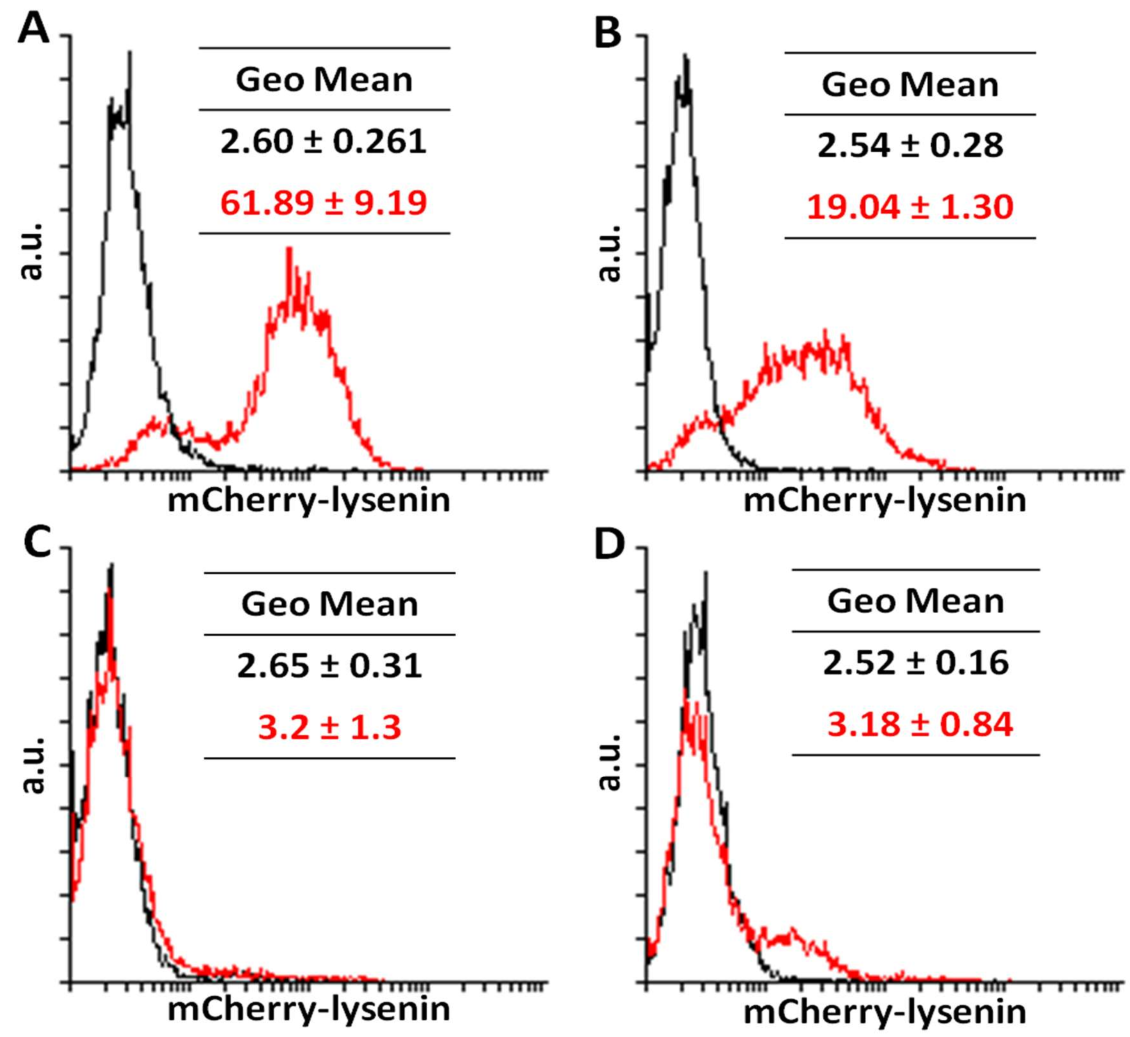


**Figure S4. FACS-mediated, time-dependent mCherry-lysenin quantification of cell growth.** LY-B cells grown in standard **(A)** and deficient medium during 24 h **(B),** 48 h **(C)** and 72 h **(D).** Black histograms correspond to control cells (without mCherry-lysenin staining) and red ones to the sample of interest (mCherry-lysenin signal). Geometric mean ± S.D. (n = 3).

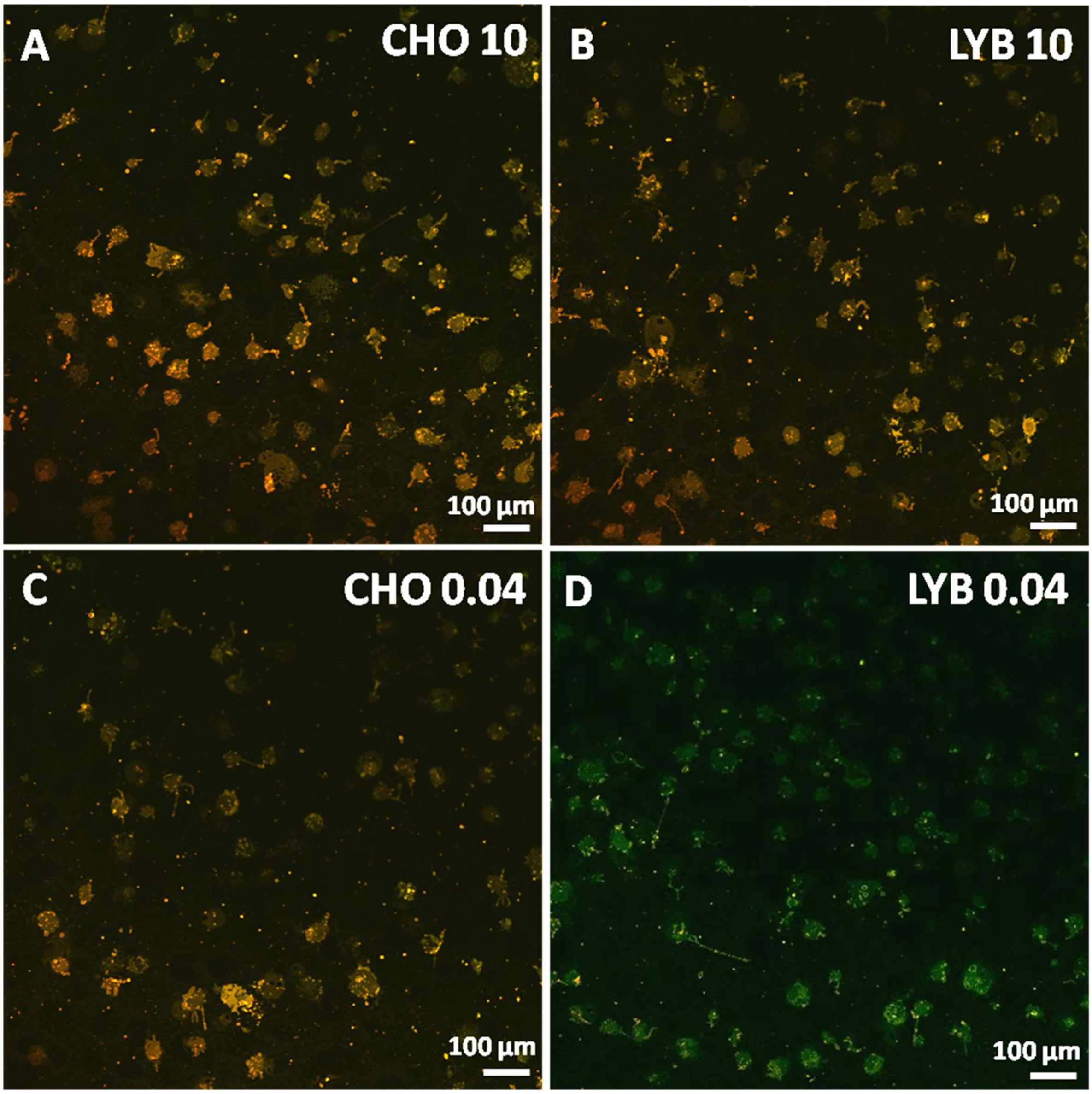


**Figure S5. Fluorescence images of PM patches stained with mCherry-lysenin (red).** CHO (**A**) and LY-B (**B**) cells grown in standard medium and CHO (**C**) and LY-B (**D**) cells grown in deficient medium. Bar = 100 µm. NBD-PE (green) was used in fluorescence images as membrane staining control.

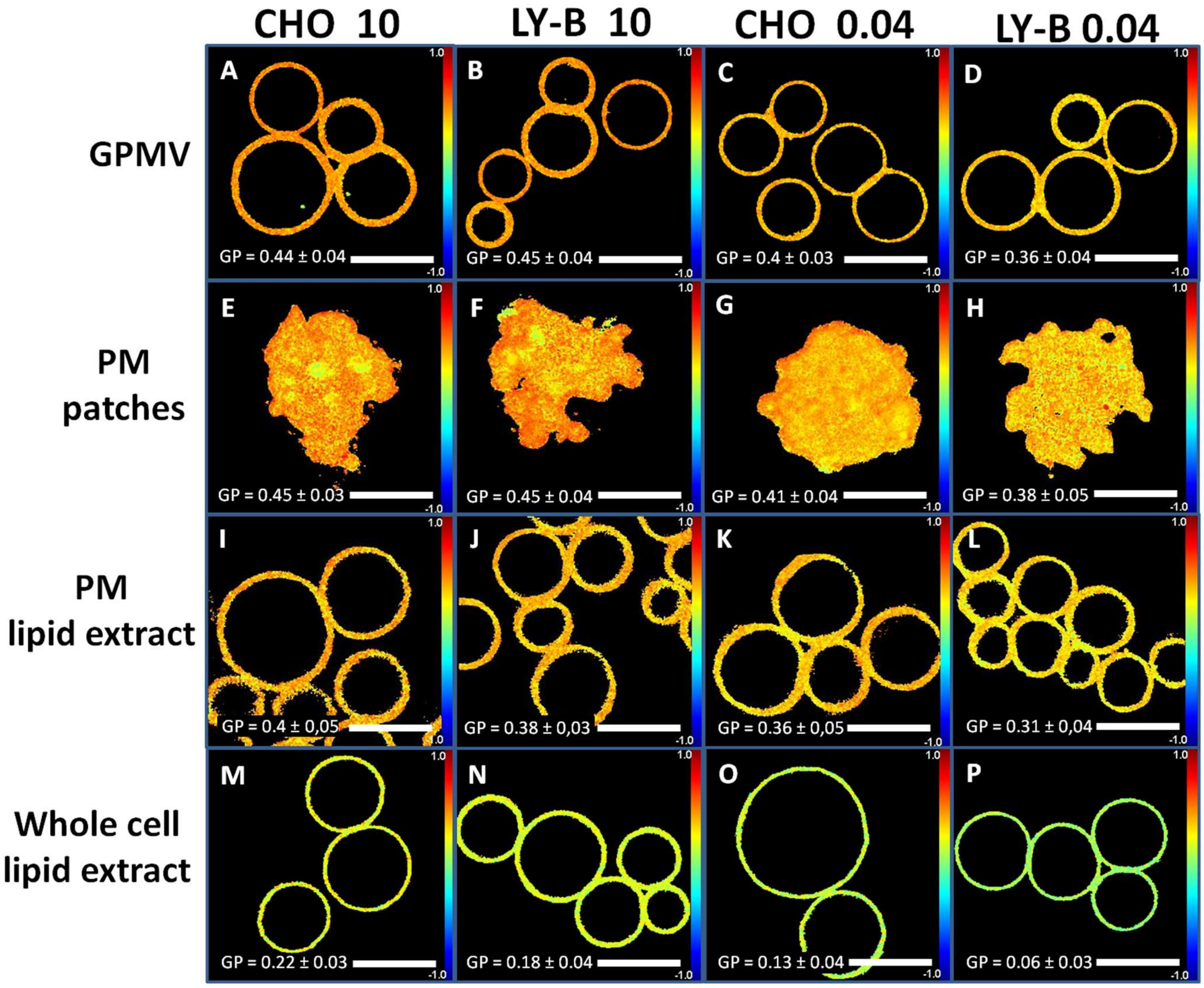


**Figure S6. Two-photon microscopy images of samples stained with Laurdan for their GP measurement. A-D** GPMV of: CHO cells grown in standard medium (**A**), LY-B cells grown in standard medium (**B**), CHO cells grown in deficient medium (**C**) and LY-B cells grown in deficient medium (**D**). Bar = 10 µm. **E-H** PM patches of: CHO cells grown in standard medium (**E**), LY-B cells grown in standard medium (**F**), CHO cells grown in deficient medium (**G**) and LY-B cells grown in deficient medium (**H**). Bar = 30 µm. **I-L** GUV formed from PM lipid extract of: CHO cells grown in standard medium (**I**), LY-B cells grown in standard medium (**J**), CHO cells grown in deficient medium (**K**) and LY-B cells grown in deficient medium. Bar = 10 µm. **M-P.** GUV formed from whole cell lipid extract of: CHO cells grown in standard medium (**M**), LY-B cells grown in standard medium (**N**), CHO cells grown in deficient medium (**O**) and LY-B cells grown in deficient medium (**P**). Bar = 10 µm.

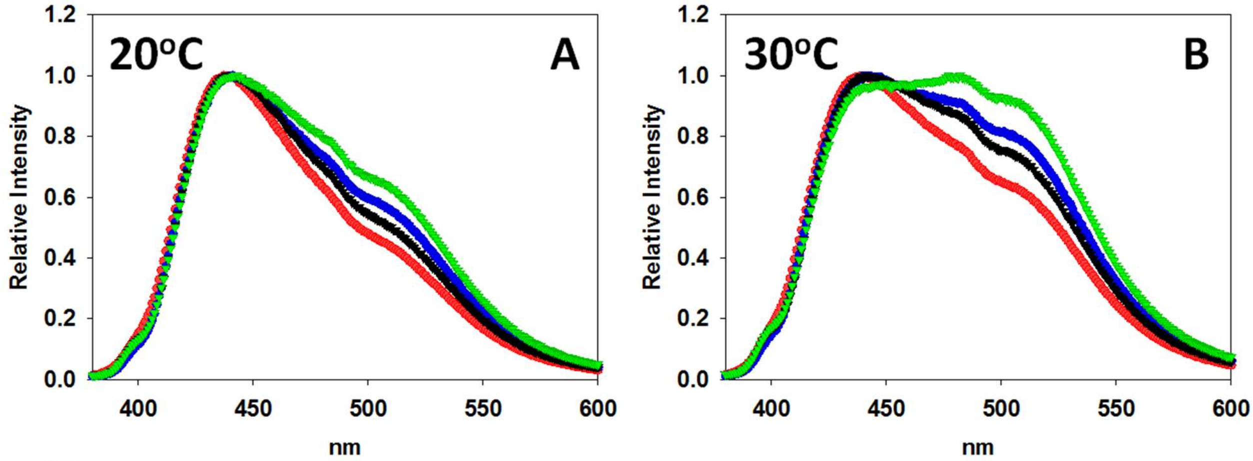


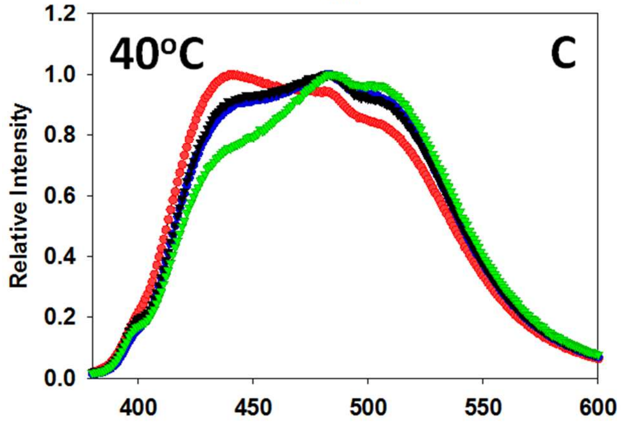


| T (ºC) | Medium | GP CHO ± SD | GP LY-B ± SD | D |
|---|---|---|---|---|
| 20 | normal | 0.20 ± 0.02 | 0.16 ± 0.04 | * |
| | deficient | 0.14 ± 0.03 | 0.04 ±0.01 | ** |
| 30 | normal | 0.09 ± 0.02 | 0.02 ± 0.05 | * |
| | deficient | 0.00 ± 0.03 | -0.11 ± 0.01 | ** |
| 40 | normal | 0.01 ± 0.02 | -0.11± 0.06 | ** |
| | deficient | -0.12 ± 0.03 | -0.27 ± 0.02 | *** |

**Figure S7: Laurdan emission spectra of SUV formed with whole cell lipid extracts.** Spectra at 20ºC (**A**), 30ºC (**B**) and 40ºC (**C**)**.** GP values (table) (**D**). Red spectra: SUV formed from CHO (10% FBS) lipid extracts. Blue spectra: SUV formed from CHO (0.04% FBS) lipid extracts. Black spectra: SUV formed from LY-B (10% FBS) lipid extract. Green spectra: SUV formed from LY-B (0.04% FBS) lipid extract. Values in the Table are averages ± S.D. (n = 3). Statistical significance was calculated with Student´s t-test: (*) $p<0.05$; (**) $p<0.01$; (***) $p<0.001$.

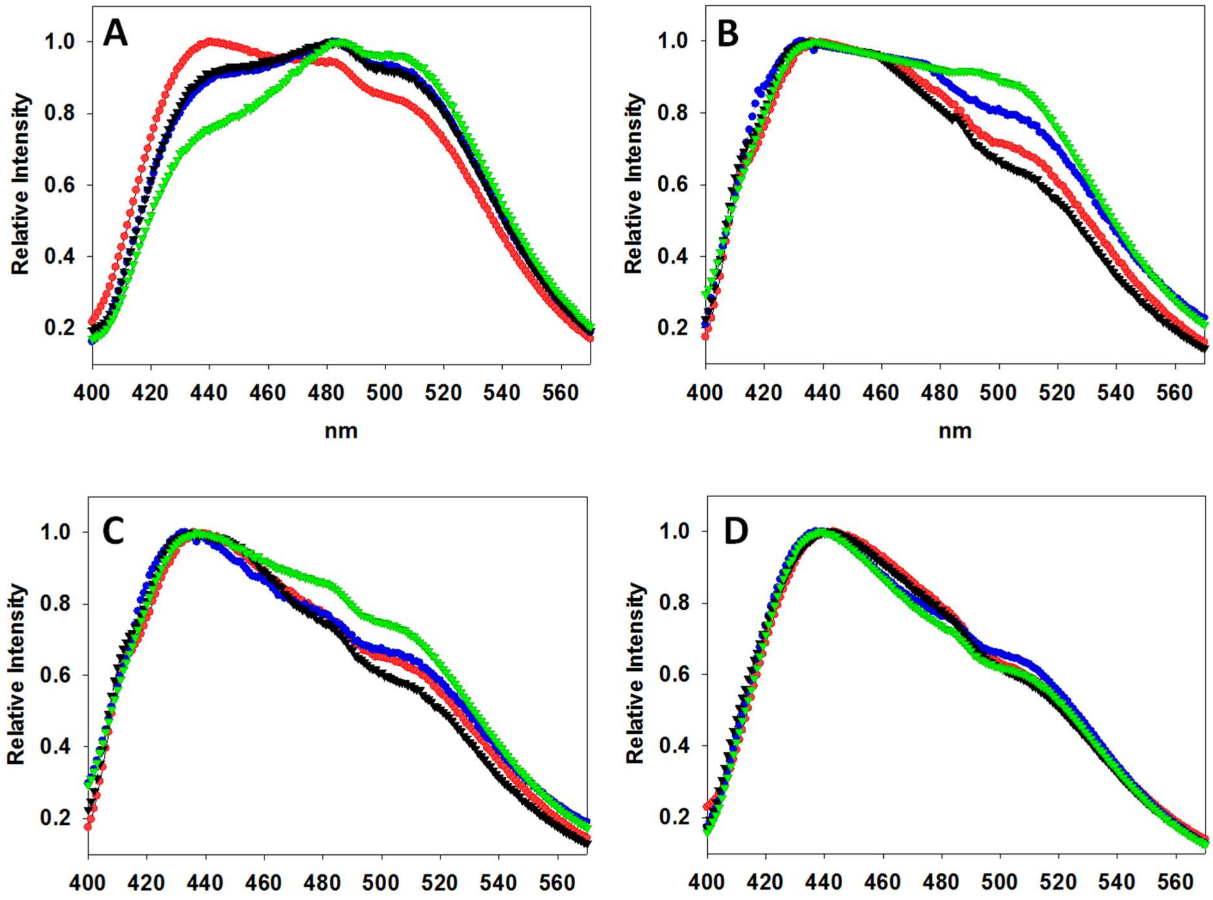


**Figure S8. Laurdan emission spectra at 40°C.** LUV formed from: whole cell lipid extract (**A**)**,** PM lipid extract (**B**), PM patches (**C**) and GPMV (D). Red spectra: SUV formed from CHO (10% FBS) lipid extracts. Blue spectra: SUV formed from CHO (0.04% FBS) lipid extracts. Black spectra: SUV formed from LY-B (10% FBS) lipid extract. Green spectra: SUV formed from LY-B (0.04% FBS) lipid extract.

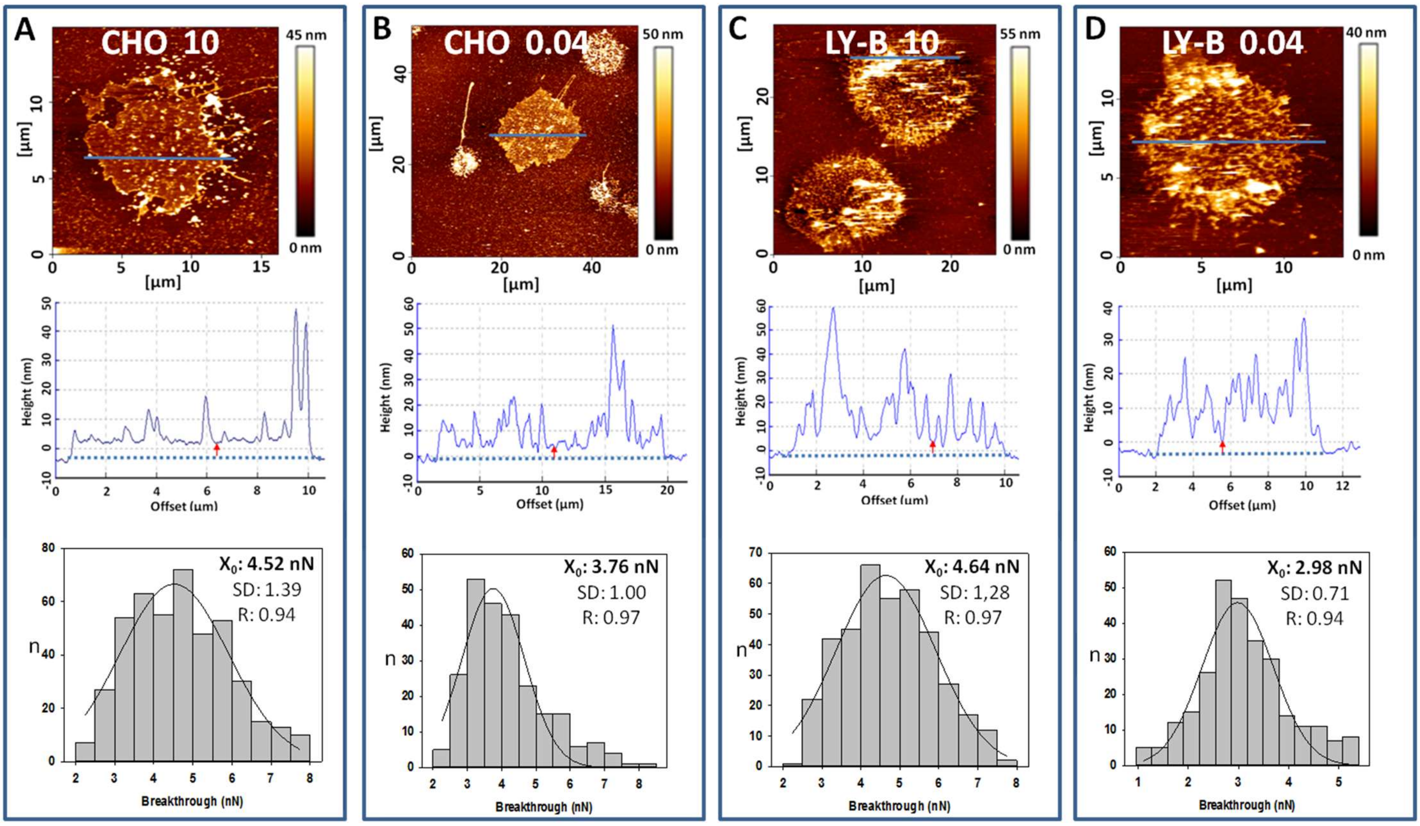


**Figure S9. Topographic image, topographic profile (along the blue line) and breakthrough distributions of PM patches.** PM patches from CHO cells grown in standard (**A**) and SL-deficient medium (**B**), and from LY-B cells grown in standard (**C**) and SL-deficient (**D**) medium.

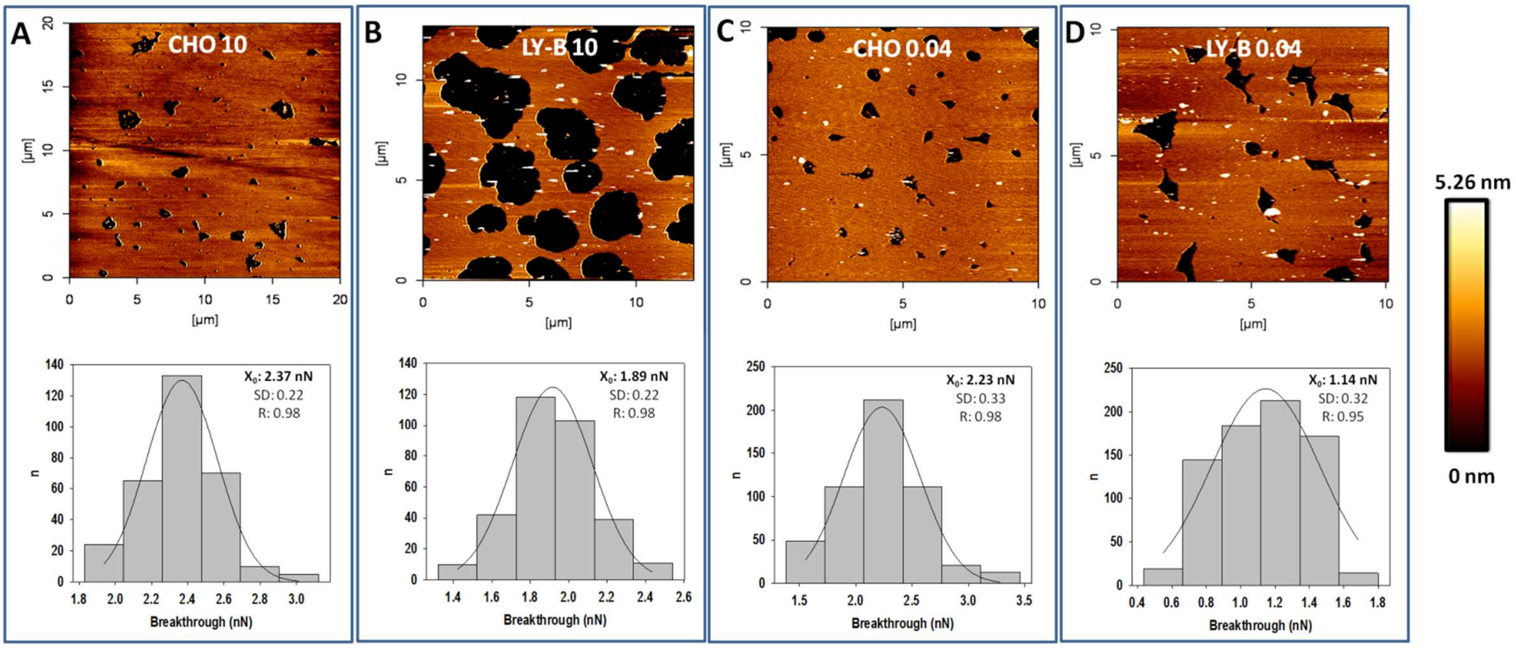


**Figure S10. Topographic images (top) and breakthrough distributions (bottom) of supported planar bilayers formed from whole cell lipid extracts.** CHO (**A**) and LY-B (**B**), cells grown in standard medium. CHO (**C**) and LY-B (**D**) cells grown in SL-deficient medium.

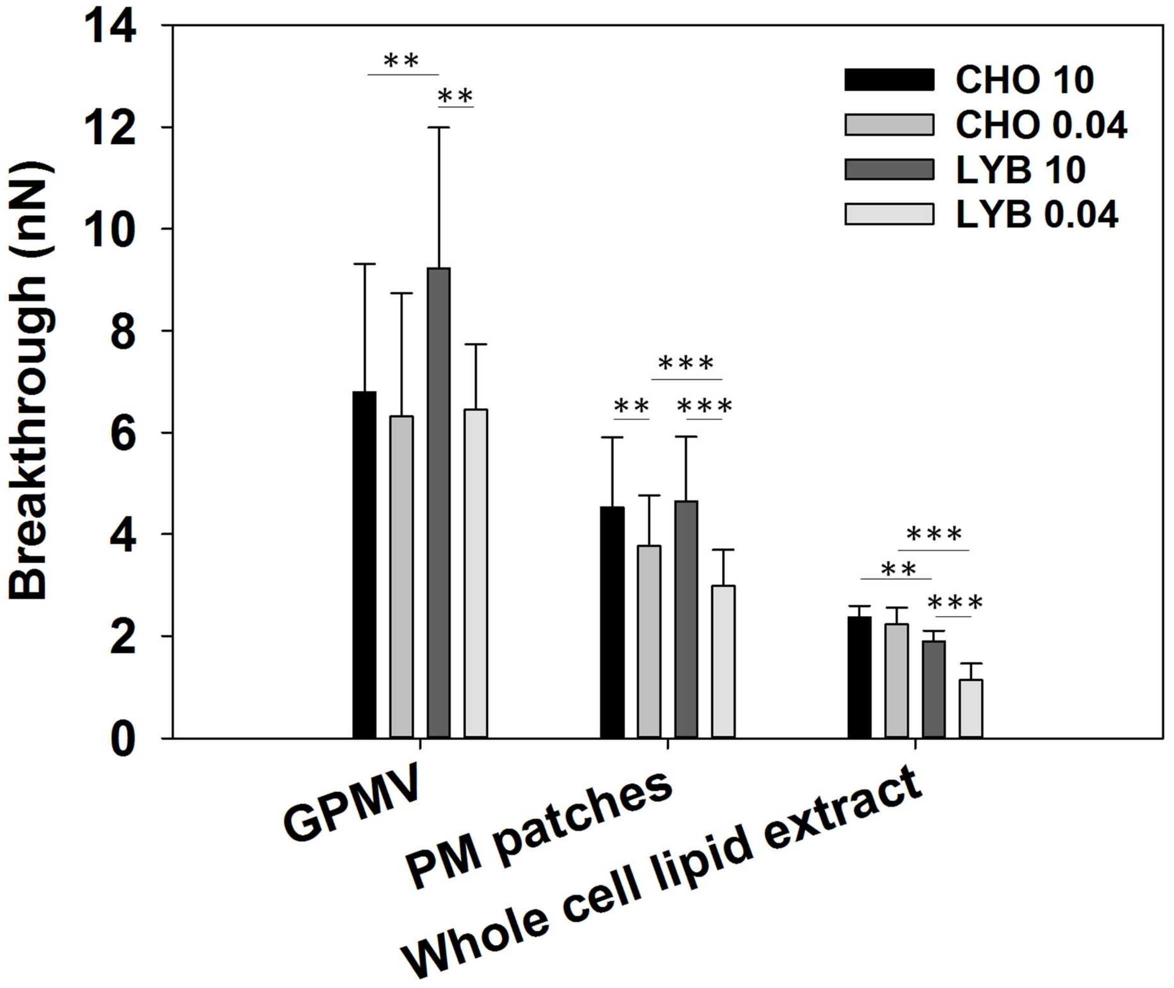


**Figure S11. Topographic breakthrough values of GPMV, PM patches and whole cell lipid extract of CHO ad LY-B cells grown in normal- and deficient- medium.** Mean values ± S.D. (n = 3). Statistical significance was calculated with Student´s t-test: (*) $p<0.05$; (**) $p<0.01$; (***) $p<0.001$.

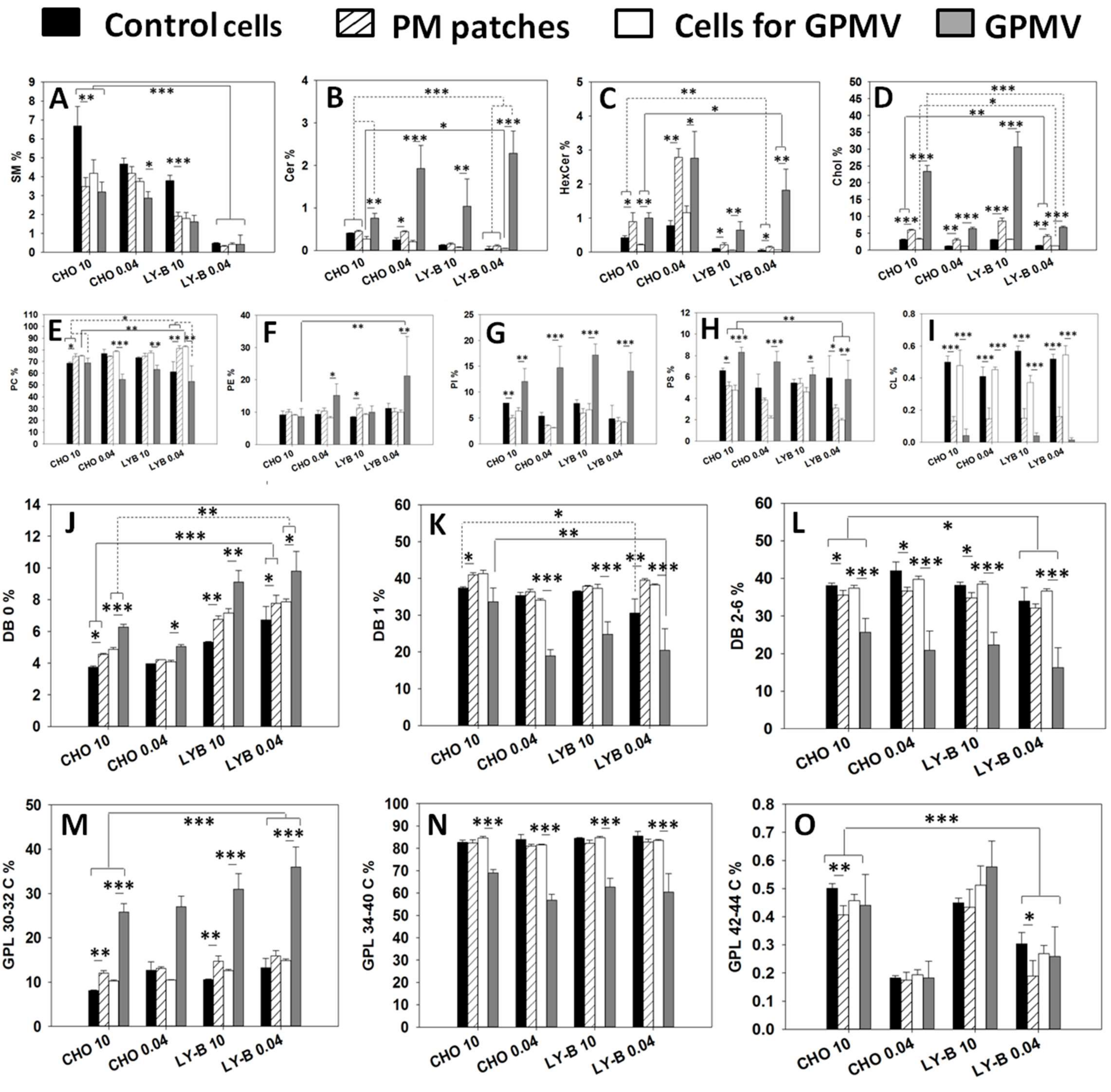


**Figure S12. Lipidomic analysis of whole cells and plasma membrane preparations.** SM (**A**), Cer (**B**), Hex Cer (**C**), Chol (**D**), PC (**E**), PE (**F**), PI (**G**), PS (**H**), CL (**I**). **J-L:** GPL saturation distribution. No double bonds (DB 0) (**J**), one double bond per molecule (DB 1) (**K**), two – six double bonds per molecule (DB 2-6) (**L**). **M-O:** GPL length distribution (number of C atoms in the two acyl chains). 30-32 C (**M**), 34-40 C (**N**), 42-44 C (**O**). Mean values ± S.D. (n = 3). Statistical significance was calculated with ANOVA and Student´s t-test: (*) p<0.05; (**) p<0.01; (***) p<0.001.

**Table S2. SM and Chol contents per cell and per µg protein in the various preparations.** Average values ± S.D. (n = 3).

| Cells (FBS conc.) | µg protein/cell | ng SM/cell | fg SM/µg prot | ng Chol/cell | fg Chol/µg prot |
|---|---|---|---|---|---|
| **CHO (10%)** | 223 ± 50 | 0.25 ± 0.040 | 1.11 ± 0.28 | 0.12 ± 0.01 | 0.52 ± 0.13 |
| **CHO (0.04%)** | 295 ± 44 | 0.17 ± 0.011 | 0.59 ± 0.10 | 0.04 ± 0.0 | 0.14 ± 0.02 |
| **LY-B (10%)** | 225 ± 46 | 0.14 ± 0.012 | 0.62 ± 0.12 | 0.11 ± 0.00 | 0.50 ± 0.10 |
| **LY-B (0.04%)** | 178 ± 39 | 0.02 ± 0.00 | 0.10 ± 0.02 | 0.05 ± 0.00 | 0.27 ± 0.07 |
