## Supplementary material for "CHO/LY-B cell growth under limiting sphingolipid supply: correlation between lipid composition and biophysical properties of sphingolipid-restricted cell membranes^+^": Supp Table 1: Supp Table 1.pdf

| CHO 10 | Control cells | SD Control cells | PM Patches | SD PM patches |
|---|---|---|---|---|
| Total_CER_H2O | 0,41 | 0 | 0,45 | 0,03 |
| Total_HexCER_H2O | 0,42 | 0,06 | 0,9 | 0,26 |
| Total_SM | 6,68 | 1,04 | 3,47 | 0,48 |
| Total_PC | 68,54 | 0,93 | 74,55 | 2,14 |
| Total_PE | 9,2 | 1,2 | 10,11 | 0,73 |
| Total_PI | 7,91 | 0,01 | 5,05 | 0,49 |
| Total_PS | 6,57 | 0,21 | 5,16 | 0,34 |
| Total_CL | 0,5 | 0,04 | 0,13 | 0,03 |
| Total_Chol | 3,12 | 0,17 | 5,91 | 0,29 |
| GPL_short (30-32C) | 8,1 | 0,12 | 12 | 0,58 |
| GPL_long (34-40C) | 82,66 | 0,96 | 82,46 | 1,31 |
| GPL_very_long (42-44C) | 0,5 | 0,02 | 0,41 | 0,03 |
| PC_Lyso | 0,14 | 0,01 | 0,05 | 0,01 |
| PE_Lyso | 0,07 | 0,01 | 0,03 | 0 |
| PI_Lyso | 0,09 | 0,01 | 0,05 | 0,01 |
| PS_Lyso | 0,01 | 0 | 0,02 | 0,01 |
| PC_Ether | 10,78 | 0,23 | 12,43 | 0,31 |
| PE_Ether | 0,23 | 0 | 0,47 | 0,15 |
| PI_Ether | 0,36 | 0,01 | 0,56 | 0,2 |
| PS_Ether | 0,28 | 0,01 | 0,21 | 0,02 |
| CER_H2O | 0,38 | 0,02 | 0,43 | 0,03 |
| DHCER_H2O | 0,02 | 0 | 0,01 | 0 |
| CERP_H2O | 0,01 | 0 | 0 | 0 |
| HexCER_H2O | 0,42 | 0,06 | 0,89 | 0,27 |
| HexDHCER_H2O | 0 | 0 | 0,01 | 0 |
| SM | 6,09 | 1,06 | 3,39 | 0,47 |
| DHSM | 0,59 | 0,02 | 0,08 | 0,01 |
| DB_0 | 3,75 | 0,07 | 4,55 | 0,06 |
| DB_1 | 37,39 | 0,33 | 40,96 | 0,58 |
| DB_2 | 23,62 | 0,44 | 24,99 | 0,88 |
| DB_3 | 3,31 | 0,06 | 2,92 | 0,12 |
| DB_4 | 4,96 | 0,03 | 3,3 | 0,07 |
| DB_5 | 3,77 | 0,07 | 2,42 | 0,09 |
| DB_6 | 2,48 | 0,04 | 1,92 | 0,12 |
| CL_2_DB | 0 | 0 | 0 | 0 |
| CL_3_DB | 0,04 | 0 | 0,02 | 0 |
| CL_4_DB | 0,13 | 0,02 | 0,07 | 0,01 |
| CL_5_DB | 0,2 | 0,01 | 0,09 | 0,01 |
| CL_6_DB | 0,1 | 0,01 | 0,04 | 0 |
| CL_7_DB | 0,02 | 0 | 0,01 | 0 |
| PC_UI_0 | 4,09 | 0,05 | 5,15 | 0,1 |
| PC_UI_1 | 35,24 | 0,58 | 40,56 | 0,62 |
| PC_UI_2 | 21,89 | 0,47 | 23,33 | 1,27 |
| PC_UI_3 | 2,55 | 0,07 | 2,31 | 0,18 |
| PC_UI_4 | 1,85 | 0,02 | 1,28 | 0,11 |
| PC_UI_5 | 1,74 | 0,02 | 1,07 | 0,09 |
| PC_UI_6 | 1,21 | 0,02 | 0,84 | 0,06 |

| | | | | |
|---|---|---|---|---|
| PE_UI_0 | 0,21 | 0,01 | 0,26 | 0,03 |
| PE_UI_1 | 1,89 | 0,02 | 2,96 | 0,24 |
| PE_UI_2 | 1,92 | 0,05 | 2,81 | 0,19 |
| PE_UI_3 | 0,4 | 0,01 | 0,56 | 0,05 |
| PE_UI_4 | 1,1 | 0,01 | 1,32 | 0,17 |
| PE_UI_5 | 1,39 | 0,04 | 1,21 | 0,06 |
| PE_UI_6 | 1,19 | 0,03 | 1 | 0,01 |
| PI_UI_0 | 0,24 | 0 | 0,12 | 0 |
| PI_UI_1 | 2,16 | 0,07 | 1,2 | 0,05 |
| PI_UI_2 | 1,75 | 0,04 | 1,46 | 0,21 |
| PI_UI_3 | 0,74 | 0,01 | 0,5 | 0,04 |
| PI_UI_4 | 2,28 | 0,07 | 1,21 | 0,04 |
| PI_UI_5 | 0,57 | 0,02 | 0,3 | 0,06 |
| PI_UI_6 | 0,17 | 0,01 | 0,26 | 0,11 |
| PS_UI_0 | 0,31 | 0,01 | 0,21 | 0 |
| PS_UI_1 | 3,8 | 0,09 | 3,2 | 0,27 |
| PS_UI_2 | 0,57 | 0,02 | 0,52 | 0,02 |
| PS_UI_3 | 0,2 | 0,01 | 0,18 | 0,02 |
| PS_UI_4 | 0,42 | 0,02 | 0,25 | 0,01 |
| PS_UI_5 | 0,84 | 0,06 | 0,47 | 0,03 |
| PS_UI_6 | 0,48 | 0,02 | 0,33 | 0,01 |
| PC_short | 7,78 | 0,13 | 11,56 | 0,48 |
| PC_long | 60,37 | 0,96 | 62,85 | 2,6 |
| PC_very_long | 0,16 | 0,01 | 0,14 | 0,03 |
| PE_short | 0,15 | 0 | 0,23 | 0,04 |
| PE_long | 7,93 | 0,13 | 9,81 | 0,69 |
| PE_very_long | 0,07 | 0 | 0,07 | 0 |
| PI_short | 0,09 | 0 | 0,11 | 0,02 |
| PI_long | 7,73 | 0,01 | 4,84 | 0,46 |
| PI_very_long | 0,09 | 0 | 0,1 | 0 |
| PS_short | 0,08 | 0 | 0,11 | 0,04 |
| PS_long | 6,29 | 0,2 | 4,96 | 0,31 |
| PS_very_long | 0,19 | 0,01 | 0,09 | 0,01 |
| CER_H2O_10 | 0 | 0 | 0 | 0 |
| CER_H2O_14 | 0 | 0 | 0 | 0 |
| CER_H2O_16 | 0,19 | 0,01 | 0,27 | 0,01 |
| CER_H2O_18 | 0,01 | 0 | 0,01 | 0 |
| CER_H2O_20 | 0 | 0 | 0 | 0 |
| CER_H2O_22 | 0,06 | 0,01 | 0,02 | 0 |
| CER_H2O_24 | 0,16 | 0,01 | 0,14 | 0,02 |
| CER_H2O_26 | 0 | 0 | 0,01 | 0 |
| HexCER_H2O_10 | 0 | 0 | 0 | 0 |
| HexCER_H2O_12 | 0,01 | 0 | 0 | 0 |
| HexCER_H2O_14 | 0 | 0 | 0 | 0 |
| HexCER_H2O_16 | 0,27 | 0,04 | 0,6 | 0,16 |
| HexCER_H2O_18 | 0 | 0 | 0,01 | 0 |
| HexCER_H2O_20 | 0 | 0 | 0 | 0 |
| HexCER_H2O_22 | 0,02 | 0,01 | 0,05 | 0,02 |
| HexCER_H2O_24 | 0,12 | 0,01 | 0,23 | 0,08 |
| HexCER_H2O_26 | 0 | 0 | 0 | 0 |

| | | | | |
|---|---|---|---|---|
| SM_14 | 0,05 | 0,01 | 0,03 | 0 |
| SM_16 | 5,32 | 1,22 | 3,23 | 0,43 |
| SM_18 | 0,12 | 0,02 | 0,01 | 0 |
| SM_20 | 0,13 | 0,03 | 0 | 0 |
| SM_22 | 0,52 | 0,12 | 0,02 | 0 |
| SM_24 | 0,54 | 0,05 | 0,18 | 0,05 |
| PC_28 | 0,07 | 0 | 0,16 | 0,02 |
| PC_30 | 1,1 | 0,08 | 2,94 | 0,37 |
| PC_32 | 6,61 | 0,07 | 8,44 | 0,09 |
| PC_34 | 30,83 | 0,55 | 33,28 | 0,99 |
| PC_36 | 23,82 | 0,4 | 24,46 | 1,4 |
| PC_38 | 4,67 | 0,05 | 4,36 | 0,21 |
| PC_40 | 1 | 0,02 | 0,71 | 0,07 |
| PC_42 | 0,13 | 0,01 | 0,11 | 0,02 |
| PC_44 | 0,03 | 0 | 0,03 | 0,01 |
| PE_28 | 0 | 0 | 0 | 0 |
| PE_30 | 0,01 | 0 | 0,03 | 0,01 |
| PE_32 | 0,13 | 0 | 0,19 | 0,03 |
| PE_34 | 1,12 | 0,02 | 1,84 | 0,21 |
| PE_36 | 3 | 0,05 | 4,4 | 0,35 |
| PE_38 | 1,76 | 0,02 | 1,83 | 0,09 |
| PE_40 | 1,96 | 0,06 | 1,71 | 0,07 |
| PE_42 | 0,05 | 0 | 0,06 | 0 |
| PE_44 | 0,01 | 0 | 0,01 | 0 |
| PI_28 | 0 | 0 | 0 | 0 |
| PI_30 | 0,01 | 0 | 0,03 | 0,01 |
| PI_32 | 0,08 | 0 | 0,07 | 0,01 |
| PI_34 | 1,16 | 0,02 | 0,54 | 0,04 |
| PI_36 | 2,89 | 0,1 | 1,9 | 0,21 |
| PI_38 | 3,04 | 0,09 | 2,04 | 0,24 |
| PI_40 | 0,55 | 0,02 | 0,33 | 0,06 |
| PI_42 | 0,06 | 0 | 0,07 | 0 |
| PI_44 | 0,02 | 0 | 0,03 | 0 |
| PS_28 | 0 | 0 | 0 | 0 |
| PS_30 | 0 | 0 | 0,02 | 0,02 |
| PS_32 | 0,08 | 0 | 0,06 | 0,02 |
| PS_34 | 0,56 | 0,02 | 0,53 | 0,02 |
| PS_36 | 3,6 | 0,08 | 3,06 | 0,24 |
| PS_38 | 0,46 | 0,03 | 0,35 | 0,01 |
| PS_40 | 1,68 | 0,09 | 1,01 | 0,05 |
| PS_42 | 0,18 | 0,01 | 0,09 | 0,01 |
| PS_44 | 0,01 | 0 | 0 | 0 |
| CL_68 | 0,06 | 0,01 | 0,03 | 0 |
| CL_70 | 0,21 | 0,02 | 0,11 | 0,02 |
| CL_72 | 0,21 | 0,02 | 0,09 | 0,01 |
| CL_74 | 0,02 | 0 | 0,01 | 0 |
| CL_76 | 0 | 0 | 0 | 0 |
| CL_78 | 0 | 0 | 0 | 0 |
| PC_Lyso_short | 0,07 | 0,01 | 0,02 | 0 |
| PC_Lyso_long | 0,07 | 0 | 0,03 | 0,01 |

| | | | | |
|---|---|---|---|---|
| PE_Lyso_short | 0,01 | 0 | 0 | 0 |
| PE_Lyso_long | 0,06 | 0,01 | 0,03 | 0 |
| PI_Lyso_short | 0,01 | 0 | 0,01 | 0 |
| PI_Lyso_long | 0,08 | 0,01 | 0,04 | 0,01 |
| PS_Lyso_short | 0 | 0 | 0,02 | 0,01 |
| PS_Lyso_long | 0 | 0 | 0 | 0 |
| PC_Lyso_UI_0 | 0,09 | 0 | 0,03 | 0 |
| PC_Lyso_UI_1 | 0,05 | 0 | 0,02 | 0 |
| PC_Lyso_UI_2 | 0 | 0 | 0 | 0 |
| PE_Lyso_UI_0 | 0,03 | 0 | 0,01 | 0 |
| PE_Lyso_UI_1 | 0,04 | 0,01 | 0,02 | 0 |
| PE_Lyso_UI_2 | 0 | 0 | 0 | 0 |
| PI_Lyso_UI_0 | 0,07 | 0,01 | 0,03 | 0,01 |
| PI_Lyso_UI_1 | 0,02 | 0 | 0,01 | 0 |
| PI_Lyso_UI_2 | 0 | 0 | 0 | 0 |
| PS_Lyso_UI_0 | 0 | 0 | 0 | 0 |
| PS_Lyso_UI_1 | 0 | 0 | 0,02 | 0,01 |
| PC_Ether_short | 0,9 | 0,04 | 1,8 | 0,27 |
| PC_Ether_long | 9,8 | 0,24 | 10,57 | 0,54 |
| PC_Ether_very_long | 0,07 | 0,01 | 0,06 | 0,02 |
| PE_Ether_short | 0,02 | 0 | 0,02 | 0,01 |
| PE_Ether_long | 0,21 | 0 | 0,45 | 0,14 |
| PI_Ether_short | 0,02 | 0 | 0,01 | 0,01 |
| PI_Ether_long | 0,33 | 0,01 | 0,53 | 0,19 |
| PI_Ether_very_long | 0,01 | 0 | 0,01 | 0 |
| PS_Ether_short | 0,06 | 0 | 0,01 | 0,01 |
| PS_Ether_long | 0,2 | 0,02 | 0,19 | 0,02 |
| PS_Ether_very_long | 0,01 | 0 | 0 | 0 |
| PC_Ether_UI_0 | 0,79 | 0,01 | 1,09 | 0,03 |
| PC_Ether_UI_1 | 5,47 | 0,17 | 6,63 | 0,12 |
| PC_Ether_UI_2 | 2,47 | 0,1 | 2,68 | 0,19 |
| PC_Ether_UI_3 | 0,52 | 0,02 | 0,53 | 0,07 |
| PC_Ether_UI_4 | 0,58 | 0,01 | 0,53 | 0,07 |
| PC_Ether_UI_5 | 0,66 | 0,02 | 0,53 | 0,05 |
| PC_Ether_UI_6 | 0,47 | 0,01 | 0,44 | 0,04 |
| PE_Ether_UI_0 | 0,02 | 0 | 0,01 | 0 |
| PE_Ether_UI_1 | 0,03 | 0 | 0,06 | 0,01 |
| PE_Ether_UI_2 | 0,01 | 0 | 0,03 | 0,01 |
| PE_Ether_UI_3 | 0,01 | 0 | 0,04 | 0,01 |
| PE_Ether_UI_4 | 0,04 | 0 | 0,21 | 0,12 |
| PE_Ether_UI_5 | 0,04 | 0 | 0,06 | 0,01 |
| PE_Ether_UI_6 | 0,07 | 0 | 0,07 | 0 |
| PI_Ether_UI_0 | 0,03 | 0 | 0 | 0 |
| PI_Ether_UI_1 | 0,09 | 0 | 0,07 | 0,01 |
| PI_Ether_UI_2 | 0,11 | 0 | 0,4 | 0,17 |
| PI_Ether_UI_3 | 0,04 | 0 | 0,03 | 0,01 |
| PI_Ether_UI_4 | 0,05 | 0 | 0,02 | 0 |
| PI_Ether_UI_5 | 0,04 | 0 | 0,03 | 0,01 |
| PI_Ether_UI_6 | 0 | 0 | 0,01 | 0 |
| PS_Ether_UI_0 | 0,07 | 0 | 0,01 | 0 |

| | | | | |
|---|---|---|---|---|
| PS_Ether_UI_1 | 0,13 | 0,01 | 0,12 | 0,01 |
| PS_Ether_UI_2 | 0,03 | 0 | 0,02 | 0 |
| PS_Ether_UI_3 | 0 | 0 | 0,04 | 0,01 |
| PS_Ether_UI_5 | 0,02 | 0 | 0,01 | 0 |
| PS_Ether_UI_6 | 0,01 | 0 | 0,01 | 0 |
| PC_Ether_30 | 0,09 | 0 | 0,18 | 0,01 |
| PC_Ether_32 | 0,81 | 0,04 | 1,62 | 0,26 |
| PC_Ether_34 | 4,72 | 0,22 | 5,27 | 0,12 |
| PC_Ether_36 | 3,49 | 0,12 | 3,53 | 0,28 |
| PC_Ether_38 | 1,49 | 0,04 | 1,46 | 0,11 |
| PC_Ether_40 | 0,37 | 0,01 | 0,31 | 0,05 |
| PC_Ether_42 | 0,06 | 0 | 0,05 | 0,02 |
| PC_Ether_44 | 0,01 | 0 | 0,01 | 0 |
| PE_Ether_30 | 0 | 0 | 0,01 | 0 |
| PE_Ether_32 | 0,02 | 0 | 0,01 | 0 |
| PE_Ether_34 | 0,05 | 0 | 0,25 | 0,13 |
| PE_Ether_36 | 0,03 | 0 | 0,06 | 0,01 |
| PE_Ether_38 | 0,09 | 0 | 0,1 | 0,01 |
| PE_Ether_40 | 0,04 | 0 | 0,03 | 0 |
| PI_Ether_30 | 0 | 0 | 0,01 | 0,01 |
| PI_Ether_32 | 0,02 | 0 | 0 | 0 |
| PI_Ether_34 | 0 | 0 | 0 | 0 |
| PI_Ether_36 | 0,14 | 0 | 0,07 | 0 |
| PI_Ether_38 | 0,14 | 0,01 | 0,45 | 0,19 |
| PI_Ether_40 | 0,04 | 0 | 0,01 | 0 |
| PI_Ether_42 | 0,01 | 0 | 0,01 | 0 |
| PI_Ether_44 | 0 | 0 | 0 | 0 |
| PS_Ether_30 | 0 | 0 | 0,01 | 0 |
| PS_Ether_32 | 0,06 | 0,01 | 0 | 0 |
| PS_Ether_34 | 0,02 | 0 | 0,06 | 0,01 |
| PS_Ether_36 | 0,11 | 0,01 | 0,09 | 0 |
| PS_Ether_38 | 0,05 | 0,01 | 0,04 | 0 |
| PS_Ether_40 | 0,02 | 0 | 0,01 | 0 |

| Cells for GPMV | SD cells for GPMV | GPMV | SD GPMV | CHO 0.04 |
|---|---|---|---|---|
| 0,27 | 0,06 | 0,76 | 0,11 | Total_CER_H2O |
| 0,21 | 0,02 | 1 | 0,16 | Total_HexCER_H2O |
| 4,18 | 0,72 | 3,19 | 0,53 | Total_SM |
| 74,91 | 0,48 | 68,79 | 4,18 | Total_PC |
| 9,08 | 0,24 | 8,69 | 2,43 | Total_PE |
| 6,37 | 0,62 | 12,07 | 2,49 | Total_PI |
| 4,76 | 0,46 | 8,32 | 0,48 | Total_PS |
| 0,48 | 0,1 | 0,04 | 0,04 | Total_CL |
| 3,29 | 0,2 | 23,4 | 1,77 | Total_Chol |
| 10,3 | 0,21 | 25,79 | 1,91 | GPL_short (30-32C) |
| 84,63 | 0,77 | 68,98 | 1,55 | GPL_long (34-40C) |
| 0,46 | 0,02 | 0,44 | 0,11 | GPL_very_long (42-44C) |
| 0,08 | 0 | 0,27 | 0,02 | PC_Lyso |
| 0,07 | 0 | 0,13 | 0,04 | PE_Lyso |
| 0,04 | 0 | 0,21 | 0,08 | PI_Lyso |
| 0,01 | 0 | 0,46 | 0,23 | PS_Lyso |
| 11 | 0,24 | 17,65 | 1,46 | PC_Ether |
| 0,23 | 0,01 | 4,75 | 2,17 | PE_Ether |
| 0,25 | 0,04 | 5,11 | 1,1 | PI_Ether |
| 0,15 | 0,01 | 1,06 | 0,38 | PS_Ether |
| 0,25 | 0,06 | 0,69 | 0,1 | CER_H2O |
| 0,02 | 0,01 | 0,05 | 0,01 | DHCER_H2O |
| 0 | 0 | 0,02 | 0 | CERP_H2O |
| 0,21 | 0,02 | 0,98 | 0,16 | HexCER_H2O |
| 0 | 0 | 0,02 | 0 | HexDHCER_H2O |
| 4,09 | 0,7 | 3,07 | 0,52 | SM |
| 0,09 | 0,01 | 0,13 | 0,01 | DHSM |
| 4,87 | 0,12 | 6,28 | 0,18 | DB_0 |
| 41,31 | 0,94 | 33,65 | 3,76 | DB_1 |
| 23,42 | 0,05 | 14,12 | 1,71 | DB_2 |
| 3,41 | 0,15 | 2,29 | 0,36 | DB_3 |
| 4,7 | 0,21 | 1,79 | 0,15 | DB_4 |
| 3,39 | 0,19 | 2,14 | 0,19 | DB_5 |
| 2,46 | 0,17 | 5,29 | 1,34 | DB_6 |
| 0,06 | 0,06 | 0 | 0 | CL_2_DB |
| 0,04 | 0 | 0 | 0 | CL_3_DB |
| 0,11 | 0,02 | 0 | 0 | CL_4_DB |
| 0,13 | 0,01 | 0 | 0,01 | CL_5_DB |
| 0,1 | 0,01 | 0,02 | 0,01 | CL_6_DB |
| 0,02 | 0 | 0 | 0 | CL_7_DB |
| 5,38 | 0,13 | 7,47 | 0,12 | PC_UI_0 |
| 40,08 | 0,84 | 42,47 | 2,02 | PC_UI_1 |
| 21,89 | 0,19 | 14 | 1,73 | PC_UI_2 |
| 2,68 | 0,12 | 1,74 | 0,24 | PC_UI_3 |
| 2,01 | 0,13 | 1,18 | 0,15 | PC_UI_4 |
| 1,65 | 0,13 | 0,95 | 0,13 | PC_UI_5 |
| 1,22 | 0,12 | 0,97 | 0,08 | PC_UI_6 |

| | | | | |
|---|---|---|---|---|
| 0,21 | 0,01 | 0,43 | 0,07 | PE_UI_0 |
| 2,23 | 0,05 | 1,69 | 0,2 | PE_UI_1 |
| 2,2 | 0,05 | 1,13 | 0,21 | PE_UI_2 |
| 0,5 | 0,03 | 0,53 | 0,16 | PE_UI_3 |
| 1,23 | 0,04 | 3,12 | 1,16 | PE_UI_4 |
| 1,42 | 0,07 | 0,39 | 0,08 | PE_UI_5 |
| 1,29 | 0,07 | 0,39 | 0,06 | PE_UI_6 |
| 0,16 | 0,02 | 0,28 | 0,03 | PI_UI_0 |
| 1,79 | 0,32 | 1,13 | 0,34 | PI_UI_1 |
| 1,43 | 0,15 | 3,65 | 0,21 | PI_UI_2 |
| 0,63 | 0,04 | 0,66 | 0,08 | PI_UI_3 |
| 1,81 | 0,09 | 0,43 | 0,08 | PI_UI_4 |
| 0,41 | 0,02 | 1,1 | 0,3 | PI_UI_5 |
| 0,14 | 0,01 | 2,24 | 0,49 | PI_UI_6 |
| 0,18 | 0,02 | 0,3 | 0,06 | PS_UI_0 |
| 2,95 | 0,29 | 3 | 0,16 | PS_UI_1 |
| 0,43 | 0,04 | 0,23 | 0,03 | PS_UI_2 |
| 0,17 | 0,01 | 0,82 | 0,34 | PS_UI_3 |
| 0,33 | 0,04 | 0,24 | 0,06 | PS_UI_4 |
| 0,6 | 0,05 | 0,36 | 0,09 | PS_UI_5 |
| 0,37 | 0,03 | 0,69 | 0,1 | PS_UI_6 |
| 10,03 | 0,22 | 22,12 | 1,34 | PC_short |
| 64,71 | 0,52 | 46,53 | 5,3 | PC_long |
| 0,17 | 0,01 | 0,14 | 0,03 | PC_very_long |
| 0,15 | 0,01 | 1,24 | 0,23 | PE_short |
| 8,85 | 0,24 | 7,45 | 2,21 | PE_long |
| 0,08 | 0 | 0 | 0 | PE_very_long |
| 0,1 | 0,01 | 0,77 | 0,05 | PI_short |
| 6,16 | 0,59 | 11,02 | 2,31 | PI_long |
| 0,11 | 0,01 | 0,28 | 0,16 | PI_very_long |
| 0,03 | 0 | 1,65 | 0,46 | PS_short |
| 4,9 | 0,44 | 3,98 | 0,17 | PS_long |
| 0,1 | 0,02 | 0,01 | 0,02 | PS_very_long |
| 0 | 0 | 0,01 | 0 | CER_H2O_10 |
| 0 | 0 | 0,01 | 0,01 | CER_H2O_14 |
| 0,13 | 0,03 | 0,21 | 0,02 | CER_H2O_16 |
| 0,01 | 0 | 0,09 | 0,03 | CER_H2O_18 |
| 0 | 0 | 0,04 | 0,01 | CER_H2O_20 |
| 0,01 | 0 | 0,05 | 0,01 | CER_H2O_22 |
| 0,11 | 0,03 | 0,25 | 0,03 | CER_H2O_24 |
| 0 | 0 | 0,1 | 0,03 | CER_H2O_26 |
| 0 | 0 | 0,03 | 0,02 | HexCER_H2O_10 |
| 0 | 0 | 0,02 | 0,01 | HexCER_H2O_12 |
| 0 | 0 | 0,03 | 0,02 | HexCER_H2O_14 |
| 0,13 | 0,01 | 0,61 | 0,1 | HexCER_H2O_16 |
| 0 | 0 | 0,03 | 0 | HexCER_H2O_18 |
| 0 | 0 | 0,01 | 0,01 | HexCER_H2O_20 |
| 0,01 | 0 | 0,05 | 0 | HexCER_H2O_22 |
| 0,07 | 0,01 | 0,2 | 0,05 | HexCER_H2O_24 |
| 0 | 0 | 0,01 | 0 | HexCER_H2O_26 |

| | | | | |
|---|---|---|---|---|
| 0,04 | 0 | 0,03 | 0 | SM_14 |
| 3,78 | 0,7 | 2,88 | 0,48 | SM_16 |
| 0,01 | 0,01 | 0,01 | 0 | SM_18 |
| 0 | 0 | 0 | 0 | SM_20 |
| 0,03 | 0 | 0,08 | 0 | SM_22 |
| 0,32 | 0,03 | 0,18 | 0,04 | SM_24 |
| 0,14 | 0,02 | 0,42 | 0,04 | PC_28 |
| 1,83 | 0,07 | 4,97 | 0,4 | PC_30 |
| 8,03 | 0,14 | 16,65 | 1,55 | PC_32 |
| 35,53 | 1,26 | 26,39 | 3 | PC_34 |
| 23,35 | 0,45 | 15,67 | 1,9 | PC_36 |
| 4,78 | 0,34 | 3,64 | 0,53 | PC_38 |
| 1 | 0,08 | 0,64 | 0,06 | PC_40 |
| 0,14 | 0,01 | 0,1 | 0,02 | PC_42 |
| 0,03 | 0 | 0,04 | 0,01 | PC_44 |
| 0 | 0 | 0,04 | 0,01 | PE_28 |
| 0,01 | 0 | 0,44 | 0,1 | PE_30 |
| 0,13 | 0 | 0,69 | 0,1 | PE_32 |
| 1,34 | 0,04 | 4,75 | 1,96 | PE_34 |
| 3,38 | 0,05 | 1,73 | 0,39 | PE_36 |
| 1,92 | 0,1 | 0,52 | 0,14 | PE_38 |
| 2,15 | 0,1 | 0,39 | 0,13 | PE_40 |
| 0,07 | 0 | 0 | 0 | PE_42 |
| 0,01 | 0 | 0 | 0 | PE_44 |
| 0 | 0 | 0,03 | 0,03 | PI_28 |
| 0,02 | 0 | 0,25 | 0,08 | PI_30 |
| 0,07 | 0,01 | 0,33 | 0,06 | PI_32 |
| 0,94 | 0,17 | 0,28 | 0,18 | PI_34 |
| 2,28 | 0,31 | 4,23 | 1,12 | PI_36 |
| 2,44 | 0,12 | 5,06 | 0,86 | PI_38 |
| 0,46 | 0,03 | 1,4 | 0,48 | PI_40 |
| 0,08 | 0,01 | 0,12 | 0,09 | PI_42 |
| 0,03 | 0 | 0,16 | 0,1 | PI_44 |
| 0 | 0 | 0 | 0 | PS_28 |
| 0 | 0 | 0,35 | 0,07 | PS_30 |
| 0,02 | 0 | 0,84 | 0,17 | PS_32 |
| 0,52 | 0,05 | 1,12 | 0,27 | PS_34 |
| 2,77 | 0,26 | 1,82 | 0,18 | PS_36 |
| 0,36 | 0,03 | 0,2 | 0,03 | PS_38 |
| 1,24 | 0,12 | 0,84 | 0,11 | PS_40 |
| 0,09 | 0,02 | 0,01 | 0,02 | PS_42 |
| 0,01 | 0 | 0 | 0 | PS_44 |
| 0,05 | 0,01 | 0 | 0 | CL_68 |
| 0,23 | 0,07 | 0 | 0 | CL_70 |
| 0,17 | 0,01 | 0 | 0 | CL_72 |
| 0,02 | 0,01 | 0,02 | 0,01 | CL_74 |
| 0 | 0 | 0,02 | 0,03 | CL_76 |
| 0 | 0 | 0 | 0 | CL_78 |
| 0,03 | 0 | 0,09 | 0,01 | PC_Lyso_short |
| 0,05 | 0 | 0,19 | 0,02 | PC_Lyso_long |

| | | | | |
|---|---|---|---|---|
| 0,01 | 0 | 0,07 | 0,04 | PE_Lyso_short |
| 0,06 | 0 | 0,06 | 0,03 | PE_Lyso_long |
| 0 | 0 | 0,16 | 0,08 | PI_Lyso_short |
| 0,04 | 0 | 0,05 | 0,03 | PI_Lyso_long |
| 0 | 0 | 0,46 | 0,23 | PS_Lyso_short |
| 0 | 0 | 0 | 0 | PS_Lyso_long |
| 0,05 | 0 | 0,2 | 0,01 | PC_Lyso_UI_0 |
| 0,04 | 0 | 0,04 | 0,01 | PC_Lyso_UI_1 |
| 0 | 0 | 0,04 | 0,01 | PC_Lyso_UI_2 |
| 0,03 | 0 | 0,01 | 0,01 | PE_Lyso_UI_0 |
| 0,04 | 0 | 0,09 | 0,05 | PE_Lyso_UI_1 |
| 0 | 0 | 0,02 | 0,01 | PE_Lyso_UI_2 |
| 0,03 | 0 | 0,05 | 0,03 | PI_Lyso_UI_0 |
| 0,01 | 0 | 0,16 | 0,08 | PI_Lyso_UI_1 |
| 0 | 0 | 0 | 0 | PI_Lyso_UI_2 |
| 0 | 0 | 0 | 0 | PS_Lyso_UI_0 |
| 0 | 0 | 0,46 | 0,23 | PS_Lyso_UI_1 |
| 1,1 | 0,03 | 10,38 | 2,01 | PC_Ether_short |
| 9,83 | 0,22 | 7,22 | 0,74 | PC_Ether_long |
| 0,07 | 0 | 0,05 | 0,01 | PC_Ether_very_long |
| 0 | 0 | 0,32 | 0,06 | PE_Ether_short |
| 0,23 | 0,01 | 4,43 | 2,12 | PE_Ether_long |
| 0,01 | 0 | 0,14 | 0,03 | PI_Ether_short |
| 0,23 | 0,04 | 4,86 | 1,04 | PI_Ether_long |
| 0,01 | 0 | 0,11 | 0,09 | PI_Ether_very_long |
| 0 | 0 | 0,11 | 0,03 | PS_Ether_short |
| 0,14 | 0,01 | 0,95 | 0,36 | PS_Ether_long |
| 0,01 | 0 | 0 | 0 | PS_Ether_very_long |
| 0,95 | 0,04 | 1,65 | 0,02 | PC_Ether_UI_0 |
| 5,45 | 0,15 | 13,2 | 1,58 | PC_Ether_UI_1 |
| 2,39 | 0,09 | 1,5 | 0,19 | PC_Ether_UI_2 |
| 0,52 | 0,02 | 0,31 | 0,04 | PC_Ether_UI_3 |
| 0,61 | 0,02 | 0,31 | 0,05 | PC_Ether_UI_4 |
| 0,61 | 0,03 | 0,31 | 0,05 | PC_Ether_UI_5 |
| 0,48 | 0,03 | 0,37 | 0,04 | PC_Ether_UI_6 |
| 0 | 0 | 0,17 | 0,04 | PE_Ether_UI_0 |
| 0,04 | 0 | 0,19 | 0,02 | PE_Ether_UI_1 |
| 0,01 | 0 | 0,13 | 0,04 | PE_Ether_UI_2 |
| 0,01 | 0 | 0,28 | 0,1 | PE_Ether_UI_3 |
| 0,04 | 0 | 3,86 | 1,95 | PE_Ether_UI_4 |
| 0,04 | 0 | 0,11 | 0,05 | PE_Ether_UI_5 |
| 0,07 | 0 | 0,01 | 0,01 | PE_Ether_UI_6 |
| 0 | 0 | 0,05 | 0,01 | PI_Ether_UI_0 |
| 0,06 | 0,01 | 0,33 | 0,09 | PI_Ether_UI_1 |
| 0,1 | 0,01 | 4,33 | 1,02 | PI_Ether_UI_2 |
| 0,03 | 0,01 | 0,11 | 0,04 | PI_Ether_UI_3 |
| 0,03 | 0 | 0,03 | 0,01 | PI_Ether_UI_4 |
| 0,02 | 0 | 0,21 | 0,07 | PI_Ether_UI_5 |
| 0,01 | 0 | 0,04 | 0,01 | PI_Ether_UI_6 |
| 0,01 | 0 | 0,08 | 0,02 | PS_Ether_UI_0 |

| | | | | |
|---|---|---|---|---|
| 0,09 | 0,01 | 0,17 | 0,02 | PS_Ether_UI_1 |
| 0,02 | 0 | 0,03 | 0,02 | PS_Ether_UI_2 |
| 0 | 0 | 0,76 | 0,35 | PS_Ether_UI_3 |
| 0,01 | 0 | 0,01 | 0 | PS_Ether_UI_5 |
| 0,01 | 0 | 0,02 | 0,01 | PS_Ether_UI_6 |
| 0,15 | 0,01 | 0,35 | 0,01 | PC_Ether_30 |
| 0,95 | 0,04 | 10,04 | 2 | PC_Ether_32 |
| 4,66 | 0,19 | 3,76 | 0,41 | PC_Ether_34 |
| 3,35 | 0,1 | 2,36 | 0,2 | PC_Ether_36 |
| 1,46 | 0,08 | 0,91 | 0,16 | PC_Ether_38 |
| 0,36 | 0,02 | 0,2 | 0,01 | PC_Ether_40 |
| 0,06 | 0 | 0,04 | 0,01 | PC_Ether_42 |
| 0,01 | 0 | 0,01 | 0 | PC_Ether_44 |
| 0 | 0 | 0,21 | 0,03 | PE_Ether_30 |
| 0 | 0 | 0,11 | 0,03 | PE_Ether_32 |
| 0,07 | 0 | 4,13 | 2,01 | PE_Ether_34 |
| 0,03 | 0 | 0,16 | 0,06 | PE_Ether_36 |
| 0,09 | 0 | 0,13 | 0,06 | PE_Ether_38 |
| 0,04 | 0 | 0 | 0,01 | PE_Ether_40 |
| 0 | 0 | 0,11 | 0,03 | PI_Ether_30 |
| 0 | 0 | 0,03 | 0 | PI_Ether_32 |
| 0 | 0 | 0,04 | 0,01 | PI_Ether_34 |
| 0,08 | 0,02 | 0,13 | 0,02 | PI_Ether_36 |
| 0,11 | 0,02 | 4,69 | 1,02 | PI_Ether_38 |
| 0,03 | 0 | 0 | 0 | PI_Ether_40 |
| 0,01 | 0 | 0,11 | 0,09 | PI_Ether_42 |
| 0 | 0 | 0 | 0 | PI_Ether_44 |
| 0 | 0 | 0,09 | 0,02 | PS_Ether_30 |
| 0 | 0 | 0,02 | 0,01 | PS_Ether_32 |
| 0,01 | 0 | 0,86 | 0,37 | PS_Ether_34 |
| 0,07 | 0,01 | 0,06 | 0,02 | PS_Ether_36 |
| 0,03 | 0 | 0,03 | 0,01 | PS_Ether_38 |
| 0,02 | 0 | 0 | 0 | PS_Ether_40 |

| Control cells | SD Control cells | PM Patches | SD PM patches | Cells for GPMV | SD cells for GPMV |
|---|---|---|---|---|---|
| 0,25 | 0,05 | 0,44 | 0,02 | 0,21 | 0,03 |
| 0,77 | 0,15 | 2,79 | 0,26 | 1,16 | 0,19 |
| 4,67 | 0,31 | 4,18 | 0,36 | 3,75 | 0,16 |
| 76,94 | 3,46 | 74,42 | 0,55 | 78,66 | 0,8 |
| 9,4 | 1,15 | 10,37 | 0,91 | 8,32 | 0,3 |
| 5,41 | 0,69 | 3,58 | 0,11 | 3,12 | 0,06 |
| 4,97 | 1,27 | 3,84 | 0,17 | 2,16 | 0,2 |
| 0,41 | 0,06 | 0,15 | 0,07 | 0,45 | 0,02 |
| 1,13 | 0,1 | 2,99 | 0,47 | 1,15 | 0 |
| 12,66 | 1,88 | 10,99 | 0,38 | 10,48 | 0,07 |
| 84 | 2,2 | 81,05 | 0,76 | 81,58 | 0,32 |
| 0,18 | 0,01 | 0,17 | 0,03 | 0,19 | 0,02 |
| 0,05 | 0,01 | 0,11 | 0,01 | 0,09 | 0,01 |
| 0,05 | 0,01 | 0,11 | 0 | 0,08 | 0,01 |
| 0,07 | 0,01 | 0,05 | 0,02 | 0,09 | 0,01 |
| 0,02 | 0,01 | 0,02 | 0,01 | 0,01 | 0 |
| 11,92 | 0,65 | 13,5 | 0,12 | 13,66 | 0,19 |
| 0,95 | 0,35 | 0,63 | 0,11 | 0,2 | 0,01 |
| 0,36 | 0,41 | 1,53 | 0,03 | 0,12 | 0 |
| 0,19 | 1 | 0,19 | 0,02 | 0,08 | 0 |
| 0,24 | 0,04 | 0,42 | 0,02 | 0,2 | 0,03 |
| 0,01 | 0 | 0,01 | 0 | 0 | 0 |
| 0 | 0 | 0,01 | 0 | 0 | 0 |
| 0,77 | 0,15 | 2,78 | 0,25 | 1,16 | 0,19 |
| 0 | 0 | 0,01 | 0 | 0,01 | 0 |
| 4,55 | 0,3 | 4,04 | 0,35 | 5,69 | 0,16 |
| 0,12 | 0,01 | 0,14 | 0,01 | 0,16 | 0,01 |
| 3,95 | 0,01 | 4,21 | 0,01 | 4,07 | 0,1 |
| 35,38 | 0,85 | 36,38 | 0,68 | 34,09 | 0,44 |
| 37,62 | 2,08 | 33 | 0,75 | 35,71 | 0,7 |
| 1,27 | 0,1 | 1,21 | 0,03 | 1,2 | 0,06 |
| 1,31 | 0,11 | 0,98 | 0,09 | 1,26 | 0,06 |
| 1,08 | 0,07 | 0,81 | 0,05 | 1,01 | 0,03 |
| 0,74 | 0,03 | 0,68 | 0,04 | 0,6 | 0,01 |
| 0 | 0 | 0 | 0 | 0 | 0 |
| 0,04 | 0,01 | 0,02 | 0,01 | 0,06 | 0,01 |
| 0,23 | 0,04 | 0,11 | 0,06 | 0,3 | 0,01 |
| 0,03 | 0,01 | 0,02 | 0,01 | 0,08 | 0,01 |
| 0 | 0 | 0 | 0 | 0,01 | 0 |
| 0 | 0 | 0 | 0 | 0 | 0 |
| 4,29 | 0,08 | 4,68 | 0,06 | 4,73 | 0,11 |
| 35,32 | 0,29 | 37,18 | 1,1 | 37 | 0,74 |
| 34,48 | 2,88 | 30,39 | 0,65 | 34,65 | 0,86 |
| 1,1 | 0,14 | 1,02 | 0,03 | 1,08 | 0,04 |
| 0,38 | 0,05 | 0,28 | 0,01 | 0,32 | 0,01 |
| 0,5 | 0,06 | 0,39 | 0,01 | 0,43 | 0,02 |
| 0,48 | 0,05 | 0,48 | 0,02 | 0,45 | 0,02 |

| | | | | | |
|---|---|---|---|---|---|
| 0,83 | 0,32 | 0,32 | 0,04 | 0,22 | 0,01 |
| 2,62 | 0,47 | 3,8 | 0,44 | 2,76 | 0,13 |
| 4,26 | 0,3 | 4,31 | 0,32 | 3,44 | 0,13 |
| 0,31 | 0,02 | 0,36 | 0,02 | 0,28 | 0,01 |
| 0,55 | 0,08 | 0,73 | 0,11 | 0,6 | 0,03 |
| 0,58 | 0,04 | 0,51 | 0,07 | 0,64 | 0,01 |
| 0,36 | 0,02 | 0,34 | 0,03 | 0,39 | 0,01 |
| 0,81 | 0,35 | 0,11 | 0,02 | 0,13 | 0,01 |
| 1,16 | 0,12 | 0,93 | 0,04 | 0,8 | 0,04 |
| 2,52 | 0,26 | 1,68 | 0,14 | 1,41 | 0,05 |
| 0,32 | 0,04 | 0,22 | 0,01 | 0,19 | 0,01 |
| 0,55 | 0,05 | 0,37 | 0,02 | 0,45 | 0,03 |
| 0,18 | 0,02 | 0,11 | 0 | 0,1 | 0,01 |
| 0,12 | 0,05 | 0,16 | 0,06 | 0,04 | 0,01 |
| 1,97 | 0,99 | 0,18 | 0 | 0,1 | 0,01 |
| 2,56 | 0,35 | 2,71 | 0,15 | 1,41 | 0,15 |
| 0,57 | 0,06 | 0,55 | 0,04 | 0,37 | 0,04 |
| 0,07 | 0,01 | 0,1 | 0,01 | 0,03 | 0 |
| 0,12 | 0,02 | 0,08 | 0 | 0,08 | 0 |
| 0,16 | 0,01 | 0,12 | 0,01 | 0,1 | 0,01 |
| 0,1 | 0,02 | 0,1 | 0,01 | 0,05 | 0 |
| 9,41 | 0,31 | 10,51 | 0,37 | 10,21 | 0,07 |
| 67,45 | 3,76 | 63,8 | 0,18 | 68,36 | 0,85 |
| 0,07 | 0,01 | 0,11 | 0,01 | 0,08 | 0 |
| 0,87 | 0,33 | 0,26 | 0,02 | 0,19 | 0,01 |
| 8,67 | 0,85 | 10,08 | 0,91 | 8,08 | 0,3 |
| 0,03 | 0 | 0,03 | 0,01 | 0,05 | 0,01 |
| 0,57 | 0,26 | 0,09 | 0,01 | 0,04 | 0 |
| 4,99 | 0,54 | 3,48 | 0,12 | 3,03 | 0,07 |
| 0,05 | 0,01 | 0,02 | 0,01 | 0,05 | 0,01 |
| 1,91 | 1,02 | 0,14 | 0,01 | 0,03 | 0 |
| 3,68 | 0,45 | 3,69 | 0,16 | 2,11 | 0,19 |
| 0,02 | 0 | 0,01 | 0 | 0,01 | 0 |
| 0 | 0 | 0 | 0 | 0 | 0 |
| 0 | 0 | 0 | 0 | 0 | 0 |
| 0,16 | 0,02 | 0,28 | 0,02 | 0,14 | 0,02 |
| 0,01 | 0 | 0,01 | 0 | 0 | 0 |
| 0 | 0 | 0 | 0 | 0 | 0 |
| 0,01 | 0 | 0,02 | 0,01 | 0,01 | 0 |
| 0,07 | 0,01 | 0,12 | 0,01 | 0,05 | 0,01 |
| 0 | 0,01 | 0 | 0 | 0 | 0 |
| 0 | 0 | 0 | 0 | 0 | 0 |
| 0 | 0 | 0 | 0 | 0 | 0 |
| 0,01 | 0 | 0,01 | 0 | 0,01 | 0 |
| 0,55 | 0,12 | 2,08 | 0,19 | 0,83 | 0,15 |
| 0,01 | 0 | 0,03 | 0,01 | 0,01 | 0 |
| 0 | 0 | 0,01 | 0 | 0,01 | 0 |
| 0,06 | 0,01 | 0,16 | 0,04 | 0,07 | 0,01 |
| 0,15 | 0,02 | 0,49 | 0,08 | 0,24 | 0,04 |
| 0 | 0 | 0 | 0 | 0 | 0 |

| | | | | | |
|---|---|---|---|---|---|
| 0,03 | 0 | 0,02 | 0 | 0,03 | 0 |
| 4,27 | 0,28 | 3,82 | 0,37 | 5,37 | 0,1 |
| 0,01 | 0 | 0,03 | 0,03 | 0,01 | 0 |
| 0 | 0 | 0,02 | 0,01 | 0 | 0 |
| 0,05 | 0 | 0,04 | 0,01 | 0,05 | 0,01 |
| 0,31 | 0,03 | 0,25 | 0,02 | 0,4 | 0,06 |
| 0,09 | 0 | 0,17 | 0,02 | 0,1 | 0,01 |
| 2,36 | 0,28 | 3,27 | 0,32 | 2,52 | 0,03 |
| 7,28 | 0,2 | 7,05 | 0,13 | 7,56 | 0,08 |
| 30,81 | 1,13 | 29,75 | 0,85 | 31,77 | 0,31 |
| 31,35 | 2,49 | 29,55 | 0,86 | 31,83 | 0,64 |
| 3,82 | 0,3 | 3,97 | 0,13 | 4,23 | 0,02 |
| 0,48 | 0,06 | 0,45 | 0,01 | 0,47 | 0 |
| 0,06 | 0,01 | 0,08 | 0,01 | 0,07 | 0 |
| 0,01 | 0 | 0,03 | 0,01 | 0,01 | 0 |
| 0 | 0 | 0 | 0 | 0 | 0 |
| 0,02 | 0,01 | 0,04 | 0,01 | 0,01 | 0 |
| 0,84 | 0,32 | 0,21 | 0,01 | 0,18 | 0,01 |
| 2,17 | 0,28 | 2,32 | 0,1 | 1,86 | 0,06 |
| 4,81 | 0,52 | 6,02 | 0,66 | 4,34 | 0,2 |
| 0,94 | 0,07 | 1 | 0,09 | 0,98 | 0,06 |
| 0,74 | 0,04 | 0,63 | 0,09 | 0,83 | 0,04 |
| 0,03 | 0 | 0,03 | 0,01 | 0,04 | 0 |
| 0 | 0 | 0 | 0 | 0,01 | 0 |
| 0 | 0 | 0 | 0 | 0 | 0 |
| 0,01 | 0 | 0,02 | 0 | 0,01 | 0 |
| 0,55 | 0,25 | 0,05 | 0,01 | 0,03 | 0 |
| 0,49 | 0,05 | 0,34 | 0,02 | 0,36 | 0,02 |
| 2,89 | 0,31 | 1,92 | 0,07 | 1,63 | 0,07 |
| 1,33 | 0,14 | 1,06 | 0,08 | 0,81 | 0,03 |
| 0,22 | 0,02 | 0,12 | 0,03 | 0,14 | 0,01 |
| 0,04 | 0,01 | 0,01 | 0,01 | 0,04 | 0,01 |
| 0,01 | 0 | 0 | 0 | 0,01 | 0 |
| 0 | 0 | 0 | 0 | 0 | 0 |
| 0,01 | 0,01 | 0,02 | 0,01 | 0 | 0 |
| 1,88 | 1,01 | 0,09 | 0,01 | 0,03 | 0 |
| 0,62 | 0,07 | 0,56 | 0,02 | 0,42 | 0,02 |
| 2,47 | 0,29 | 2,65 | 0,15 | 1,35 | 0,16 |
| 0,22 | 0,03 | 0,2 | 0 | 0,13 | 0 |
| 0,36 | 0,05 | 0,27 | 0,01 | 0,21 | 0,02 |
| 0,02 | 0 | 0,01 | 0 | 0,01 | 0 |
| 0 | 0 | 0 | 0 | 0 | 0 |
| 0,13 | 0,02 | 0,1 | 0,03 | 0,19 | 0,01 |
| 0,14 | 0,03 | 0,14 | 0,03 | 0,2 | 0,01 |
| 0,04 | 0,01 | 0,06 | 0,01 | 0,07 | 0,01 |
| 0 | 0 | 0 | 0 | 0 | 0 |
| 0 | 0 | 0 | 0 | 0 | 0 |
| 0 | 0 | 0 | 0 | 0 | 0 |
| 0,02 | 0 | 0,02 | 0 | 0,03 | 0 |
| 0,03 | 0 | 0,09 | 0,01 | 0,06 | 0,01 |

| | | | | | |
|---|---|---|---|---|---|
| 0,01 | 0 | 0 | 0 | 0,01 | 0 |
| 0,04 | 0,01 | 0,11 | 0 | 0,07 | 0,01 |
| 0,01 | 0 | 0,01 | 0 | 0 | 0 |
| 0,06 | 0,01 | 0,04 | 0,02 | 0,09 | 0,01 |
| 0,01 | 0,01 | 0,02 | 0,01 | 0 | 0 |
| 0 | 0 | 0 | 0 | 0,01 | 0 |
| 0,02 | 0 | 0,02 | 0 | 0,03 | 0 |
| 0,02 | 0 | 0,08 | 0,01 | 0,05 | 0,01 |
| 0 | 0 | 0 | 0 | 0 | 0 |
| 0,02 | 0 | 0,01 | 0 | 0,01 | 0 |
| 0,03 | 0 | 0,1 | 0,01 | 0,06 | 0,01 |
| 0 | 0 | 0 | 0 | 0 | 0 |
| 0,05 | 0,01 | 0,04 | 0,02 | 0,07 | 0,01 |
| 0,02 | 0 | 0,01 | 0 | 0,02 | 0,01 |
| 0 | 0 | 0 | 0 | 0 | 0 |
| 0 | 0 | 0 | 0 | 0 | 0 |
| 0,01 | 0,01 | 0,02 | 0,01 | 0 | 0 |
| 1,67 | 0,13 | 2,05 | 0,17 | 1,55 | 0,06 |
| 10,21 | 0,75 | 11,39 | 0,18 | 12,06 | 0,17 |
| 0,03 | 0,01 | 0,06 | 0,01 | 0,04 | 0 |
| 0,62 | 0,29 | 0,02 | 0 | 0 | 0 |
| 0,32 | 0,06 | 0,61 | 0,11 | 0,19 | 0,01 |
| 0,51 | 0,25 | 0,01 | 0,01 | 0 | 0 |
| 0,45 | 0,16 | 0,32 | 0,04 | 0,11 | 0 |
| 0,01 | 0 | 0 | 0 | 0 | 0 |
| 1,84 | 0,98 | 0,01 | 0 | 0 | 0 |
| 0,11 | 0,02 | 0,18 | 0,02 | 0,08 | 0 |
| 0 | 0 | 0 | 0 | 0 | 0 |
| 0,75 | 0,02 | 0,99 | 0,02 | 0,97 | 0,02 |
| 6,37 | 0,15 | 7,84 | 0,23 | 7,64 | 0,32 |
| 3,68 | 0,4 | 3,61 | 0,13 | 4,05 | 0,15 |
| 0,34 | 0,04 | 0,33 | 0,02 | 0,33 | 0,01 |
| 0,16 | 0,02 | 0,15 | 0,01 | 0,15 | 0,01 |
| 0,23 | 0,03 | 0,25 | 0,01 | 0,23 | 0,01 |
| 0,28 | 0,03 | 0,33 | 0,01 | 0,29 | 0,01 |
| 0,62 | 0,29 | 0,01 | 0 | 0 | 0 |
| 0,05 | 0,01 | 0,06 | 0 | 0,04 | 0 |
| 0,03 | 0 | 0,05 | 0 | 0,02 | 0 |
| 0,04 | 0 | 0,07 | 0,01 | 0,03 | 0 |
| 0,14 | 0,05 | 0,32 | 0,08 | 0,03 | 0 |
| 0,03 | 0 | 0,06 | 0,02 | 0,03 | 0 |
| 0,04 | 0 | 0,07 | 0,01 | 0,04 | 0 |
| 0,7 | 0,35 | 0 | 0 | 0 | 0 |
| 0,03 | 0 | 0,03 | 0 | 0,02 | 0 |
| 0,19 | 0,06 | 0,26 | 0,03 | 0,07 | 0 |
| 0,02 | 0 | 0,01 | 0 | 0,01 | 0 |
| 0,01 | 0 | 0,01 | 0,01 | 0,01 | 0 |
| 0,02 | 0,01 | 0,02 | 0 | 0,01 | 0 |
| 0 | 0 | 0,01 | 0 | 0 | 0 |
| 1,84 | 0,98 | 0,01 | 0 | 0,01 | 0 |

| | | | | | |
|---|---|---|---|---|---|
| 0,06 | 0,01 | 0,08 | 0,01 | 0,05 | 0 |
| 0,01 | 0 | 0,01 | 0 | 0,01 | 0 |
| 0,04 | 0,01 | 0,07 | 0,01 | 0,01 | 0 |
| 0 | 0 | 0 | 0 | 0 | 0 |
| 0 | 0 | 0 | 0 | 0 | 0 |
| 0,12 | 0 | 0,17 | 0 | 0,15 | 0,01 |
| 1,55 | 0,14 | 1,88 | 0,17 | 1,41 | 0,06 |
| 5,47 | 0,29 | 6,22 | 0,19 | 6,54 | 0,23 |
| 3,7 | 0,35 | 3,97 | 0,11 | 4,33 | 0,08 |
| 0,86 | 0,1 | 1 | 0,03 | 1,01 | 0,06 |
| 0,18 | 0,03 | 0,2 | 0,01 | 0,19 | 0 |
| 0,03 | 0,01 | 0,05 | 0,01 | 0,03 | 0 |
| 0 | 0 | 0,02 | 0 | 0,01 | 0 |
| 0,01 | 0 | 0,01 | 0 | 0 | 0 |
| 0,62 | 0,29 | 0,01 | 0 | 0 | 0 |
| 0,2 | 0,06 | 0,41 | 0,09 | 0,08 | 0 |
| 0,05 | 0 | 0,09 | 0,01 | 0,04 | 0,01 |
| 0,06 | 0 | 0,09 | 0,02 | 0,06 | 0 |
| 0,02 | 0 | 0,02 | 0,01 | 0,01 | 0 |
| 0,01 | 0 | 0,01 | 0 | 0 | 0 |
| 0,51 | 0,24 | 0 | 0 | 0 | 0 |
| 0 | 0 | 0,01 | 0 | 0 | 0 |
| 0,23 | 0,1 | 0,03 | 0,01 | 0,04 | 0 |
| 0,21 | 0,06 | 0,28 | 0,03 | 0,06 | 0 |
| 0,01 | 0 | 0 | 0 | 0,01 | 0 |
| 0,01 | 0 | 0 | 0 | 0 | 0 |
| 0 | 0 | 0 | 0 | 0 | 0 |
| 0 | 0 | 0,01 | 0 | 0 | 0 |
| 1,83 | 0,98 | 0 | 0 | 0 | 0 |
| 0,06 | 0,02 | 0,1 | 0,01 | 0,02 | 0 |
| 0,04 | 0 | 0,05 | 0,01 | 0,03 | 0 |
| 0,02 | 0 | 0,02 | 0 | 0,02 | 0 |
| 0 | 0 | 0 | 0 | 0,01 | 0 |

| GPMV | SD GPMV | LYB 10 | Control cells | SD Control cells | PM Patches |
|---|---|---|---|---|---|
| 1,93 | 0,55 | Total_CER_H2O | 0,12 | 0,01 | 0,15 |
| 2,76 | 0,78 | Total_HexCER_H2O | 0,11 | 0,01 | 0,22 |
| 2,87 | 0,34 | Total_SM | 3,78 | 0,3 | 1,9 |
| 54,71 | 4,56 | Total_PC | 73,54 | 0,69 | 74,82 |
| 15,21 | 3,51 | Total_PE | 8,55 | 0,11 | 11,27 |
| 14,71 | 4,2 | Total_PI | 7,87 | 0,66 | 5,98 |
| 7,4 | 0,99 | Total_PS | 5,43 | 0,33 | 5,37 |
| 0 | 0 | Total_CL | 0,57 | 0,03 | 0,15 |
| 6,39 | 0,35 | Total_Chol | 3,05 | 0,13 | 8,65 |
| 27,02 | 2,42 | GPL_short (30-32C) | 10,57 | 0,11 | 14,72 |
| 56,79 | 2,67 | GPL_long (34-40C) | 84,5 | 0,29 | 82,28 |
| 0,18 | 0,06 | GPL_very_long (42-44C) | 0,45 | 0,02 | 0,43 |
| 0,14 | 0,04 | PC_Lyso | 0,12 | 0 | 0,06 |
| 0,02 | 0,03 | PE_Lyso | 0,07 | 0,01 | 0,07 |
| 0,22 | 0,16 | PI_Lyso | 0,09 | 0,02 | 0,05 |
| 0,38 | 0,12 | PS_Lyso | 0 | 0 | 0,04 |
| 26,17 | 1,79 | PC_Ether | 14,3 | 0,31 | 15,98 |
| 10,54 | 2,16 | PE_Ether | 0,25 | 0,01 | 0,67 |
| 7,06 | 2,29 | PI_Ether | 0,42 | 0,04 | 0,75 |
| 2,6 | 0,37 | PS_Ether | 0,27 | 0,01 | 0,28 |
| 1,54 | 0,37 | CER_H2O | 0,11 | 0,01 | 0,14 |
| 0,27 | 0,14 | DHCER_H2O | 0,01 | 0 | 0,01 |
| 0,11 | 0,05 | CERP_H2O | 0 | 0 | 0 |
| 2,56 | 0,68 | HexCER_H2O | 0,1 | 0,01 | 0,2 |
| 0,2 | 0,11 | HexDHCER_H2O | 0 | 0 | 0,02 |
| 2,55 | 1,66 | SM | 3,56 | 0,23 | 1,86 |
| 0,31 | 0,11 | DHSM | 0,23 | 0,07 | 0,04 |
| 5,05 | 0,12 | DB_0 | 5,33 | 0,04 | 6,77 |
| 18,94 | 1,68 | DB_1 | 36,42 | 0,24 | 37,93 |
| 11,53 | 1,48 | DB_2 | 23,7 | 0,27 | 22,71 |
| 2,17 | 1,74 | DB_3 | 3,28 | 0,07 | 3,19 |
| 0,22 | 0,04 | DB_4 | 4,85 | 0,27 | 3,77 |
| 1,12 | 0,32 | DB_5 | 3,84 | 0,16 | 2,83 |
| 5,88 | 1,47 | DB_6 | 2,51 | 0,09 | 2,34 |
| 0 | 0 | CL_2_DB | 0 | 0 | 0 |
| 0 | 0 | CL_3_DB | 0,04 | 0 | 0,02 |
| 0 | 0 | CL_4_DB | 0,15 | 0,01 | 0,06 |
| 0 | 0 | CL_5_DB | 0,23 | 0,01 | 0,1 |
| 0 | 0 | CL_6_DB | 0,11 | 0 | 0,05 |
| 0 | 0 | CL_7_DB | 0,02 | 0 | 0,01 |
| 5,27 | 0,47 | PC_UI_0 | 6,15 | 0,11 | 7,97 |
| 36,86 | 2,35 | PC_UI_1 | 36,35 | 0,24 | 38,49 |
| 11,27 | 1,97 | PC_UI_2 | 22,43 | 0,45 | 21,05 |
| 0,45 | 0,05 | PC_UI_3 | 2,75 | 0,05 | 2,5 |
| 0,14 | 0,03 | PC_UI_4 | 2,26 | 0,05 | 1,84 |
| 0,23 | 0,08 | PC_UI_5 | 2,17 | 0,05 | 1,65 |
| 0,48 | 0,02 | PC_UI_6 | 1,59 | 0,04 | 1,32 |

| | | | | | |
|---|---|---|---|---|---|
| 0,57 | 0,08 | PE_UI_0 | 0,23 | 0 | 0,31 |
| 2,08 | 0,38 | PE_UI_1 | 2,03 | 0,04 | 3,36 |
| 1,49 | 0,83 | PE_UI_2 | 2,1 | 0,07 | 2,96 |
| 1,57 | 1,74 | PE_UI_3 | 0,46 | 0,01 | 0,66 |
| 8,91 | 2,05 | PE_UI_4 | 1,13 | 0,02 | 1,56 |
| 0,15 | 0,03 | PE_UI_5 | 1,42 | 0,02 | 1,34 |
| 0,45 | 0,06 | PE_UI_6 | 1,18 | 0,01 | 1,08 |
| 0,85 | 0,34 | PI_UI_0 | 0,25 | 0,01 | 0,13 |
| 1,1 | 0,22 | PI_UI_1 | 2,1 | 0,14 | 1,28 |
| 6,24 | 2,27 | PI_UI_2 | 1,88 | 0,14 | 1,71 |
| 0,94 | 0,22 | PI_UI_3 | 0,7 | 0,06 | 0,62 |
| 0,16 | 0,06 | PI_UI_4 | 2,12 | 0,26 | 1,38 |
| 1,52 | 0,26 | PI_UI_5 | 0,62 | 0,05 | 0,39 |
| 3,91 | 1,49 | PI_UI_6 | 0,19 | 0,02 | 0,47 |
| 0,72 | 0,12 | PS_UI_0 | 0,27 | 0,02 | 0,21 |
| 2,94 | 0,28 | PS_UI_1 | 3,12 | 0,17 | 3,24 |
| 0,17 | 0,06 | PS_UI_2 | 0,46 | 0,02 | 0,49 |
| 2,04 | 0,28 | PS_UI_3 | 0,15 | 0,02 | 0,22 |
| 0,04 | 0,01 | PS_UI_4 | 0,35 | 0,03 | 0,28 |
| 0,06 | 0,07 | PS_UI_5 | 0,73 | 0,06 | 0,56 |
| 1,43 | 0,49 | PS_UI_6 | 0,37 | 0,03 | 0,37 |
| 20,38 | 1,58 | PC_short | 10,2 | 0,11 | 14,11 |
| 28,19 | 4,28 | PC_long | 63,25 | 0,6 | 60,55 |
| 0,07 | 0,03 | PC_very_long | 0,21 | 0,01 | 0,16 |
| 2 | 0,29 | PE_short | 0,16 | 0 | 0,3 |
| 12,61 | 3,28 | PE_long | 8,34 | 0,11 | 10,89 |
| 0 | 0 | PE_very_long | 0,06 | 0 | 0,08 |
| 1,6 | 0,55 | PI_short | 0,12 | 0,01 | 0,16 |
| 12,53 | 3,81 | PI_long | 7,67 | 0,65 | 5,7 |
| 0,11 | 0,08 | PI_very_long | 0,09 | 0 | 0,13 |
| 3,04 | 0,55 | PS_short | 0,09 | 0,01 | 0,16 |
| 3,46 | 0,46 | PS_long | 5,24 | 0,32 | 5,14 |
| 0 | 0 | PS_very_long | 0,1 | 0,01 | 0,07 |
| 0,06 | 0,03 | CER_H2O_10 | 0 | 0 | 0 |
| 0,08 | 0,05 | CER_H2O_14 | 0 | 0 | 0 |
| 0,5 | 0,18 | CER_H2O_16 | 0,05 | 0,01 | 0,07 |
| 0,25 | 0,08 | CER_H2O_18 | 0 | 0 | 0,01 |
| 0,1 | 0,03 | CER_H2O_20 | 0 | 0 | 0 |
| 0,14 | 0,08 | CER_H2O_22 | 0,03 | 0 | 0,01 |
| 0,54 | 0,12 | CER_H2O_24 | 0,04 | 0 | 0,05 |
| 0,25 | 0,02 | CER_H2O_26 | 0 | 0 | 0,01 |
| 0,13 | 0,06 | HexCER_H2O_10 | 0 | 0 | 0 |
| 0,05 | 0,04 | HexCER_H2O_12 | 0 | 0 | 0 |
| 0,13 | 0,12 | HexCER_H2O_14 | 0 | 0 | 0 |
| 1,57 | 0,1 | HexCER_H2O_16 | 0,07 | 0 | 0,13 |
| 0,15 | 0,1 | HexCER_H2O_18 | 0 | 0 | 0 |
| 0,07 | 0,07 | HexCER_H2O_20 | 0 | 0 | 0 |
| 0,3 | 0,13 | HexCER_H2O_22 | 0,01 | 0 | 0,01 |
| 0,36 | 0,19 | HexCER_H2O_24 | 0,03 | 0 | 0,06 |
| 0 | 0 | HexCER_H2O_26 | 0 | 0 | 0 |

| | | | | | |
|---|---|---|---|---|---|
| 0,03 | 0,01 | SM_14 | 0,05 | 0 | 0,02 |
| 1,93 | 0,86 | SM_16 | 2,89 | 0,1 | 1,72 |
| 0,33 | 0,47 | SM_18 | 0,08 | 0,04 | 0,02 |
| 0,19 | 0,27 | SM_20 | 0,07 | 0,04 | 0,01 |
| 0,23 | 0,06 | SM_22 | 0,31 | 0,14 | 0,02 |
| 0,15 | 0,1 | SM_24 | 0,38 | 0,05 | 0,11 |
| 0,52 | 0,07 | PC_28 | 0,12 | 0 | 0,26 |
| 2,64 | 0,21 | PC_30 | 1,44 | 0,01 | 3,42 |
| 23,23 | 1,77 | PC_32 | 8,57 | 0,1 | 10,41 |
| 14,4 | 2,54 | PC_34 | 31,85 | 0,11 | 31,19 |
| 11,54 | 1,43 | PC_36 | 24,5 | 0,57 | 23,09 |
| 1,94 | 0,31 | PC_38 | 5,62 | 0,08 | 5,28 |
| 0,23 | 0,03 | PC_40 | 1,24 | 0,04 | 0,95 |
| 0,03 | 0,01 | PC_42 | 0,17 | 0,01 | 0,13 |
| 0,04 | 0,02 | PC_44 | 0,04 | 0 | 0,03 |
| 0 | 0 | PE_28 | 0 | 0 | 0,01 |
| 0,8 | 0,07 | PE_30 | 0,01 | 0 | 0,06 |
| 1,8 | 0,23 | PE_32 | 0,14 | 0 | 0,21 |
| 9,85 | 2,16 | PE_34 | 1,24 | 0,04 | 2,09 |
| 2,57 | 2,52 | PE_36 | 3,24 | 0,11 | 4,84 |
| 0,17 | 0,01 | PE_38 | 1,8 | 0,04 | 2,03 |
| 0 | 0 | PE_40 | 2 | 0,05 | 1,89 |
| 0 | 0 | PE_42 | 0,05 | 0 | 0,06 |
| 0 | 0 | PE_44 | 0,01 | 0 | 0,01 |
| 0,01 | 0,01 | PI_28 | 0 | 0 | 0 |
| 0,6 | 0,2 | PI_30 | 0,02 | 0 | 0,05 |
| 1,33 | 0,33 | PI_32 | 0,09 | 0 | 0,09 |
| 0,21 | 0,06 | PI_34 | 1,17 | 0,1 | 0,57 |
| 4,8 | 1,44 | PI_36 | 2,91 | 0,2 | 2,15 |
| 6,7 | 2,27 | PI_38 | 2,86 | 0,31 | 2,42 |
| 0,73 | 0,18 | PI_40 | 0,64 | 0,06 | 0,53 |
| 0,04 | 0,05 | PI_42 | 0,06 | 0 | 0,07 |
| 0,07 | 0,04 | PI_44 | 0,02 | 0 | 0,06 |
| 0,01 | 0,01 | PS_28 | 0 | 0 | 0 |
| 0,77 | 0,16 | PS_30 | 0 | 0 | 0,03 |
| 2,79 | 0,38 | PS_32 | 0,08 | 0,01 | 0,09 |
| 2,42 | 0,32 | PS_34 | 0,51 | 0,03 | 0,51 |
| 0,66 | 0,2 | PS_36 | 3,03 | 0,18 | 3,13 |
| 0,13 | 0,06 | PS_38 | 0,37 | 0,03 | 0,36 |
| 0,24 | 0,2 | PS_40 | 1,38 | 0,11 | 1,15 |
| 0 | 0 | PS_42 | 0,1 | 0,01 | 0,07 |
| 0 | 0 | PS_44 | 0 | 0 | 0 |
| 0 | 0 | CL_68 | 0,07 | 0,01 | 0,02 |
| 0 | 0 | CL_70 | 0,24 | 0,02 | 0,11 |
| 0 | 0 | CL_72 | 0,23 | 0,01 | 0,12 |
| 0 | 0 | CL_74 | 0,02 | 0 | 0 |
| 0 | 0 | CL_76 | 0 | 0 | 0,01 |
| 0 | 0 | CL_78 | 0 | 0 | 0 |
| 0,05 | 0,01 | PC_Lyso_short | 0,06 | 0 | 0,02 |
| 0,09 | 0,03 | PC_Lyso_long | 0,06 | 0 | 0,04 |

| | | | | | |
|---|---|---|---|---|---|
| 0 | 0 | PE_Lyso_short | 0,01 | 0 | 0,01 |
| 0,02 | 0,03 | PE_Lyso_long | 0,06 | 0,01 | 0,05 |
| 0,14 | 0,06 | PI_Lyso_short | 0,01 | 0 | 0,02 |
| 0,09 | 0,11 | PI_Lyso_long | 0,08 | 0,01 | 0,03 |
| 0,38 | 0,12 | PS_Lyso_short | 0 | 0 | 0,04 |
| 0 | 0 | PS_Lyso_long | 0 | 0 | 0 |
| 0,06 | 0,04 | PC_Lyso_Ul_0 | 0,08 | 0 | 0,03 |
| 0,02 | 0 | PC_Lyso_Ul_1 | 0,03 | 0 | 0,02 |
| 0,06 | 0,01 | PC_Lyso_Ul_2 | 0 | 0 | 0 |
| 0 | 0 | PE_Lyso_Ul_0 | 0,04 | 0 | 0,01 |
| 0,02 | 0,03 | PE_Lyso_Ul_1 | 0,03 | 0 | 0,04 |
| 0 | 0 | PE_Lyso_Ul_2 | 0 | 0 | 0,01 |
| 0,11 | 0,15 | PI_Lyso_Ul_0 | 0,07 | 0,01 | 0,03 |
| 0,11 | 0,02 | PI_Lyso_Ul_1 | 0,02 | 0 | 0,02 |
| 0 | 0 | PI_Lyso_Ul_2 | 0 | 0 | 0 |
| 0 | 0 | PS_Lyso_Ul_0 | 0 | 0 | 0 |
| 0,38 | 0,12 | PS_Lyso_Ul_1 | 0 | 0 | 0,04 |
| 20,46 | 1,96 | PC_Ether_short | 1,4 | 0,06 | 2,83 |
| 5,69 | 0,77 | PC_Ether_long | 12,82 | 0,25 | 13,09 |
| 0,02 | 0,01 | PC_Ether_very_long | 0,09 | 0 | 0,07 |
| 0,68 | 0,13 | PE_Ether_short | 0,02 | 0 | 0,03 |
| 9,86 | 2,2 | PE_Ether_long | 0,22 | 0,01 | 0,63 |
| 0,27 | 0,02 | PI_Ether_short | 0,03 | 0 | 0,02 |
| 6,75 | 2,26 | PI_Ether_long | 0,39 | 0,03 | 0,71 |
| 0,04 | 0,04 | PI_Ether_very_long | 0,01 | 0 | 0,02 |
| 0,31 | 0,1 | PS_Ether_short | 0,07 | 0,01 | 0,01 |
| 2,29 | 0,33 | PS_Ether_long | 0,2 | 0 | 0,26 |
| 0 | 0 | PS_Ether_very_long | 0,01 | 0 | 0 |
| 1,61 | 0,16 | PC_Ether_Ul_0 | 1,24 | 0,06 | 1,73 |
| 22,65 | 1,79 | PC_Ether_Ul_1 | 6,74 | 0,15 | 8,03 |
| 1,41 | 0,26 | PC_Ether_Ul_2 | 3,01 | 0,1 | 2,86 |
| 0,11 | 0,03 | PC_Ether_Ul_3 | 0,69 | 0,02 | 0,67 |
| 0,05 | 0,01 | PC_Ether_Ul_4 | 0,89 | 0,01 | 0,93 |
| 0,09 | 0,02 | PC_Ether_Ul_5 | 0,99 | 0,01 | 0,99 |
| 0,25 | 0 | PC_Ether_Ul_6 | 0,74 | 0,01 | 0,78 |
| 0,27 | 0,01 | PE_Ether_Ul_0 | 0,03 | 0 | 0,02 |
| 0,32 | 0,03 | PE_Ether_Ul_1 | 0,03 | 0 | 0,07 |
| 0,31 | 0,14 | PE_Ether_Ul_2 | 0,02 | 0 | 0,04 |
| 0,58 | 0,25 | PE_Ether_Ul_3 | 0,01 | 0 | 0,05 |
| 8,91 | 2,05 | PE_Ether_Ul_4 | 0,04 | 0 | 0,33 |
| 0,15 | 0,03 | PE_Ether_Ul_5 | 0,04 | 0 | 0,07 |
| 0 | 0 | PE_Ether_Ul_6 | 0,07 | 0 | 0,09 |
| 0,13 | 0,06 | PI_Ether_Ul_0 | 0,03 | 0 | 0,01 |
| 0,27 | 0,09 | PI_Ether_Ul_1 | 0,09 | 0,01 | 0,09 |
| 5,77 | 2,26 | PI_Ether_Ul_2 | 0,14 | 0,01 | 0,55 |
| 0,1 | 0,03 | PI_Ether_Ul_3 | 0,04 | 0,01 | 0,03 |
| 0,08 | 0,04 | PI_Ether_Ul_4 | 0,06 | 0 | 0,02 |
| 0,61 | 0,05 | PI_Ether_Ul_5 | 0,05 | 0,01 | 0,04 |
| 0,1 | 0,03 | PI_Ether_Ul_6 | 0 | 0 | 0,01 |
| 0,19 | 0,08 | PS_Ether_Ul_0 | 0,08 | 0,01 | 0,01 |

| | | | | | |
|---|---|---|---|---|---|
| 0,26 | 0,06 | PS_Ether_UI_1 | 0,13 | 0 | 0,15 |
| 0,08 | 0,03 | PS_Ether_UI_2 | 0,03 | 0 | 0,03 |
| 2,03 | 0,28 | PS_Ether_UI_3 | 0 | 0 | 0,07 |
| 0 | 0 | PS_Ether_UI_5 | 0,02 | 0 | 0,01 |
| 0,04 | 0,03 | PS_Ether_UI_6 | 0,01 | 0 | 0,01 |
| 0,29 | 0,06 | PC_Ether_30 | 0,16 | 0 | 0,34 |
| 20,17 | 1,91 | PC_Ether_32 | 1,24 | 0,05 | 2,49 |
| 3,33 | 0,4 | PC_Ether_34 | 5,8 | 0,11 | 6,18 |
| 1,87 | 0,29 | PC_Ether_36 | 4,38 | 0,16 | 4,19 |
| 0,44 | 0,1 | PC_Ether_38 | 2,12 | 0,01 | 2,25 |
| 0,06 | 0,02 | PC_Ether_40 | 0,53 | 0,02 | 0,46 |
| 0,02 | 0,01 | PC_Ether_42 | 0,07 | 0 | 0,06 |
| 0 | 0 | PC_Ether_44 | 0,02 | 0 | 0,01 |
| 0,37 | 0,01 | PE_Ether_30 | 0 | 0 | 0,02 |
| 0,31 | 0,14 | PE_Ether_32 | 0,02 | 0 | 0,01 |
| 9,53 | 2,14 | PE_Ether_34 | 0,06 | 0 | 0,38 |
| 0,17 | 0,14 | PE_Ether_36 | 0,03 | 0 | 0,08 |
| 0,15 | 0,03 | PE_Ether_38 | 0,09 | 0 | 0,13 |
| 0 | 0 | PE_Ether_40 | 0,04 | 0 | 0,04 |
| 0,22 | 0,02 | PI_Ether_30 | 0 | 0 | 0,02 |
| 0,06 | 0,02 | PI_Ether_32 | 0,02 | 0 | 0 |
| 0,1 | 0,03 | PI_Ether_34 | 0,01 | 0 | 0,01 |
| 0,13 | 0,04 | PI_Ether_36 | 0,17 | 0,01 | 0,09 |
| 6,52 | 2,25 | PI_Ether_38 | 0,17 | 0,02 | 0,6 |
| 0 | 0 | PI_Ether_40 | 0,04 | 0 | 0,01 |
| 0,04 | 0,04 | PI_Ether_42 | 0,01 | 0 | 0,02 |
| 0 | 0 | PI_Ether_44 | 0 | 0 | 0 |
| 0,23 | 0,11 | PS_Ether_30 | 0 | 0 | 0,01 |
| 0,08 | 0,03 | PS_Ether_32 | 0,07 | 0,01 | 0 |
| 2,26 | 0,32 | PS_Ether_34 | 0,02 | 0 | 0,1 |
| 0,01 | 0,01 | PS_Ether_36 | 0,11 | 0 | 0,1 |
| 0,01 | 0,01 | PS_Ether_38 | 0,05 | 0,01 | 0,04 |
| 0 | 0 | PS_Ether_40 | 0,02 | 0 | 0,02 |

| SD PM patches | Cells for GPMV | SD cells for GPMV | GPMV | SD GPMV |
|---|---|---|---|---|
| 0,02 | 0,07 | 0,01 | 1,04 | 0,64 |
| 0,06 | 0,05 | 0,01 | 0,65 | 0,24 |
| 0,22 | 1,79 | 0,32 | 1,61 | 0,34 |
| 2,35 | 77,42 | 1,79 | 63,21 | 3,53 |
| 0,99 | 9,29 | 0,26 | 10,05 | 1,87 |
| 0,87 | 6,6 | 1,19 | 17,22 | 2,12 |
| 0,48 | 4,59 | 0,41 | 6,18 | 0,64 |
| 0,06 | 0,37 | 0,04 | 0,04 | 0,02 |
| 0,9 | 3,14 | 0,13 | 30,7 | 4,46 |
| 1,21 | 12,6 | 0,28 | 30,93 | 3,53 |
| 1,44 | 84,78 | 0,64 | 62,62 | 3,91 |
| 0,06 | 0,51 | 0,07 | 0,58 | 0,09 |
| 0,01 | 0,06 | 0,02 | 0,28 | 0,04 |
| 0,05 | 0,05 | 0,01 | 0,16 | 0,11 |
| 0,01 | 0,03 | 0,01 | 0,36 | 0,16 |
| 0,01 | 0 | 0 | 0,47 | 0,09 |
| 0,06 | 14,09 | 0,09 | 23,8 | 2,41 |
| 0,14 | 0,24 | 0 | 6,3 | 1,8 |
| 0,19 | 0,3 | 0,06 | 7,46 | 1,69 |
| 0,05 | 0,15 | 0,02 | 1,56 | 0,22 |
| 0,02 | 0,07 | 0,01 | 0,9 | 0,62 |
| 0 | 0 | 0 | 0,11 | 0,03 |
| 0 | 0 | 0 | 0,04 | 0,02 |
| 0,04 | 0,05 | 0,01 | 0,47 | 0,09 |
| 0,02 | 0 | 0 | 0,18 | 0,16 |
| 0,21 | 1,62 | 0,31 | 1,48 | 0,34 |
| 0,01 | 0,14 | 0,01 | 0,14 | 0,02 |
| 0,21 | 7,16 | 0,28 | 9,1 | 0,73 |
| 0,13 | 37,31 | 1,08 | 24,82 | 3,38 |
| 0,87 | 22,6 | 0,07 | 9,42 | 1,68 |
| 0,05 | 3,68 | 0,11 | 2,59 | 0,29 |
| 0,14 | 5,41 | 0,11 | 1,38 | 0,34 |
| 0,12 | 4 | 0,23 | 2,28 | 0,15 |
| 0,23 | 2,81 | 0,15 | 6,69 | 0,87 |
| 0 | 0,01 | 0 | 0 | 0 |
| 0,01 | 0,03 | 0 | 0 | 0 |
| 0,02 | 0,08 | 0,01 | 0 | 0 |
| 0,02 | 0,14 | 0,01 | 0,01 | 0,01 |
| 0,01 | 0,1 | 0,01 | 0,03 | 0,01 |
| 0 | 0,02 | 0 | 0 | 0 |
| 0,25 | 8,12 | 0,25 | 10,78 | 0,84 |
| 0,47 | 37,64 | 0,73 | 38,92 | 0,37 |
| 1,36 | 21,41 | 0,42 | 9,28 | 1,71 |
| 0,2 | 2,99 | 0,18 | 1,32 | 0,21 |
| 0,15 | 2,78 | 0,29 | 0,98 | 0,23 |
| 0,18 | 2,53 | 0,38 | 0,85 | 0,15 |
| 0,11 | 1,93 | 0,3 | 1,07 | 0,1 |

| | | | | |
|---|---|---|---|---|
| 0,06 | 0,21 | 0,01 | 0,52 | 0,17 |
| 0,53 | 2,2 | 0,17 | 1,93 | 0,22 |
| 0,21 | 2,22 | 0,12 | 1 | 0,09 |
| 0,03 | 0,52 | 0,01 | 0,52 | 0,02 |
| 0,2 | 1,32 | 0,04 | 5,34 | 1,54 |
| 0,13 | 1,53 | 0,01 | 0,3 | 0,04 |
| 0,09 | 1,29 | 0,03 | 0,43 | 0,07 |
| 0,01 | 0,15 | 0,04 | 0,59 | 0,14 |
| 0,15 | 1,64 | 0,52 | 1,58 | 0,24 |
| 0,28 | 1,39 | 0,29 | 6,35 | 1,62 |
| 0,09 | 0,7 | 0,12 | 1,48 | 0,5 |
| 0,14 | 2,05 | 0,17 | 0,49 | 0,09 |
| 0,07 | 0,49 | 0,06 | 1,77 | 0,12 |
| 0,21 | 0,18 | 0,03 | 4,95 | 0,89 |
| 0,03 | 0,16 | 0,01 | 0,49 | 0,11 |
| 0,26 | 2,43 | 0,3 | 3,09 | 0,52 |
| 0,05 | 0,38 | 0,02 | 0,2 | 0,08 |
| 0,03 | 0,16 | 0 | 1,05 | 0,25 |
| 0,02 | 0,39 | 0,02 | 0,17 | 0,07 |
| 0,07 | 0,68 | 0,02 | 0,32 | 0,06 |
| 0,05 | 0,39 | 0,06 | 0,86 | 0,14 |
| 1 | 12,35 | 0,28 | 25,42 | 2,1 |
| 3,18 | 64,85 | 1,65 | 35,56 | 5,6 |
| 0,05 | 0,22 | 0,04 | 0,14 | 0,03 |
| 0,12 | 0,13 | 0,02 | 1,78 | 0,64 |
| 0,9 | 9,08 | 0,25 | 8,13 | 1,28 |
| 0,01 | 0,08 | 0 | 0 | 0 |
| 0,05 | 0,11 | 0,01 | 1,42 | 0,41 |
| 0,82 | 6,36 | 1,16 | 15,25 | 2,34 |
| 0,04 | 0,14 | 0,04 | 0,43 | 0,06 |
| 0,06 | 0,02 | 0 | 2,31 | 0,61 |
| 0,42 | 4,49 | 0,4 | 3,68 | 0,64 |
| 0,01 | 0,08 | 0,01 | 0,01 | 0,01 |
| 0 | 0 | 0 | 0,01 | 0 |
| 0 | 0 | 0 | 0,02 | 0,01 |
| 0,01 | 0,04 | 0,01 | 0,28 | 0,25 |
| 0 | 0 | 0 | 0,16 | 0,07 |
| 0 | 0 | 0 | 0,05 | 0,02 |
| 0 | 0 | 0 | 0,08 | 0,03 |
| 0,01 | 0,02 | 0,01 | 0,3 | 0,16 |
| 0,01 | 0 | 0 | 0,13 | 0,11 |
| 0 | 0 | 0 | 0,03 | 0,02 |
| 0 | 0 | 0 | 0,03 | 0,02 |
| 0 | 0 | 0 | 0,04 | 0,02 |
| 0,03 | 0,03 | 0,01 | 0,17 | 0,05 |
| 0 | 0 | 0 | 0,04 | 0,01 |
| 0 | 0 | 0 | 0,02 | 0,02 |
| 0 | 0 | 0 | 0,04 | 0,03 |
| 0,02 | 0,01 | 0 | 0,26 | 0,21 |
| 0 | 0 | 0 | 0,01 | 0,01 |

| | | | | |
|---|---|---|---|---|
| 0 | 0,03 | 0 | 0,03 | 0 |
| 0,17 | 1,57 | 0,3 | 1,34 | 0,32 |
| 0,02 | 0 | 0 | 0,01 | 0 |
| 0,01 | 0 | 0 | 0,01 | 0 |
| 0,01 | 0,02 | 0 | 0,12 | 0,02 |
| 0,01 | 0,16 | 0,02 | 0,1 | 0,02 |
| 0,05 | 0,2 | 0,02 | 0,53 | 0,05 |
| 0,52 | 2,54 | 0,13 | 3,91 | 0,43 |
| 0,44 | 9,58 | 0,14 | 22,98 | 2,47 |
| 0,85 | 32,7 | 0,83 | 20,24 | 3,14 |
| 1,89 | 24,28 | 0,51 | 11,55 | 1,91 |
| 0,65 | 6,37 | 0,79 | 2,98 | 0,47 |
| 0,18 | 1,46 | 0,29 | 0,59 | 0,11 |
| 0,04 | 0,18 | 0,04 | 0,1 | 0,02 |
| 0,01 | 0,03 | 0,01 | 0,05 | 0,01 |
| 0,01 | 0 | 0 | 0,05 | 0,05 |
| 0,03 | 0,01 | 0 | 0,66 | 0,18 |
| 0,06 | 0,11 | 0,01 | 1,11 | 0,34 |
| 0,29 | 1,24 | 0,11 | 5,88 | 1,67 |
| 0,4 | 3,48 | 0,18 | 1,57 | 0,31 |
| 0,2 | 1,97 | 0,04 | 0,36 | 0,1 |
| 0,13 | 2,35 | 0,01 | 0,27 | 0,05 |
| 0,01 | 0,07 | 0 | 0 | 0 |
| 0 | 0,01 | 0 | 0 | 0 |
| 0 | 0 | 0 | 0,09 | 0,11 |
| 0,02 | 0,03 | 0 | 0,46 | 0,1 |
| 0,02 | 0,07 | 0 | 0,73 | 0,22 |
| 0,1 | 0,79 | 0,23 | 0,3 | 0,1 |
| 0,25 | 2,2 | 0,56 | 5,77 | 0,83 |
| 0,4 | 2,73 | 0,29 | 7,22 | 1,59 |
| 0,14 | 0,61 | 0,1 | 1,86 | 0,12 |
| 0,02 | 0,09 | 0,03 | 0,24 | 0,02 |
| 0,02 | 0,04 | 0,01 | 0,19 | 0,06 |
| 0 | 0 | 0 | 0,01 | 0,01 |
| 0,01 | 0 | 0 | 0,6 | 0,15 |
| 0,03 | 0,01 | 0 | 1,43 | 0,47 |
| 0,04 | 0,35 | 0,08 | 1,39 | 0,17 |
| 0,23 | 2,4 | 0,22 | 1,46 | 0,54 |
| 0,04 | 0,34 | 0,01 | 0,2 | 0,06 |
| 0,14 | 1,4 | 0,11 | 0,62 | 0,19 |
| 0,01 | 0,07 | 0,01 | 0,01 | 0,01 |
| 0 | 0 | 0 | 0 | 0 |
| 0 | 0,03 | 0,01 | 0 | 0 |
| 0,03 | 0,15 | 0,02 | 0 | 0 |
| 0,03 | 0,17 | 0,02 | 0 | 0 |
| 0 | 0,01 | 0 | 0,04 | 0,01 |
| 0,01 | 0 | 0 | 0 | 0 |
| 0 | 0 | 0 | 0 | 0 |
| 0 | 0,03 | 0,01 | 0,08 | 0,01 |
| 0,01 | 0,04 | 0,01 | 0,2 | 0,03 |

| | | | | |
|---|---|---|---|---|
| 0,02 | 0,01 | 0 | 0,11 | 0,08 |
| 0,03 | 0,05 | 0,01 | 0,05 | 0,03 |
| 0 | 0 | 0 | 0,25 | 0,12 |
| 0,01 | 0,03 | 0,01 | 0,1 | 0,04 |
| 0,01 | 0 | 0 | 0,46 | 0,1 |
| 0 | 0 | 0 | 0,02 | 0,02 |
| 0 | 0,04 | 0,01 | 0,18 | 0,02 |
| 0,01 | 0,03 | 0,01 | 0,04 | 0,01 |
| 0 | 0 | 0 | 0,06 | 0,01 |
| 0,01 | 0,02 | 0,01 | 0 | 0,01 |
| 0,03 | 0,03 | 0,01 | 0,14 | 0,08 |
| 0,02 | 0 | 0 | 0,02 | 0,03 |
| 0,01 | 0,02 | 0,01 | 0,07 | 0,03 |
| 0,01 | 0,01 | 0 | 0,25 | 0,13 |
| 0 | 0 | 0 | 0,03 | 0,01 |
| 0 | 0 | 0 | 0,01 | 0,02 |
| 0,01 | 0 | 0 | 0,46 | 0,1 |
| 0,36 | 1,59 | 0,05 | 16,5 | 3,65 |
| 0,4 | 12,4 | 0,14 | 7,24 | 1,22 |
| 0,02 | 0,1 | 0,02 | 0,06 | 0,01 |
| 0,02 | 0 | 0 | 0,52 | 0,17 |
| 0,15 | 0,24 | 0 | 5,78 | 1,69 |
| 0,01 | 0,01 | 0 | 0,25 | 0,06 |
| 0,19 | 0,27 | 0,06 | 6,98 | 1,75 |
| 0,01 | 0,01 | 0 | 0,22 | 0,04 |
| 0,01 | 0 | 0 | 0,25 | 0,04 |
| 0,04 | 0,14 | 0,02 | 1,31 | 0,22 |
| 0 | 0,01 | 0 | 0 | 0 |
| 0,08 | 1,39 | 0,07 | 2,48 | 0,11 |
| 0,35 | 6,35 | 0,15 | 18,65 | 2,96 |
| 0,19 | 2,65 | 0,05 | 1,23 | 0,25 |
| 0,07 | 0,65 | 0,01 | 0,27 | 0,07 |
| 0,08 | 1,04 | 0,04 | 0,34 | 0,11 |
| 0,09 | 1,14 | 0,1 | 0,35 | 0,06 |
| 0,06 | 0,88 | 0,1 | 0,47 | 0,06 |
| 0,01 | 0 | 0 | 0,24 | 0,06 |
| 0,02 | 0,04 | 0 | 0,26 | 0,08 |
| 0,01 | 0,01 | 0 | 0,2 | 0,08 |
| 0,01 | 0,01 | 0 | 0,32 | 0,06 |
| 0,12 | 0,05 | 0 | 5,16 | 1,58 |
| 0,01 | 0,05 | 0 | 0,12 | 0,04 |
| 0,01 | 0,08 | 0 | 0,01 | 0,01 |
| 0 | 0 | 0 | 0,12 | 0,02 |
| 0,01 | 0,06 | 0,02 | 0,62 | 0,08 |
| 0,18 | 0,12 | 0,01 | 5,81 | 1,65 |
| 0,01 | 0,04 | 0,01 | 0,2 | 0,09 |
| 0,01 | 0,04 | 0,01 | 0,11 | 0,03 |
| 0,01 | 0,03 | 0,01 | 0,49 | 0,13 |
| 0 | 0,01 | 0 | 0,11 | 0,03 |
| 0,01 | 0,01 | 0 | 0,15 | 0,02 |

| | | | | |
|---|---|---|---|---|
| 0,02 | 0,09 | 0,01 | 0,3 | 0,07 |
| 0,01 | 0,02 | 0 | 0,07 | 0,03 |
| 0,03 | 0,01 | 0 | 0,99 | 0,25 |
| 0 | 0,01 | 0 | 0,01 | 0 |
| 0 | 0,01 | 0 | 0,03 | 0,01 |
| 0,05 | 0,24 | 0,01 | 0,44 | 0,02 |
| 0,32 | 1,36 | 0,06 | 16,07 | 3,65 |
| 0,25 | 5,28 | 0,26 | 3,92 | 0,51 |
| 0,26 | 4,15 | 0,07 | 2,18 | 0,45 |
| 0,21 | 2,38 | 0,17 | 0,91 | 0,21 |
| 0,09 | 0,59 | 0,09 | 0,22 | 0,05 |
| 0,02 | 0,08 | 0,02 | 0,04 | 0,01 |
| 0,01 | 0,02 | 0 | 0,02 | 0,01 |
| 0,01 | 0 | 0 | 0,33 | 0,09 |
| 0 | 0 | 0 | 0,2 | 0,08 |
| 0,12 | 0,06 | 0 | 5,42 | 1,65 |
| 0,01 | 0,03 | 0 | 0,22 | 0,05 |
| 0,02 | 0,1 | 0 | 0,13 | 0,04 |
| 0 | 0,05 | 0 | 0 | 0 |
| 0 | 0,01 | 0 | 0,22 | 0,04 |
| 0 | 0 | 0 | 0,03 | 0,03 |
| 0,01 | 0 | 0 | 0,11 | 0,04 |
| 0 | 0,09 | 0,02 | 0,23 | 0,08 |
| 0,19 | 0,14 | 0,03 | 6,64 | 1,81 |
| 0 | 0,04 | 0,01 | 0 | 0,01 |
| 0,01 | 0,01 | 0 | 0,21 | 0,03 |
| 0 | 0 | 0 | 0,01 | 0,01 |
| 0,01 | 0 | 0 | 0,19 | 0,02 |
| 0 | 0 | 0 | 0,06 | 0,03 |
| 0,03 | 0,01 | 0 | 1,18 | 0,25 |
| 0,01 | 0,07 | 0,01 | 0,09 | 0,04 |
| 0,01 | 0,04 | 0 | 0,04 | 0,01 |
| 0 | 0,02 | 0 | 0 | 0 |

| LYB 0.04 | Control cells | SD Control cells | PM Patches | SD PM patches |
|---|---|---|---|---|
| Total_CER_H2O | 0,04 | 0,01 | 0,11 | 0,02 |
| Total_HexCER_H2O | 0,06 | 0,02 | 0,14 | 0,03 |
| Total_SM | 0,48 | 0,03 | 0,5 | 0,05 |
| Total_PC | 61,18 | 8,55 | 81,35 | 2,05 |
| Total_PE | 11,2 | 1,58 | 10,07 | 1,13 |
| Total_PI | 4,88 | 2,55 | 4,47 | 0,65 |
| Total_PS | 5,9 | 2,1 | 3,09 | 0,31 |
| Total_CL | 0,52 | 0,03 | 0,16 | 0,06 |
| Total_Chol | 1,3 | 0,12 | 4,09 | 0,55 |
| GPL_short (30-32C) | 29,14 | 6,16 | 15,93 | 1,19 |
| GPL_long (34-40C) | 85,5 | 2,09 | 82,87 | 1,26 |
| GPL_very_long (42-44C) | 0,3 | 0,04 | 0,19 | 0,05 |
| PC_Lyso | 0,08 | 0,02 | 0,15 | 0,05 |
| PE_Lyso | 0,07 | 0,01 | 0,13 | 0,03 |
| PI_Lyso | 0,08 | 0,06 | 0,05 | 0,01 |
| PS_Lyso | 0,05 | 0,03 | 0,03 | 0,01 |
| PC_Ether | 15,15 | 0,72 | 17,07 | 0,66 |
| PE_Ether | 0,41 | 0,16 | 1,18 | 0,13 |
| PI_Ether | 0,42 | 0,17 | 0,64 | 0,15 |
| PS_Ether | 0,2 | 0,04 | 0,3 | 0,08 |
| CER_H2O | 0,04 | 0,01 | 0,09 | 0,02 |
| DHCER_H2O | 0,01 | 0 | 0,01 | 0 |
| CERP_H2O | 0 | 0 | 0 | 0 |
| HexCER_H2O | 0,06 | 0,02 | 0,14 | 0,03 |
| HexDHCER_H2O | 0 | 0,01 | 0,01 | 0 |
| SM | 0,47 | 0,03 | 0,48 | 0,05 |
| DHSM | 0,01 | 0 | 0,02 | 0 |
| DB_0 | 6,73 | 0,84 | 7,77 | 0,51 |
| DB_1 | 30,61 | 3,83 | 39,52 | 0,39 |
| DB_2 | 25,16 | 2,85 | 27,82 | 0,46 |
| DB_3 | 1,94 | 0,2 | 1,27 | 0,09 |
| DB_4 | 2,62 | 0,21 | 1,17 | 0,21 |
| DB_5 | 2,33 | 0,25 | 0,88 | 0,11 |
| DB_6 | 1,94 | 0,07 | 1,01 | 0,22 |
| CL_2_DB | 0 | 0 | 0 | 0 |
| CL_3_DB | 0,02 | 0 | 0,01 | 0,01 |
| CL_4_DB | 0,15 | 0,03 | 0,18 | 0,04 |
| CL_5_DB | 0,05 | 0 | 0,03 | 0,02 |
| CL_6_DB | 0,01 | 0 | 0 | 0 |
| CL_7_DB | 0 | 0 | 0 | 0 |
| PC_UI_0 | 7,14 | 0,9 | 9,2 | 0,63 |
| PC_UI_1 | 28,94 | 4,07 | 42,83 | 1,04 |
| PC_UI_2 | 20,87 | 3,32 | 26,74 | 1,18 |
| PC_UI_3 | 1,42 | 0,16 | 1,1 | 0,07 |
| PC_UI_4 | 0,77 | 0,1 | 0,39 | 0,04 |
| PC_UI_5 | 0,92 | 0,13 | 0,46 | 0,04 |
| PC_UI_6 | 0,77 | 0,1 | 0,64 | 0,04 |

| | | | | |
|---|---|---|---|---|
| PE_UI_0 | 3,58 | 1,26 | 0,38 | 0,07 |
| PE_UI_1 | 3,36 | 0,12 | 3,78 | 0,34 |
| PE_UI_2 | 4,22 | 0,21 | 3,3 | 0,43 |
| PE_UI_3 | 0,54 | 0,02 | 0,39 | 0,04 |
| PE_UI_4 | 1,54 | 0,18 | 1,29 | 0,16 |
| PE_UI_5 | 1,19 | 0,06 | 0,57 | 0,1 |
| PE_UI_6 | 0,83 | 0,1 | 0,37 | 0,05 |
| PI_UI_0 | 3,51 | 1,38 | 0,14 | 0,02 |
| PI_UI_1 | 1,01 | 0,2 | 1,17 | 0,09 |
| PI_UI_2 | 2,5 | 0,57 | 1,93 | 0,3 |
| PI_UI_3 | 0,58 | 0,14 | 0,28 | 0,04 |
| PI_UI_4 | 1,22 | 0,13 | 0,41 | 0,08 |
| PI_UI_5 | 0,56 | 0,17 | 0,19 | 0,02 |
| PI_UI_6 | 0,65 | 0,15 | 0,36 | 0,17 |
| PS_UI_0 | 10,02 | 4,07 | 0,15 | 0,02 |
| PS_UI_1 | 1,99 | 0,24 | 2,1 | 0,15 |
| PS_UI_2 | 0,35 | 0,04 | 0,33 | 0,02 |
| PS_UI_3 | 0,18 | 0,08 | 0,2 | 0,08 |
| PS_UI_4 | 0,11 | 0 | 0,06 | 0,01 |
| PS_UI_5 | 0,19 | 0,04 | 0,1 | 0,01 |
| PS_UI_6 | 0,19 | 0,04 | 0,15 | 0,05 |
| PC_short | 12,4 | 0,88 | 15,18 | 1,09 |
| PC_long | 59,53 | 1,83 | 66,04 | 2,17 |
| PC_very_long | 0,15 | 0,03 | 0,13 | 0,04 |
| PE_short | 3,62 | 1,34 | 0,36 | 0,11 |
| PE_long | 14,01 | 0,32 | 9,69 | 1,03 |
| PE_very_long | 0,06 | 0,03 | 0,02 | 0 |
| PI_short | 2,94 | 1,48 | 0,15 | 0,03 |
| PI_long | 8,43 | 1,06 | 4,29 | 0,65 |
| PI_very_long | 0,11 | 0,04 | 0,03 | 0,01 |
| PS_short | 10,18 | 4,18 | 0,23 | 0,06 |
| PS_long | 3,5 | 0,31 | 2,85 | 0,26 |
| PS_very_long | 0,02 | 0,01 | 0,01 | 0 |
| CER_H2O_10 | 0 | 0 | 0 | 0 |
| CER_H2O_14 | 0 | 0 | 0 | 0 |
| CER_H2O_16 | 0,02 | 0,02 | 0,04 | 0 |
| CER_H2O_18 | 0 | 0 | 0,01 | 0 |
| CER_H2O_20 | 0 | 0 | 0 | 0 |
| CER_H2O_22 | 0 | 0 | 0,01 | 0 |
| CER_H2O_24 | 0,01 | 0,02 | 0,03 | 0,01 |
| CER_H2O_26 | 0 | 0,01 | 0,01 | 0,01 |
| HexCER_H2O_10 | 0 | 0 | 0 | 0 |
| HexCER_H2O_12 | 0 | 0 | 0 | 0 |
| HexCER_H2O_14 | 0 | 0 | 0 | 0 |
| HexCER_H2O_16 | 0,05 | 0,02 | 0,09 | 0,02 |
| HexCER_H2O_18 | 0 | 0 | 0,01 | 0 |
| HexCER_H2O_20 | 0 | 0 | 0 | 0 |
| HexCER_H2O_22 | 0,01 | 0 | 0,01 | 0,01 |
| HexCER_H2O_24 | 0,01 | 0,01 | 0,02 | 0,01 |
| HexCER_H2O_26 | 0 | 0 | 0 | 0 |

| | | | | |
|---|---|---|---|---|
| SM_14 | 0,01 | 0 | 0,01 | 0 |
| SM_16 | 0,42 | 0,02 | 0,45 | 0,05 |
| SM_18 | 0 | 0 | 0 | 0 |
| SM_20 | 0 | 0 | 0 | 0 |
| SM_22 | 0,02 | 0 | 0,02 | 0 |
| SM_24 | 0,03 | 0 | 0,03 | 0,01 |
| PC_28 | 0,19 | 0,02 | 0,24 | 0,03 |
| PC_30 | 2,66 | 0,33 | 2,71 | 0,24 |
| PC_32 | 9,35 | 0,63 | 12,21 | 0,86 |
| PC_34 | 21,29 | 3,9 | 33,01 | 1,17 |
| PC_36 | 21,96 | 3,29 | 28,52 | 1 |
| PC_38 | 4,16 | 0,75 | 3,89 | 0,13 |
| PC_40 | 0,79 | 0,19 | 0,51 | 0,04 |
| PC_42 | 0,13 | 0,03 | 0,1 | 0,03 |
| PC_44 | 0,02 | 0 | 0,03 | 0,01 |
| PE_28 | 0 | 0 | 0,01 | 0,01 |
| PE_30 | 0,08 | 0,02 | 0,07 | 0,03 |
| PE_32 | 3,52 | 1,31 | 0,27 | 0,06 |
| PE_34 | 2,26 | 0,22 | 2,48 | 0,27 |
| PE_36 | 5,86 | 0,26 | 5,23 | 0,49 |
| PE_38 | 1,92 | 0,1 | 1,18 | 0,17 |
| PE_40 | 1,49 | 0,12 | 0,68 | 0,12 |
| PE_42 | 0,06 | 0,03 | 0,02 | 0 |
| PE_44 | 0 | 0 | 0 | 0 |
| PI_28 | 0,04 | 0,18 | 0 | 0 |
| PI_30 | 0,08 | 0,05 | 0,04 | 0 |
| PI_32 | 2,81 | 1,26 | 0,1 | 0,03 |
| PI_34 | 0,36 | 0,05 | 0,37 | 0,02 |
| PI_36 | 3,23 | 0,5 | 2,23 | 0,39 |
| PI_38 | 2,61 | 0,52 | 1,42 | 0,16 |
| PI_40 | 0,73 | 0,08 | 0,24 | 0,08 |
| PI_42 | 0,07 | 0,03 | 0,02 | 0,01 |
| PI_44 | 0,03 | 0,02 | 0,01 | 0,01 |
| PS_28 | 0 | 0 | 0 | 0 |
| PS_30 | 0,07 | 0,03 | 0,05 | 0,01 |
| PS_32 | 10,06 | 4,12 | 0,15 | 0,03 |
| PS_34 | 0,34 | 0,1 | 0,59 | 0,11 |
| PS_36 | 1,81 | 0,18 | 1,88 | 0,13 |
| PS_38 | 0,16 | 0,02 | 0,13 | 0,01 |
| PS_40 | 0,41 | 0,06 | 0,25 | 0,04 |
| PS_42 | 0,02 | 0,01 | 0,01 | 0 |
| PS_44 | 0 | 0 | 0 | 0 |
| CL_68 | 0,05 | 0,01 | 0,07 | 0,02 |
| CL_70 | 0,11 | 0,02 | 0,11 | 0,03 |
| CL_72 | 0,07 | 0 | 0,03 | 0,02 |
| CL_74 | 0 | 0 | 0 | 0 |
| CL_76 | 0 | 0 | 0 | 0 |
| CL_78 | 0 | 0,02 | 0 | 0 |
| PC_Lyso_short | 0,03 | 0,01 | 0,03 | 0,02 |
| PC_Lyso_long | 0,05 | 0,01 | 0,12 | 0,04 |

| | | | | |
|---|---|---|---|---|
| PE_Lyso_short | 0,02 | 0,01 | 0,01 | 0,01 |
| PE_Lyso_long | 0,05 | 0,01 | 0,12 | 0,03 |
| PI_Lyso_short | 0,02 | 0,02 | 0,01 | 0,01 |
| PI_Lyso_long | 0,06 | 0,04 | 0,04 | 0,01 |
| PS_Lyso_short | 0,05 | 0,03 | 0,03 | 0,01 |
| PS_Lyso_long | 0 | 0 | 0 | 0 |
| PC_Lyso_UI_0 | 0,04 | 0,02 | 0,04 | 0,03 |
| PC_Lyso_UI_1 | 0,03 | 0,01 | 0,1 | 0,02 |
| PC_Lyso_UI_2 | 0,01 | 0 | 0,01 | 0 |
| PE_Lyso_UI_0 | 0,03 | 0,01 | 0,01 | 0 |
| PE_Lyso_UI_1 | 0,05 | 0,01 | 0,13 | 0,04 |
| PE_Lyso_UI_2 | 0 | 0 | 0 | 0 |
| PI_Lyso_UI_0 | 0,06 | 0,03 | 0,03 | 0,01 |
| PI_Lyso_UI_1 | 0,03 | 0,02 | 0,01 | 0,01 |
| PI_Lyso_UI_2 | 0 | 0,01 | 0 | 0 |
| PS_Lyso_UI_0 | 0 | 0 | 0 | 0 |
| PS_Lyso_UI_1 | 0,05 | 0,03 | 0,03 | 0,01 |
| PC_Ether_short | 4,82 | 0,36 | 4,38 | 0,41 |
| PC_Ether_long | 10,25 | 0,85 | 12,62 | 0,78 |
| PC_Ether_very_long | 0,08 | 0,01 | 0,06 | 0,03 |
| PE_Ether_short | 0,33 | 1,29 | 0,05 | 0,03 |
| PE_Ether_long | 0,09 | 0,26 | 1,14 | 0,11 |
| PI_Ether_short | 0,26 | 0,13 | 0,02 | 0 |
| PI_Ether_long | 0,15 | 0,06 | 0,62 | 0,15 |
| PI_Ether_very_long | 0 | 0 | 0,01 | 0 |
| PS_Ether_short | 0,2 | 0,04 | 0,02 | 0,01 |
| PS_Ether_long | 0 | 0 | 0,28 | 0,08 |
| PS_Ether_very_long | 0 | 0 | 0 | 0 |
| PC_Ether_UI_0 | 1,5 | 0,07 | 1,97 | 0,11 |
| PC_Ether_UI_1 | 8,21 | 0,48 | 9,88 | 0,58 |
| PC_Ether_UI_2 | 3,36 | 0,23 | 3,9 | 0,3 |
| PC_Ether_UI_3 | 0,52 | 0,02 | 0,38 | 0,04 |
| PC_Ether_UI_4 | 0,39 | 0 | 0,22 | 0,01 |
| PC_Ether_UI_5 | 0,61 | 0,01 | 0,3 | 0,02 |
| PC_Ether_UI_6 | 0,63 | 0,02 | 0,41 | 0,01 |
| PE_Ether_UI_0 | 0,32 | 1,28 | 0,02 | 0,01 |
| PE_Ether_UI_1 | 0,01 | 0,02 | 0,08 | 0,02 |
| PE_Ether_UI_2 | 0 | 0,01 | 0,06 | 0,01 |
| PE_Ether_UI_3 | 0,01 | 0,02 | 0,1 | 0,01 |
| PE_Ether_UI_4 | 0,05 | 0,22 | 0,74 | 0,07 |
| PE_Ether_UI_5 | 0,01 | 0,01 | 0,1 | 0,02 |
| PE_Ether_UI_6 | 0,01 | 0,01 | 0,08 | 0,01 |
| PI_Ether_UI_0 | 0,33 | 0,13 | 0,01 | 0 |
| PI_Ether_UI_1 | 0,01 | 0 | 0,05 | 0,01 |
| PI_Ether_UI_2 | 0,06 | 0,03 | 0,5 | 0,13 |
| PI_Ether_UI_3 | 0 | 0 | 0,02 | 0,01 |
| PI_Ether_UI_4 | 0 | 0 | 0,01 | 0,01 |
| PI_Ether_UI_5 | 0,01 | 0 | 0,04 | 0,01 |
| PI_Ether_UI_6 | 0 | 0 | 0,01 | 0 |
| PS_Ether_UI_0 | 0,2 | 0,04 | 0,02 | 0,01 |

| | | | | |
|---|---|---|---|---|
| PS_Ether_UI_1 | 0 | 0 | 0,08 | 0,01 |
| PS_Ether_UI_2 | 0 | 0 | 0,02 | 0 |
| PS_Ether_UI_3 | 0 | 0 | 0,18 | 0,08 |
| PS_Ether_UI_5 | 0 | 0 | 0 | 0 |
| PS_Ether_UI_6 | 0 | 0 | 0 | 0 |
| PC_Ether_30 | 0,47 | 0,01 | 0,42 | 0,03 |
| PC_Ether_32 | 4,25 | 0,35 | 3,96 | 0,39 |
| PC_Ether_34 | 4,67 | 0,51 | 7,09 | 0,53 |
| PC_Ether_36 | 3,6 | 0,33 | 4,22 | 0,33 |
| PC_Ether_38 | 1,7 | 0,07 | 1,12 | 0,05 |
| PC_Ether_40 | 0,41 | 0,03 | 0,2 | 0,02 |
| PC_Ether_42 | 0,06 | 0 | 0,05 | 0,02 |
| PC_Ether_44 | 0,01 | 0 | 0,01 | 0,01 |
| PE_Ether_30 | 0 | 0,01 | 0,03 | 0,02 |
| PE_Ether_32 | 0,32 | 1,29 | 0,02 | 0,01 |
| PE_Ether_34 | 0,06 | 0,24 | 0,84 | 0,07 |
| PE_Ether_36 | 0,01 | 0,01 | 0,12 | 0,01 |
| PE_Ether_38 | 0,01 | 0,03 | 0,15 | 0,03 |
| PE_Ether_40 | 0 | 0,01 | 0,02 | 0 |
| PI_Ether_30 | 0 | 0 | 0,02 | 0 |
| PI_Ether_32 | 0,24 | 0,11 | 0 | 0 |
| PI_Ether_34 | 0 | 0 | 0,01 | 0 |
| PI_Ether_36 | 0,11 | 0,03 | 0,04 | 0,01 |
| PI_Ether_38 | 0,06 | 0,04 | 0,56 | 0,14 |
| PI_Ether_40 | 0 | 0 | 0 | 0 |
| PI_Ether_42 | 0 | 0 | 0,01 | 0 |
| PI_Ether_44 | 0 | 0 | 0 | 0 |
| PS_Ether_30 | 0 | 0 | 0,02 | 0,01 |
| PS_Ether_32 | 0,2 | 0,2 | 0,01 | 0 |
| PS_Ether_34 | 0 | 0 | 0,22 | 0,08 |
| PS_Ether_36 | 0 | 0 | 0,05 | 0,01 |
| PS_Ether_38 | 0 | 0 | 0,01 | 0 |
| PS_Ether_40 | 0 | 0 | 0 | 0 |

| Cells for GPMV | SD cells for GPMV | GPMV | SD GPMV |
|---|---|---|---|
| 0,05 | 0,01 | 2,29 | 0,52 |
| 0,06 | 0,01 | 1,82 | 0,62 |
| 0,41 | 0,08 | 0,43 | 0,48 |
| 82,62 | 0,92 | 53,01 | 13,23 |
| 9,99 | 0,7 | 21,21 | 12,22 |
| 4,15 | 0,19 | 14,07 | 3,58 |
| 1,96 | 0,14 | 5,77 | 1,76 |
| 0,55 | 0,06 | 0,01 | 0,01 |
| 1,3 | 0 | 6,8 | 0,28 |
| 14,88 | 0,31 | 36,21 | 4,46 |
| 83,58 | 0,35 | 60,48 | 8,15 |
| 0,27 | 0,03 | 0,26 | 0,11 |
| 0,08 | 0,02 | 0,15 | 0,03 |
| 0,1 | 0,03 | 0,05 | 0,09 |
| 0,08 | 0,01 | 0,7 | 0,51 |
| 0,01 | 0 | 0,4 | 0,29 |
| 15,2 | 0,37 | 29,2 | 2,22 |
| 0,29 | 0,01 | 18,29 | 1,25 |
| 0,19 | 0,01 | 5,88 | 1,91 |
| 0,08 | 0,01 | 1,65 | 0,53 |
| 0,04 | 0,01 | 1,88 | 0,45 |
| 0 | 0 | 0,26 | 0,14 |
| 0 | 0 | 0,15 | 0,09 |
| 0,06 | 0,01 | 1,25 | 0,97 |
| 0 | 0 | 0,57 | 0,45 |
| 0,41 | 0,08 | 0,42 | 0,39 |
| 0 | 0 | 0,01 | 0,1 |
| 7,86 | 0,18 | 9,8 | 1,24 |
| 38,21 | 0,33 | 20,45 | 5,9 |
| 30,92 | 0,32 | 8,66 | 2,68 |
| 1,53 | 0,03 | 1,36 | 0,58 |
| 1,85 | 0,08 | 0,65 | 0,73 |
| 1,46 | 0,11 | 1,2 | 0,6 |
| 0,88 | 0,07 | 4,42 | 0,65 |
| 0 | 0 | 0 | 0 |
| 0,04 | 0,01 | 0 | 0 |
| 0,37 | 0,04 | 0 | 0 |
| 0,12 | 0,01 | 0 | 0 |
| 0,02 | 0 | 0 | 0 |
| 0 | 0 | 0 | 0 |
| 9,22 | 0,22 | 8,93 | 3,16 |
| 40,79 | 0,26 | 34,31 | 6,87 |
| 29,2 | 0,33 | 8,45 | 2,85 |
| 1,33 | 0,05 | 0,37 | 0,13 |
| 0,57 | 0,04 | 0,2 | 0,13 |
| 0,79 | 0,07 | 0,22 | 0,08 |
| 0,72 | 0,02 | 0,53 | 0,2 |

| | | | |
|---|---|---|---|
| 0,3 | 0,03 | 0,41 | 0,23 |
| 3,19 | 0,31 | 1,87 | 0,33 |
| 3,87 | 0,34 | 0,94 | 0,17 |
| 0,39 | 0,03 | 0,96 | 0,58 |
| 0,87 | 0,02 | 16,13 | 11,66 |
| 0,85 | 0,05 | 0,55 | 0,55 |
| 0,52 | 0,05 | 0,33 | 0,17 |
| 0,13 | 0,01 | 1,29 | 0,72 |
| 1,05 | 0,12 | 1,57 | 0,8 |
| 1,73 | 0,13 | 4,91 | 1,54 |
| 0,29 | 0,01 | 1,22 | 0,74 |
| 0,67 | 0,04 | 0,57 | 0,64 |
| 0,21 | 0,01 | 1,52 | 0,86 |
| 0,08 | 0,01 | 2,99 | 0,95 |
| 0,1 | 0,01 | 0,83 | 0,37 |
| 1,31 | 0,12 | 2,63 | 0,86 |
| 0,26 | 0,01 | 0,16 | 0,12 |
| 0,03 | 0 | 1,07 | 0,24 |
| 0,08 | 0 | 0,02 | 0,01 |
| 0,11 | 0,01 | 0,03 | 0,02 |
| 0,07 | 0 | 1,03 | 0,42 |
| 14,55 | 0,3 | 26,8 | 3,75 |
| 67,93 | 0,99 | 28,33 | 9,5 |
| 0,14 | 0,02 | 0,03 | 0,02 |
| 0,25 | 0,02 | 2,47 | 0,81 |
| 9,68 | 0,68 | 18,93 | 12,5 |
| 0,06 | 0,01 | 0 | 0 |
| 0,06 | 0 | 3,15 | 2,01 |
| 4,05 | 0,2 | 10,93 | 1,51 |
| 0,05 | 0,01 | 0,23 | 0,18 |
| 0,02 | 0 | 3,79 | 1,22 |
| 1,92 | 0,14 | 2,28 | 0,54 |
| 0,02 | 0 | 0 | 0 |
| 0 | 0 | 0,03 | 0 |
| 0 | 0 | 0,05 | 0,02 |
| 0,03 | 0,01 | 0,44 | 0,39 |
| 0 | 0 | 0,3 | 0,29 |
| 0 | 0 | 0,1 | 0,04 |
| 0 | 0 | 0,14 | 0,11 |
| 0,01 | 0 | 0,77 | 0,67 |
| 0 | 0 | 0,46 | 0,47 |
| 0 | 0 | 0,09 | 0,03 |
| 0 | 0 | 0,1 | 0,08 |
| 0 | 0 | 0,15 | 0,1 |
| 0,04 | 0,01 | 0,37 | 0,38 |
| 0 | 0 | 0,15 | 0,14 |
| 0 | 0 | 0,11 | 0,1 |
| 0 | 0 | 0,24 | 0,17 |
| 0,02 | 0 | 0,61 | 0,5 |
| 0 | 0 | 0,01 | 0,01 |

| | | | |
|---|---|---|---|
| 0,01 | 0 | 0,01 | 0,01 |
| 0,37 | 0,08 | 0,2 | 0,18 |
| 0 | 0 | 0,04 | 0,1 |
| 0 | 0 | 0,03 | 0,08 |
| 0,01 | 0 | 0,14 | 0,1 |
| 0,02 | 0,01 | 0,02 | 0,02 |
| 0,17 | 0 | 0,48 | 0,12 |
| 2,74 | 0,17 | 2,49 | 1,15 |
| 11,61 | 0,46 | 21,62 | 2,58 |
| 33,79 | 1,42 | 16,08 | 5,9 |
| 28,76 | 0,79 | 10,31 | 2,82 |
| 4,59 | 0,27 | 1,64 | 0,65 |
| 0,73 | 0,03 | 0,21 | 0,12 |
| 0,12 | 0,01 | 0,01 | 0,01 |
| 0,02 | 0 | 0,02 | 0,02 |
| 0 | 0 | 0 | 0 |
| 0,01 | 0 | 0,77 | 0,31 |
| 0,22 | 0,02 | 1,49 | 0,51 |
| 1,96 | 0,2 | 17,14 | 11,95 |
| 5,14 | 0,41 | 1,2 | 0,21 |
| 1,43 | 0,07 | 0,55 | 0,55 |
| 1,07 | 0,07 | 0,01 | 0,01 |
| 0,05 | 0,01 | 0 | 0 |
| 0,01 | 0 | 0 | 0 |
| 0 | 0 | 0,41 | 0,55 |
| 0,01 | 0 | 0,93 | 0,7 |
| 0,04 | 0 | 1,26 | 0,59 |
| 0,48 | 0,05 | 0,44 | 0,28 |
| 1,99 | 0,18 | 4,1 | 1,22 |
| 1,2 | 0,02 | 5,17 | 1,77 |
| 0,3 | 0,02 | 0,84 | 0,3 |
| 0,04 | 0,01 | 0,1 | 0,09 |
| 0,01 | 0 | 0,13 | 0,1 |
| 0 | 0 | 0,01 | 0,02 |
| 0 | 0 | 0,63 | 0,35 |
| 0,02 | 0 | 2,5 | 0,84 |
| 0,28 | 0,01 | 1,43 | 0,38 |
| 1,26 | 0,12 | 0,72 | 0,31 |
| 0,13 | 0,01 | 0,03 | 0,02 |
| 0,24 | 0,01 | 0,04 | 0,03 |
| 0,02 | 0 | 0 | 0 |
| 0 | 0 | 0 | 0 |
| 0,2 | 0,03 | 0 | 0 |
| 0,25 | 0,01 | 0 | 0 |
| 0,09 | 0,02 | 0 | 0 |
| 0 | 0 | 0 | 0 |
| 0 | 0 | 0 | 0 |
| 0 | 0 | 0,01 | 0,01 |
| 0,03 | 0,01 | 0,06 | 0,01 |
| 0,06 | 0,01 | 0,09 | 0,02 |

| | | | |
|---|---|---|---|
| 0,01 | 0 | 0,01 | 0,03 |
| 0,09 | 0,03 | 0,03 | 0,06 |
| 0 | 0 | 0,31 | 0,2 |
| 0,07 | 0,01 | 0,39 | 0,32 |
| 0 | 0 | 0,35 | 0,21 |
| 0 | 0 | 0,06 | 0,09 |
| 0,04 | 0,01 | 0,08 | 0,02 |
| 0,04 | 0,01 | 0,02 | 0,02 |
| 0 | 0 | 0,05 | 0,01 |
| 0,02 | 0,01 | 0 | 0 |
| 0,07 | 0,02 | 0,05 | 0,09 |
| 0 | 0 | 0 | 0 |
| 0,06 | 0 | 0,35 | 0,23 |
| 0,02 | 0 | 0,27 | 0,2 |
| 0 | 0 | 0,08 | 0,09 |
| 0 | 0 | 0,05 | 0,09 |
| 0 | 0 | 0,35 | 0,22 |
| 2,64 | 0,12 | 21,62 | 0,7 |
| 12,51 | 0,43 | 7,59 | 1,1 |
| 0,06 | 0 | 0,01 | 0,01 |
| 0 | 0 | 0,55 | 0,37 |
| 0,28 | 0,01 | 17,74 | 1,27 |
| 0,01 | 0 | 0,55 | 0,46 |
| 0,17 | 0,01 | 5,26 | 1,84 |
| 0,01 | 0 | 0,06 | 0,06 |
| 0 | 0 | 0,32 | 0,27 |
| 0,07 | 0,01 | 1,33 | 0,36 |
| 0 | 0 | 0 | 0 |
| 1,75 | 0,05 | 2,76 | 0,71 |
| 7,88 | 0,3 | 24,22 | 1,2 |
| 3,99 | 0,08 | 1,55 | 0,37 |
| 0,44 | 0,02 | 0,13 | 0,03 |
| 0,28 | 0,01 | 0,07 | 0,03 |
| 0,42 | 0,02 | 0,12 | 0,04 |
| 0,44 | 0,01 | 0,36 | 0,12 |
| 0 | 0 | 0,25 | 0,15 |
| 0,04 | 0 | 0,29 | 0,07 |
| 0,03 | 0,01 | 0,18 | 0,14 |
| 0,04 | 0,01 | 0,89 | 0,47 |
| 0,05 | 0 | 16,13 | 1,17 |
| 0,06 | 0 | 0,55 | 0,55 |
| 0,07 | 0 | 0 | 0 |
| 0 | 0 | 0,27 | 0,22 |
| 0,03 | 0 | 0,43 | 0,19 |
| 0,1 | 0,01 | 4,25 | 1,72 |
| 0,02 | 0 | 0,2 | 0,18 |
| 0,01 | 0 | 0,08 | 0,04 |
| 0,02 | 0 | 0,47 | 0,3 |
| 0 | 0 | 0,17 | 0,08 |
| 0,01 | 0 | 0,21 | 0,18 |

| | | | |
|---|---|---|---|
| 0,04 | 0,01 | 0,27 | 0,16 |
| 0,01 | 0 | 0,09 | 0,09 |
| 0,01 | 0 | 1,06 | 0,24 |
| 0 | 0 | 0 | 0 |
| 0 | 0 | 0,02 | 0,02 |
| 0,39 | 0,01 | 0,61 | 0,1 |
| 2,24 | 0,12 | 21,01 | 0,68 |
| 6,68 | 0,55 | 4,61 | 1,33 |
| 4,22 | 0,16 | 2,32 | 0,4 |
| 1,31 | 0,05 | 0,6 | 0,18 |
| 0,3 | 0,02 | 0,06 | 0,05 |
| 0,05 | 0 | 0,01 | 0,01 |
| 0,01 | 0 | 0 | 0 |
| 0 | 0 | 0,37 | 0,23 |
| 0 | 0 | 0,18 | 0,14 |
| 0,09 | 0,01 | 16,93 | 1,21 |
| 0,07 | 0,01 | 0,25 | 0,13 |
| 0,1 | 0 | 0,55 | 0,55 |
| 0,02 | 0 | 0 | 0 |
| 0 | 0 | 0,51 | 0,41 |
| 0 | 0 | 0,05 | 0,05 |
| 0 | 0 | 0,16 | 0,07 |
| 0,05 | 0,01 | 0,27 | 0,28 |
| 0,1 | 0,01 | 4,82 | 1,87 |
| 0,02 | 0 | 0,01 | 0,02 |
| 0,01 | 0 | 0,06 | 0,06 |
| 0 | 0 | 0 | 0 |
| 0 | 0 | 0,23 | 0,17 |
| 0 | 0 | 0,09 | 0,09 |
| 0,02 | 0 | 1,31 | 0,34 |
| 0,03 | 0 | 0,02 | 0,02 |
| 0,02 | 0 | 0 | 0 |
| 0 | 0 | 0 | 0 |
